# Rapid learning of neural circuitry from holographic ensemble stimulation enabled by model-based compressed sensing

**DOI:** 10.1101/2022.09.14.507926

**Authors:** Marcus A. Triplett, Marta Gajowa, Benjamin Antin, Masato Sadahiro, Hillel Adesnik, Liam Paninski

## Abstract

Discovering how neural computations are implemented in the cortex at the level of monosynaptic connectivity requires probing for the existence of synapses from possibly thousands of presynaptic candidate neurons. Two-photon optogenetics has been shown to be a promising technology for mapping such monosynaptic connections via serial stimulation of neurons with single-cell resolution. However, this approach is limited in its ability to uncover connectivity at large scales because stimulating neurons one-by-one requires prohibitively long experiments. Here we developed novel computational tools that, when combined, enable learning of monosynaptic connectivity from high-speed holographic neural ensemble stimulation. First, we developed a model-based compressed sensing algorithm that identifies connections from postsynaptic responses evoked by stimulation of many neurons at once, considerably increasing the rate at which the existence and strength of synapses are screened. We show that this model-based approach, explicitly incorporating known biophysics of optogenetic mapping experiments, is critical for accurately determining synaptic connectivity using compressed sensing. Second, we developed a deep learning method that isolates the postsynaptic response evoked by each stimulus, allowing stimulation to rapidly switch between ensembles without waiting for the postsynaptic response to return to baseline. We then validated our approach by performing large-scale connectivity mapping experiments in slices from layer 2/3 of mouse primary visual cortex. Together, our system increases the throughput of monosynaptic connectivity mapping by an order of magnitude over existing approaches, enabling the acquisition of connectivity maps at speeds needed to discover the synaptic circuitry implementing neural computations.

## Introduction

The structure of synaptic connectivity is central to how the brain processes information, stores long-term memories, and mediates cognition. To uncover such connectivity, two-photon optogenetic stimulation has emerged as a promising technology due to its ability to flexibly probe neurons with single-cell resolution while monitoring postsynaptic currents (PSCs) using whole-cell recordings [Packer et al., 2012, Baker et al., 2016, Shemesh et al., 2017, Naka et al., 2019, Hage et al., 2022, Printz et al., 2023]. Yet existing optogenetic circuit mapping techniques have been limited to probing connectivity from small numbers of neurons that must be slowly stimulated one-by-one, and therefore require aggregating small-scale maps across experiments to obtain large-scale maps of connectivity [Hage et al., 2022]. By contrast, an ideal monosynaptic connectivity mapping technique would enable large numbers of synaptic connections to be identified at high speed within a single experimental session. This would provide a crucial advantage in that each experiment would produce a more comprehensive and representative map of neural circuitry, rather than having to pool together smaller experiments that each provide just a partial view of connectivity.

Multiple strategies could serve to advance optogenetic circuit mapping towards this state, although each introduces experimental and computational challenges. The simplest strategy to improve the throughput of a connectivity mapping experiment is to increase the rate at which stimulation switches between neurons. This is primarily determined by the refresh rate of the holographic spatial light modulator (SLM) if using advanced light sculpting techniques, or, in the case of laser scanning approaches, just the time required for a neuron to integrate photocurrent until it elicits an action potential [Rickgauer and Tank, 2009]. However, while this approach allows a mapping experiment to be completed in a shorter time frame, naively stimulating too quickly confounds postsynaptic measurements because the membrane conductance of the postsynaptic neuron will not have sufficient time to return to baseline conditions before the next stimulus is applied. The speed at which a mapping experiment can feasibly take place therefore becomes limited by the ability to computationally demix PSC waveforms that overlap in time.

Simulation studies predict that connectivity mapping could also be greatly accelerated through the use of compressed sensing, where, in principle, sparse connectivity could be reconstructed from few measurements provided that stimulation is applied to ensembles of randomly selected neurons at once [Hu and Chklovskii, 2009, Fletcher et al., 2011, Mishchenko and Paninski, 2012, Shababo et al., 2013, Draelos and Pearson, 2020, Navarro and Oweiss, 2023]. However, the efficacy of existing compressed sensing algorithms is fundamentally limited outside of simplified simulations because they neglect the complex biophysics intrinsic to mapping experiments. For example, failing to elicit spikes in presynaptic neurons (due to inadequate laser power or physiological stochasticity [Mardinly et al., 2018]), as well as neurotransmitter failing to be released following successful presynaptic spikes [Rosenmund et al., 1993], are both known to significantly impact the accuracy of compressed sensing [Hu and Chklovskii, 2009, Navarro and Oweiss, 2023]. Further, high rates of spontaneous synaptic currents could lead to both false-positive connections and mischaracterized synaptic weights [Shababo et al., 2013, Printz et al., 2023].

To overcome these limitations and enable high-speed, large-scale connectivity mapping experiments, we developed new computational tools for inferring connectivity from holographic stimulation of neural ensembles. While existing optogenetic circuit mapping techniques perform slow single-neuron stimulation, our tools allow experiments to proceed with rapid holographic ensemble stimulation, minimizing downtime of the holographic SLM since evoked PSCs are isolated in time, deconfounded of spontaneous PSCs, and cleaned of electrical noise by a computational demixing procedure post-hoc. To infer connectivity, we developed a model-based compressed sensing algorithm that simultaneously estimates synaptic weights, presynaptic spikes, and how such spikes depend on laser power, while accounting for biophysical constraints and any residual spontaneous PSCs not already eliminated by the demixing procedure. To validate these tools, we performed connectivity mapping experiments using the recently engineered family of fast, potent ChroME2.0 opsins [Sridharan et al., 2022] together with two-photon holographic light sculpting [Pégard et al., 2017]. We routinely mapped connectivity from hundreds of presynaptic candidates within 680×680×100 *µ*m^3^ volumes of cortical slices, together totalling more than 12,000 probed targets across experiments. By combining rapid ensemble stimulation with computational demixing and model-based compressed sensing, we reduced the stimulation time required to reconstruct connectivity by an order of magnitude over existing approaches, allowing large numbers of synapses to be quickly mapped within individual experimental sessions.

To facilitate the use of our techniques, we provide an open access software toolbox and web-based notebook implementation that researchers can straightforwardly use and validate in their own mapping experiments. While our analysis shows that the highest mapping throughput is obtained using holographic ensemble stimulation, our techniques also increase the accuracy and speed of more widely accessible mapping approaches based on conventional single-target stimulation, and therefore can enable fast and accurate connectivity mapping in a broad range of laboratories.

## Results

### Optogenetic circuit mapping framework

Accurately characterizing the presence and strength of a synaptic connection requires precise control over the initiation of presynaptic action potentials. Ideally, each stimulus would evoke just a single presynaptic spike since this provides a direct measurement of synaptic charge transfer. To this end, we combined the family of potent soma-targeted ChroME opsins [Mardinly et al., 2018, Sridharan et al., 2022] with a scanless computer-generated holography system named 3D-SHOT [Pégard et al., 2017] (Figure 1a). We first expressed ChroME2f in parvalbumin-expressing (PV) neurons by viral transfection with an adeno-associated virus [Sridharan et al., 2022]. We calibrated laser powers and illumination time such that brief (3-5 ms) periods of stimulation almost always led to either 0 or 1 action potential, with minimal instances of multiple action potentials resulting from a single pulse (Figure S1). Next, we confirmed through a similar process that we could reliably evoke action potentials when holographically stimulating ensembles of neurons at once (Figure S1, Figure S2). Finally, we combined holographic stimulation of ensembles of PV neurons with whole-cell recordings from pyramidal neurons. We focused our initial experiments on mapping connections of this kind for two reasons. First, measurements of evoked PSCs would not be affected by direct photocurrent artifacts caused by stimulating at locations near the recording electrode, as the postsynaptic pyramidal neuron is not sensitive to stimulation itself. Second, PV neurons make strong synaptic connections onto pyramidal neurons [Packer and Yuste, 2011, Hage et al., 2022], and therefore were likely to be reliably identified computationally.

**Figure 1:**
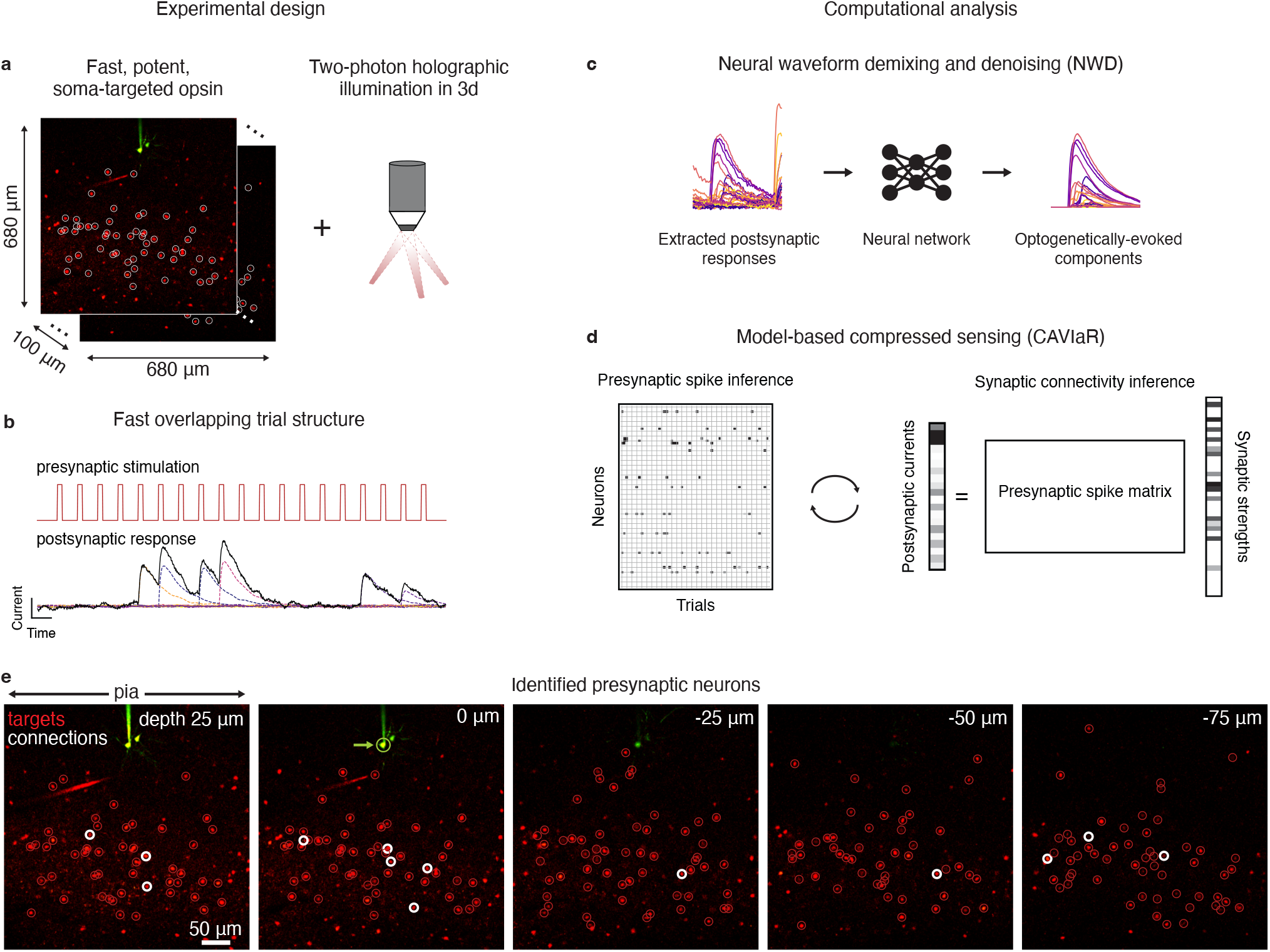
Overview of proposed high-throughput optogenetic circuit mapping framework. **a**, A fast, potent, and soma-targeted opsin (here ChroME2f) is expressed in a pool of candidate presynaptic neurons while intracellular currents are monitored from a voltage-clamped postsynaptic neuron. Two-photon holography is used to elicit spikes in ensembles of presynaptic candidates, increasing the number of putative synaptic connections tested compared to single-target stimulation. **b**, To further increase throughput, stimulation is applied at high speeds (e.g., 30−50 Hz), such that the postsynaptic membrane conductance is not guaranteed to return to baseline conditions (i.e., trials are “overlapping” in time). This allows more synapses to be tested per minute, but complicates downstream identification of their existence and strength unless the evoked PSCs are subsequently demixed. **c**, A neural network is trained to isolate the components of the postsynaptic response that are plausibly due to the optogenetic stimulus, allowing high-fidelity measurements of postsynaptic current even at very high stimulation frequencies. **d**, The “demixed” postsynaptic responses evoked by holographic ensemble stimulation are used in a model-based compressed sensing algorithm (CAVIaR) to reconstruct the underlying connectivity. Unlike traditional compressed sensing techniques, compressive mapping of synaptic circuitry requires inference of any optogenetically elicited presynaptic spikes (left) as well as the synapses themselves (right). CAVIaR alternates between inference of presynaptic spikes and synaptic connectivity, in addition to a number of other pertinent biophysical variables (see main text). **e**, Presynaptic neurons (white circles) identified via the proposed pipeline in an example experiment mapping connectivity from presynaptic PV neurons to a postsynaptic pyramidal neuron (green circle, marked by green arrow). Probed targets considered unconnected shown as red circles.

Having established the high spatiotemporal precision of our holographic stimulation technique here and in previous work [Pégard et al., 2017, Bounds et al., 2021, Sridharan et al., 2022], we then developed a computational system for processing optogenetic data and inferring synaptic connectivity. The first component of our system seeks to extract precise measurements of optogenetically-evoked PSCs. While there are many possible adverse factors that could generate variability in the membrane conductance and PSC measurements, we focused on three that we considered most critical to mapping experiments: (1) electrical noise arising from the recording electrode, which increases the variability of the observed synaptic currents and can require additional repetitions of each presynaptic stimulus to overcome; (2) spontaneous currents that, depending on their timing relative to the optogenetic stimulus, can obscure optogenetically-evoked PSCs or increase the number of false-positive connections; and (3) postsynaptic responses from preceding or subsequent trials when the interstimulus interval is shorter than the PSC decay time, which are prevalent when attempting to perform mapping experiments at very high speeds. To simultaneously eliminate these factors, we developed a deep neural network architecture for demixing and denoising PSC waveforms (Figure 1b,c) in a process that we call neural waveform demixing (NWD). The NWD network attempts to isolate optogenetically evoked PSCs stimulus-by-stimulus, such that confounding synaptic currents not driven by the optogenetic stimulus are “subtracted out” by the network and the resulting currents cleaned of electrical noise. Consequently, the NWD network allows experiments to proceed with very short interstimulus intervals since evoked PSCs from previous and subsequent trials are subtracted out and the initial baseline current reset to zero (further details given below).

The second component of our system is a model-based compressed sensing algorithm (Figure 1d). The key challenge for a compressive approach to connectivity mapping is to robustly infer presynaptic spikes from PSCs evoked by ensemble stimulation despite a multitude of unobserved sources of biophysical variability. For example, the illumination intensity required to elicit a presynaptic spike varies unpredictably between neurons because opsin is expressed differentially throughout a neural population, regardless of whether the opsin construct is delivered using a viral or transgenic targeting strategy [Bounds et al., 2021]. In a mapping experiment, this relationship between light and presynaptic spike probability cannot be observed from the presynaptic neuron itself, but must be inferred through electrophysiological measurements from the postsynaptic neuron. This inference problem is in turn complicated by the fact that PSCs can occur spontaneously, that postsynaptic responses are corrupted by electrical noise, that synaptic transmission can fail despite successful elicitation of a presynaptic spike, and that potentially tens of neurons are stimulated at a time. Accounting for the power-dependence of presynaptic spike initiation for individual neurons when only observing postsynaptic responses to ensemble stimulation is therefore computationally challenging, and constitutes only one of the inference problems that must be solved to accurately reconstruct connectivity using a compressed sensing approach.

To resolve these challenges, we embedded the compressed sensing step in a hierarchical Bayesian statistical model that captured the most critical sources of biophysical variability. Our model relates patterns of ensemble stimulation to electrophysiological measurements from the postsynaptic cell while, at the same time, accounting for variability in photoactivation and synaptic transmission across the opsin-expressing population. We developed a variational inference technique called CAVIaR (coordinate-ascent variational inference and isotonic regularization) to learn posterior distributions over the model parameters (Methods). CAVIaR identifies the existence and strength of individual synaptic connections from ensemble stimulation, substantially increasing the rate at which neural circuits can be mapped compared to single-target stimulation and increasing the accuracy compared to existing compressed sensing techniques.

### Neural waveform demixing allows mapping experiments to proceed rapidly

The NWD network has a sequential U-Net architecture [Ronneberger et al., 2015, Falk et al., 2019] that uses one dimensional convolutional filters to learn representations of the input signal at increasing levels of temporal compression. This allows the network to integrate information across the entire PSC trace, such that confounding synaptic currents preceding or following an admissible PSC “initiation zone” (typically selected to be a window of 3-12 ms following stimulation) can be accurately subtracted and the baseline current reset to zero (Figure 2a). Since the network is trained to generate noise-free, demixed PSCs from tens of thousands of simulated noisy inputs, NWD also markedly improves the overall signal quality (Figure 2). Furthermore, unlike computationally intensive algorithms for deconvolving intracellular currents [Merel et al., 2016], NWD can isolate the PSC evoked by optogenetic stimulation with just a simple forward pass through the network, with the resulting demixed PSCs usable in any connectivity inference algorithm and in real time.

**Figure 2:**
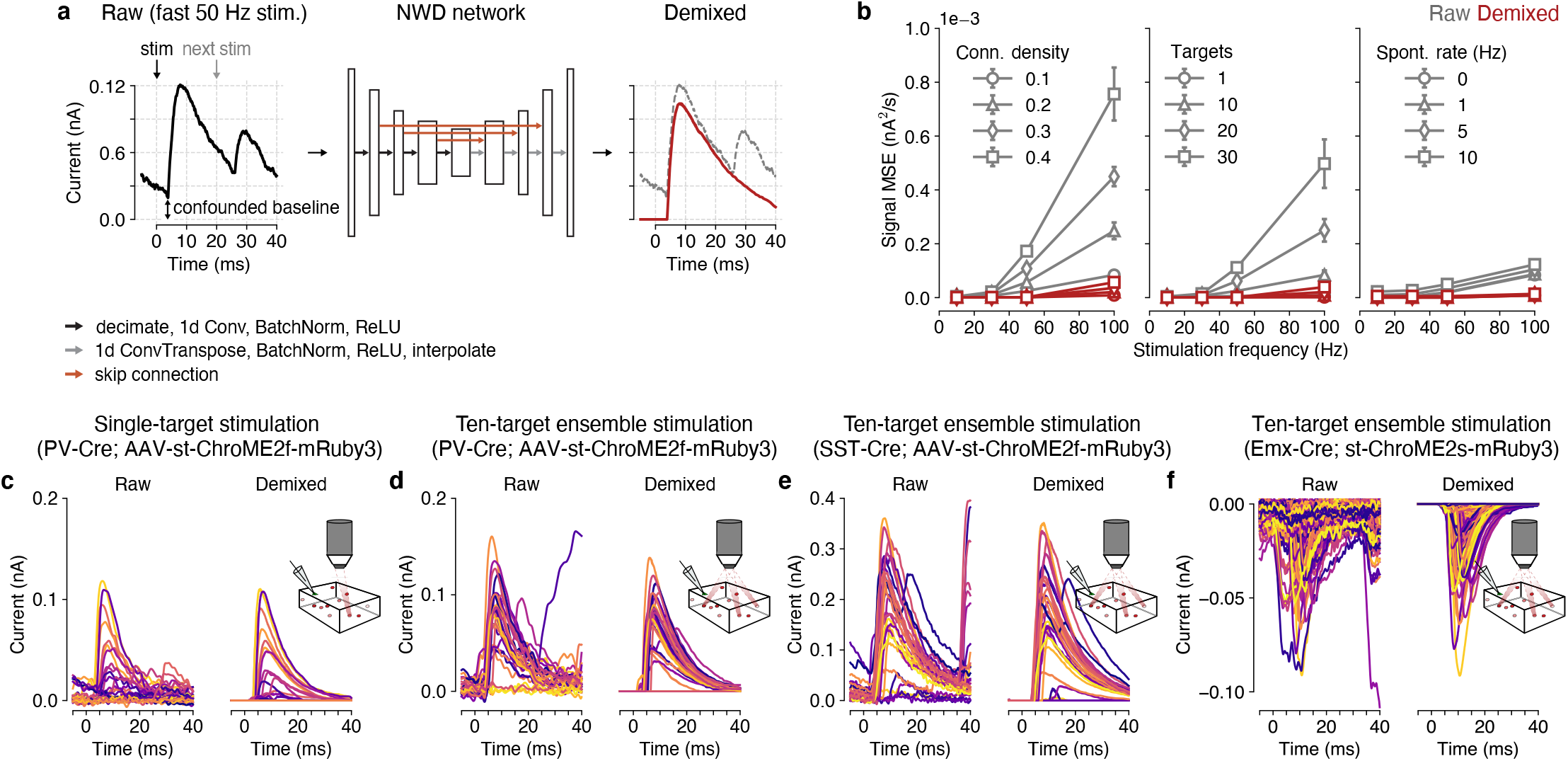
Simultaneous demixing and denoising of optogenetically evoked postsynaptic currents. **a**, Neural waveform demixing overview. Raw electrophysiological traces (left) are confounded by synaptic currents from previous and subsequent trials during fast stimulation, in addition to spontaneous synaptic currents. The NWD network (middle) is trained to isolate the PSC waveform associated with the optogenetic stimulus, resulting in a demixed trace (right, red) with the confounding synaptic currents subtracted and baseline current (at 0 ms) adjusted to 0 nA. Note that a 5 ms “pre-stimulus” window is provided to the network as additional context. **b**, Performance of NWD in simulation as a function of stimulation frequency under increasing connection density (left), number of simultaneously targeted cells (middle) and rate of spontaneous activity (right). In all cases NWD leads to a substantial improvement in signal fidelity (measured by the mean square error of the raw vs demixed trace compared to the ground truth evoked PSC). Errors are evaluated across all conditions using a single trained network. Number of simultaneously stimulated targets in left and right panels, 10. Connection density in middle and right panels, 0.1. Spontaneous PSC rate in left and middle panels, 1 Hz.**c, d**, Examples of NWD applied to optogenetically evoked inhibitory PSCs under single-target (c) and ten-target (d) holographic stimulation of PV neurons expressing ChroME2f. Panel (d) shows responses to stimulating ensembles containing the neuron in panel (c). **e**, Same as (d), but for stimulation of SST neurons. **f**, same as (c), but for stimulation of pyramidal neurons expressing ChroME2s [Sridharan et al., 2022] while recording from another pyramidal neuron.

Performing optogenetic connectivity mapping experiments at high speed poses a trade-off: the experiment can be completed much more quickly, but measurements of the postsynaptic response are much more confounded by currents from previous and subsequent trials. For this reason, a typical rate of stimulation for a connectivity mapping experiment is ∼10 Hz (i.e., stimulating a neuron approximately every 100 ms [Hage et al., 2022, Printz et al., 2023, Chen et al., 2023], much longer than the decay time of a typical PSC), which we take as a baseline rate in later method comparisons. To characterize the improvement in experiment speed-up resulting from the application of NWD, we simulated connectivity mapping experiments at full 20 kHz sampling resolution with varying synaptic connection densities and stimulation frequencies (Figure 2b, Figure S3). In a sparsely connected circuit, the primary action of NWD is to eliminate spontaneous synaptic currents and reduce electrical noise, leading to just a marginal improvement in signal fidelity compared to the raw PSC traces (Figure 2b, 0.1 connection density example). However, as synapses become more prevalent the occurrence of postsynaptic events greatly increases, with many confounding synaptic currents elicited from previous trials. Application of NWD therefore leads to a substantial improvement in the signal fidelity of the postsynaptic response at high stimulation frequencies (Figure 2b, 0.2 to 0.4 connection density examples). Similarly, the fidelity of the postsynaptic response degrades as a function of the number of simultaneously stimulated neurons and the background rate of spontaneous synaptic currents (Figure 2b, middle and right panels). However, NWD largely eliminates this effect, and thus reduces experimental constraints on the duration of interstimulus intervals since confounding synaptic currents are significantly reduced. Stimulation can therefore be increased to rates closer to the refresh rate of the SLM. In practice, this enables us to stimulate much faster than the ∼40 ms decay time typical of PSC traces in our preparations (e.g., Figure 2d).

We applied NWD to each of our experimental settings, including demixing of inhibitory PSCs from PV and somatostatin (SST) mapping experiments under both holographic single-target and ensemble stimulation (Figure 2d-e), as well as demixing of excitatory PSCs from pyramidal mapping experiments (Figure 2f). We found that we could improve the performance of the NWD network by tuning it to the time constants of either inhibitory or excitatory synaptic currents (Methods, Figure S4). In each case the NWD network led to a dramatic reduction in confounding synaptic currents and electrical noise, in agreement with our simulated results (Figure 2b).

### Ensemble stimulation can test for many synaptic connections simultaneously

We sought to capitalize on recent technological developments in opsin engineering [Baker et al., 2016, Shemesh et al., 2017, Mardinly et al., 2018, Sridharan et al., 2022], holographic light sculpting [Papagiakoumou et al., 2010, Pégard et al., 2017, Papagiakoumou et al., 2020, Adesnik and Abdeladim, 2021] and fundamental mathematical results on efficient signal reconstruction in sparse settings [Donoho, 2006, Candes and Tao, 2006, Candes et al., 2006] by developing a statistical method for inferring synaptic connectivity from holographic ensemble stimulation. This was based on the logic that pairing rapid stimulation and computational demixing with the ability to test for many potential connections with each hologram could dramatically speed up circuit mapping. We therefore created and tested a variety of connectivity inference algorithms, and ultimately found that a variational inference approach for a hierarchical Bayesian statistical model was able to overcome the limitations of existing compressed sensing approaches. Our model relates patterns of holographic stimulation to PSCs through a series of key latent variables: (1) optogenetic “power curves” that characterize the relationship between laser power and presynaptic spike probability due to variation in opsin expression and rheobase across neurons, (2) presynaptic spikes successfully elicited by photostimulation and transmitted to the postsynaptic neuron, and (3) synaptic weights that determine the amplitude of the resulting PSCs (Figure 3a). We learn posterior distributions over the latent variables using CAVIaR, an algorithm that iteratively updates the parameters of a variational approximation to the posterior such that the inferred presynaptic spikes respect the biophysical plausibility constraint of having isotonically increasing spike probabilities as a function of laser power (i.e., such that the probability of evoking a presynaptic spike increases with laser power on average [Sridharan et al., 2022]; see Methods).

**Figure 3:**
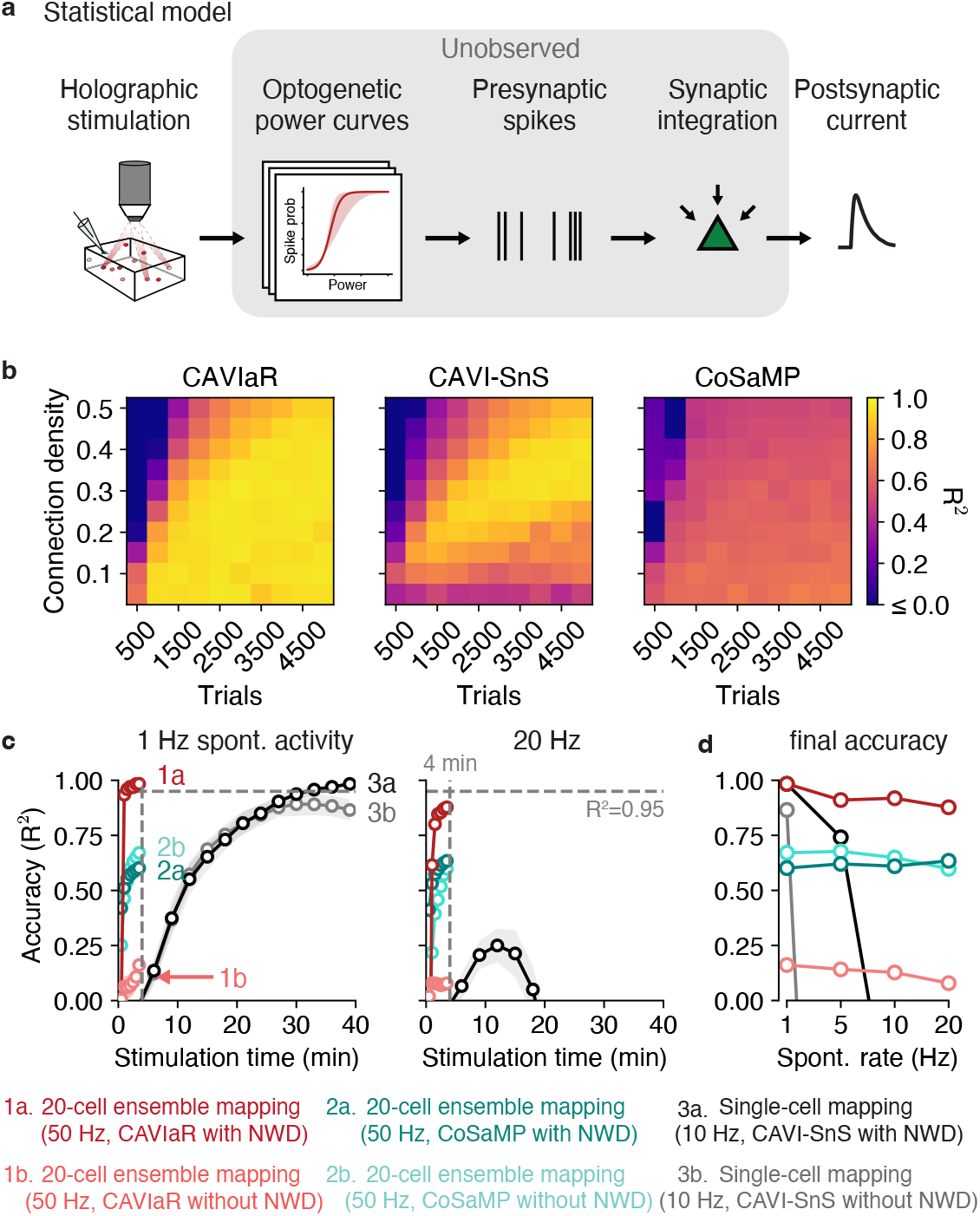
CAVIaR accurately maps synaptic connectivity at high speeds in simulation. **a**, Statistical model relating patterns of holographic stimulation to measurements of postsynaptic current via latent optogenetic power curves, presynaptic spikes, and synaptic weights. **b**, Comparison of CAVIaR (ours, left) with CAVI-SnS (middle) and CoSaMP (right) for varying connection densities as a function of increasing amounts of experimental data. Simulated presynaptic candidate population size of N=300 (hence, e.g., 4500 trials corresponds to an average of 15 stimuli per neuron). All PSCs are demixed using NWD. Note that CoSaMP requires the number of desired connections to be specified in advance; we therefore provided CoSaMP with the true number of connections, such that the reported performance represents an upper bound. R^2^ represents the coefficient of determination between the ground-truth and inferred connectivity vectors. Simulated neurons do not explicitly coincide with a specific genetically-defined cell type, but could match various cell types depending on the simulated connectivity rate. **c**, Order of magnitude improvement in mapping efficiency using NWD-accelerated connectivity mapping with ensemble stimulation. Population size, 1000. Connection density, 0.1. Left, convergence speed of CAVIaR, CoSaMP, and CAVI-SnS with and without NWD, assuming a background rate of spontaneous PSCs of 1 Hz. In this simulation, while CAVI-SnS requires more than 30 mintes of stimulation to reach an accuracy of 0.95, CAVIaR requires just 30 seconds. A high rate of spontaneous activity (20 Hz, right panel) causes fast degradation of CAVI-SnS performance due to an accumulation of false positives, but impacts CAVIaR much less. In both conditions CoSaMP does not converge to the true connectivity. Note that given additional experiment time the accuracy of CAVIaR without NWD substantially improves, but does not match CAVIaR with NWD (Figure S12). Error bars show mean *±* 1 s.d. over 20 simulations. **d**, Final accuracy of all methods as a function of spontaneous PSC rate. CAVIaR is robust against spontaneous PSCs, unlike CAVI-SnS.

A key component of our experimental design was to randomly switch between three or more laser powers while mapping. Since CAVIaR infers presynaptic spikes, we could then estimate optogenetic power curves to determine whether the plausibility constraint was satisfied. Any putative presynaptic neurons for which spike counts happen to implausibly decrease with increasing power are considered spurious, and immediately disconnected by CAVIaR. When using our software in practice, a user simply sets a parameter that imposes a minimum success rate (between 0 and 1) for evoking presynaptic spikes (given by the isotonically-constrained power curve at maximal power) and CAVIaR identifies putatively connected neurons meeting that criterion. However, PSCs can occur spontaneously, inflating the estimated presynaptic spike rate and increasing the number of false positives. We therefore simultaneously estimate the background rate of spontaneous currents and use this to adaptively adjust the plausibility criterion during inference.

We compared CAVIaR to two other approaches. (1) Standard compressed sensing. While there are a number of widely-used algorithms for compressed sensing [Rani et al., 2018], they each solve a common mathematical problem and therefore lead to closely related results. Compressive sampling matching pursuit (CoSaMP, [Needell and Tropp, 2009]) is a particularly scalable algorithm for solving the compressed sensing problem and was used in the first simulation study on compressive connectivity mapping [Hu and Chklovskii, 2009] which we therefore benchmark CAVIaR against and consider representative of the performance from related studies (e.g., ref. [Chen et al., 2023]). (2) A variational inference technique for a spike-and-slab connectivity model similar to CAVIaR but that, among other methodological differences, does not impose isotonic regularization or account for spontaneous synaptic currents (CAVI-SnS, [Shababo et al., 2013]). We evaluated the ability of each method to reconstruct connectivity in simulations where the sparsity of the underlying connectivity varied, since compressed sensing-based techniques are known to be particularly sensitive to this parameter [Donoho, 2006]. We found that CAVIaR greatly outperformed conventional compressed sensing (Figure 3b, left vs right panels), primarily because the latter does not have a model for stochastic, power-dependent presynaptic spikes. Importantly, in line with earlier results [Hu and Chklovskii, 2009], we found that algorithms failing to account for this stochasticity will yield strongly biased synaptic weight estimates since they will average over postsynaptic measurements when no presynaptic spikes occurred. CAVIaR also outperformed CAVI-SnS, which transiently demonstrated high accuracy, but ultimately generated increasingly worse estimates of connectivity with increasing amounts of data since it lacked mechanisms to prevent spontaneous PSCs from being misidentified as false positives. We found that CAVIaR consistently achieved state-of-the-art performance when repeating the analysis while varying the number of times each stimulus was repeated, the number of simultaneous targets, and the rate of background spontaneous activity as a function of the amount of experimental data collected (Figure S8, Figure S9).

Next, we estimated the speed-up obtained by combining rapid ensemble stimulation with NWD and CAVIaR, and compared this against previously established techniques. Our simulations showed that even with a large population size (in this case mapping 1000 potential presynaptic neurons), the speed of connectivity mapping was optimal when stimulating ensembles of 20 neurons at a time (Figure S9) beyond which the accuracy of presynaptic spike inference gradually degraded, though this also depended on the density of synaptic connectivity (Figure S10). Our speed-up simulations were therefore performed using 20-cell stimulation. We anticipated that the background rate of spontaneous PSCs would also be a key factor influencing the ultimate accuracy of each connectivity inference method, since a high number of spontaneous PSCs that happen to fall within the stimulation window could mimic a synapse and lead to false positives. Single-target mapping at 10 Hz with CAVI-SnS required more than 30 minutes of stimulation to cross an accuracy of 0.95 in simulations with 1000 neurons, 10% connectivity, and 1 Hz spontaneous activity (Figure 3c, left panel; accuracy measured as the *R*^2^ between the true and estimated synaptic weights). By comparison, NWD-enabled 50 Hz stimulation of 20-cell ensembles recovered synaptic connectivity at an accuracy exceeding 0.95 with just 30 seconds of stimulation using CAVIaR, resulting in a rate of connectivity inference of more than 2000 neurons per minute in low spontaneous activity conditions (exceeding an order of magnitude more neurons tested per minute compared to single-target stimulation with CAVI-SnS). In the same time period CoSaMP obtained an accuracy of less than 0.6, and in our simulations never achieved an accuracy of 0.95 regardless of stimulation time (Figure 3c,d, Figure S9). We also confirmed that similar improvements in mapping efficiency are obtained when accuracy is evaluated as a function of stimulation trials, rather than time (Figure S11).

We then increased the rate of spontaneous PSCs to 20 Hz, a more challenging regime in which the spontaneous PSC rate exceeded those in our own mapping experiments (Figure S16). This revealed several important behaviors of CAVIaR and CAVI-SnS (Figure 3c, right panel; Figure 3d). Namely, the accuracy of CAVI-SnS both with and without NWD behaved non-monotonically (similar to Figure 3b, final R^2^ at a 20 Hz spontaneous rate with and without NWD, *<*0), with connectivity estimates ultimately getting worse with additional experiment time due to the accumulation of spontaneous PSCs. However, CAVIaR with NWD obtained nearly the same accuracy when spontaneous PSCs occurred at 20 Hz as it did when they occurred at 1 Hz due to CAVIaR’s built-in mechanisms for estimating and adapting to spontaneous PSCs (final R^2^ with spontaneous rate 1 Hz, 0.98; 20 Hz, 0.88; Figure 3c,d). Similarly, CAVIaR with NWD achieved high precision and recall in simulations with a spontaneous PSC rate of 20 Hz (final precision, 1; recall, 0.8; Figure S13). These results thus indicate that our model-based compressed sensing approach should be robust in conditions with high spontaneous activity.

### Validating CAVIaR using mapping experiments in cortical slices

Having established the accuracy and speed of our connectivity mapping system in simulations, we next sought to validate our computational tools experimentally. We first wanted to confirm that connectivity maps obtained using ensemble stimulation were closely aligned to those obtained using the existing approach of single-target stimulation. We therefore performed experiments where the same population of PV neurons was mapped using randomly interleaved trials of single-target and holographic ensemble stimulation (ensembles of 10 neurons, stimulation performed at 30 Hz). We separated trials into two sets depending on whether they were obtained using single-target or ensemble stimulation, demixed the optogenetically evoked PSCs using NWD, and detected putative synapses in each set independently using CAVIaR (Figure 4a-d). For the example shown in Figure 4a, model-based compressed sensing identified the same set of synaptically connected PV neurons as with single-target stimulation and almost identical synaptic strengths (varying from 57 to 228 pA, Figure 4c,d). Across 14 different experiments we probed 2619 total presynaptic candidates (mean number of targets probed, 187; minimum, 107; maximum, 269). CAVIaR identified very similar numbers of connections when using single-target stimulation compared to ensemble stimulation (Figure 4e,f; R^2^, 0.81). Further, we confirmed that these were primarily the same set of connections when evaluated based on both their numerical synaptic strengths (Figure 4g, left; average R^2^ between connectivity vectors estimated using single-target and ensemble stimulation, 0.89), as well their binary classification (i.e., whether an ROI was connected or not, without regard to synaptic strength; Figure 4g, right; average precision, 0.95; average recall, 0.84.). Additional examples are given in Figure S14. We note, however, that connectivity mapping using single-target stimulation does not necessarily represent a perfect ground truth as false positives can still arise from spontaneous activity when using single-target stimulation (Figure 3c).

**Figure 4:**
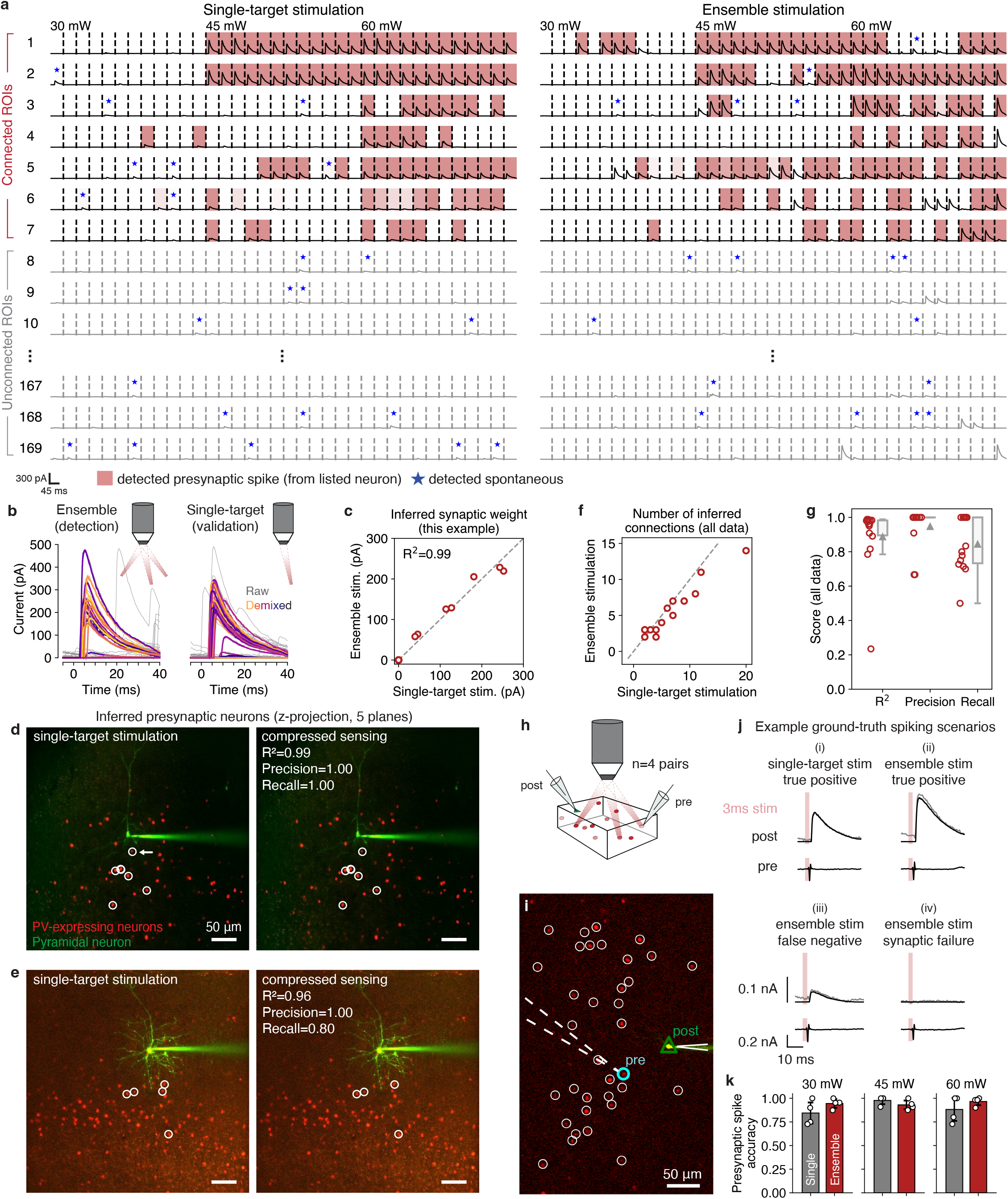
Validation of model-based compressed sensing approach for mapping PV-pyramidal connections. **a**, “Checkerboard” visualization of CAVIaR inferences. Left column shows demixed PSCs evoked by single-target stimulation of the listed ROI. Right column shows PSCs evoked by holographic stimulation of ensembles containing the listed neuron. Trials separated by dashed lines and rearranged to be in order of increasing laser power (from 30 mW to 60 mW). While all single-target stimulation trials are shown on the left, for ease of visualization only a fraction of ensemble stimulation trials are shown on the right. Shaded trials indicate when CAVIaR detects that the observed PSC was evoked by successfully spiking the listed (presynaptic) ROI; opacity of shading represents the estimated posterior uncertainty of the spike (lighter, less certain; darker, more certain). Blue stars represent when CAVIaR determines that the PSC is spontaneous (due to e.g. an uncharacteristic amplitude). Note that responses are only shaded when the neuron in the listed row actually spiked (as determined by CAVIaR). PSC waveforms that are not shaded and not labeled as spontaneous can still be attributed to other neurons that were stimulated in the same ensemble. Out of 169 stimulated ROIs, the same set of 7 were identified by CAVIaR as being presynaptic by both single-target and ensemble stimulation. **b**, Example PSCs (corresponding to neuron marked by white arrow in panel d, below) evoked by holographic stimulation of ensembles containing a putatively connected neuron (left panel). Single-target stimulation of the same neuron validates the existence of the individual synapse (right panel). Gray traces show raw PSCs, colored traces show demixed PSCs. Traces colored at random. **c**, Synaptic weights estimated by CAVIaR independently from single-target stimulation and ensemble stimulation trials show strong agreement (R^2^, 0.99). Seven synaptic connections shown; points at zero indicate no connection. **d**, Z-projection (over 5 different planes) of stimulation FOV for experiment corresponding to panels a-c. White circles show identified presynaptic neurons. Left, presynaptic neurons identified using single-target stimulation; right, presynaptic neurons identified using ensemble stimulation. NWD and CAVIaR were used in both cases. Seven connections are found using both single-target stimulation and ensemble stimulation. Note that one connection is hidden by another on a higher plane and therefore not directly visible. **e**, Additional example of similarity between synaptic connectivity maps obtained using single-target (left) vs ensemble stimulation (right) in combination with NWD and CAVIaR. **f**, Number of connections identified by CAVIaR using both single-target and ensemble stimulation across 14 experiments. Dashed gray line shows identity. **g**, Agreement between connectivity maps obtained using single-target stimulation compared to model-based compressed sensing. Agreement measured using the R^2^ (which takes synaptic strength into account; mean *±* 1 s.d., 0.89 *±* 0.19) and the precision and recall (which measures agreement using only a binary classification; mean *±* 1 s.d.; precision 0.95 *±* 0.12, recall 0.84 *±* 0.15). Gray triangles indicate mean values. **h**, Simultaneous whole-cell and cell-attached recordings provide ground-truth presynaptic spikes associated with postsynaptic responses during single-target and ensemble stimulation. **i**, Example image showing locations of presynaptic and postsynaptic neurons, among other segmented ROIs. **j**, Example scenarios from a paired recording associated with successful optogenetic generation of presynaptic spikes. While trials (i) and (ii) resulted in correct predictions of presynaptic spiking by CAVIaR (i.e., are true positives), trials (iii) and (iv) were declared trials on which the stimulated neuron did not spike, either due to an unusually low-amplitude PSC or synaptic failure. Underlaid gray traces show raw PSC without NWD, black traces show PSC with NWD. Scales at bottom left apply to all subpanels. **k**, Performance of presynaptic spike inference over *n* = 4 paired recordings, separated by single-target vs ensemble stimulation and laser power. Performance of presynaptic spike inference does not depend on power or single-target vs ensemble stimulation (*p >* 0.05 for all pairs, dependent t-test), though we note that this lack of significance could be due to our sample size.

In some experimental preparations opsin-expressing neurons do not also express a separate nuclear fluorophore for straightforward identification of photosensitive cell nuclei [Bounds et al., 2021]. We therefore considered whether our connectivity mapping system could be used in the case where neuron locations are not known *a priori* (i.e., when mapping is performed “blind”). In this context, the field of view is divided into a three-dimensional grid and locations are stimulated one grid point at a time [Wang et al., 2007, Baker et al., 2016, Naka et al., 2019]. We performed both single-target and 10-target stimulation of grid points at 30 Hz and applied NWD and CAVIaR independently to the two sets of mapping data. Both single-target and ensemble stimulation identified a common ROI, and overlaying the stimulation grid on an image of the underlying tissue showed that this ROI aligned with a unique opsin-expressing neuron (Figure S23; see Figure S24 for full, multi-plane analysis of blind grid mapping experiment). This confirmed that NWD and CAVIaR could also facilitate rapid connectivity mapping in the blind stimulation regime.

We next wanted to confirm that CAVIaR inferences are well-calibrated. Specifically, since compressive approaches to connectivity mapping depend closely on the accuracy of the presynaptic spike inference step [Hu and Chklovskii, 2009, Shababo et al., 2013], we wanted to determine whether the spikes inferred by CAVIaR were allocated to the correct presynaptic neurons. We therefore performed multiple (n=4) paired recordings to obtain ground-truth spikes. We first used single-target stimulation to identify a likely presynaptic neuron in real time, then established a cell-attached patch clamp on the presynaptic neuron, and then randomly alternated between single-target and ensemble stimulation across multiple power levels while recording presynaptic spikes (Figure 4h,i). We confirmed that each of the neurons identified as connected using single-target stimulation following establishment of the patch clamp were also considered connected when the population was subsequently mapped using ensemble stimulation.

Paired recordings revealed a range of different scenarios associated with ground-truth spiking (Figure 4j). This included true positives, where CAVIaR correctly inferred a presynaptic spike from a given PSC evoked by single-target or ensemble stimulation (scenarios (i) and (ii) in Figure 4j). This also included false negatives, however. For example, in some cases CAVIaR incorrectly estimated that an unusually small PSC did not arise by a stimulation-evoked spike of the patched presynaptic neuron but rather from another, more weakly connected neuron that was stimulated at the same time (scenario (iii)). False negatives also occurred due to synaptic failure, where the presynaptic neuron successfully elicited, but did not transmit, a spike (scenario (iv)). Despite the latter two challenging scenarios, CAVIaR achieved high accuracy in its overall estimates of presynaptic spikes with no significant degradation in its ability to infer presynaptic spikes for individual neurons, even when stimulating ten neurons at once and over a range of laser powers (Figure 4k; all pair-wise differences not significant, p *>* 0.05, dependent t-test).

### CAVIaR estimates converge rapidly and accurately predict postsynaptic responses to novel holographic stimulation patterns

To go beyond single-target confirmation of the presence and magnitude of the synapses identified using CAVIaR (Figure 4f,g), we next asked whether the inferred connectivity could be used to predict the postsynaptic response to holograms targeting combinations of neurons that the model had never seen before. We also wanted to know whether ensemble stimulation was systematically engaging polysynaptic effects. For example, PV neurons are known to provide high levels of mutual inhibition [Pfeffer et al., 2013], which could be triggered when stimulating many PV neurons at once. In this case, other PV neurons would be inhibited, potentially reducing the number of driven PV neurons and resulting in smaller inhibitory PSCs than would be expected by CAVIaR.

To simultaneously answer these questions, we performed an out-of-sample testing method that we refer to as leave-one-hologram-out cross-validation (LOHO-CV, Figure 5a). Assuming that *H* different holograms are used in the experiment (where each hologram targets a different set of neurons), LOHO-CV works by running the CAVIaR algorithm on data corresponding to *H* − 1 holograms, such that the algorithm has never observed postsynaptic responses from simultaneous stimulation of the neurons targeted in the *H*th hologram. Then, samples from the posterior distribution over the model parameters are used to predict the mean postsynaptic response to hologram *H* at three different power levels, a process that includes sampling over the model’s uncertainty about whether spikes will be elicited by stimulation at any given laser power. The predicted response is compared against the held-out response, and the process continues with a different hologram selected to be held-out until the response to every hologram has been predicted.

**Figure 5:**
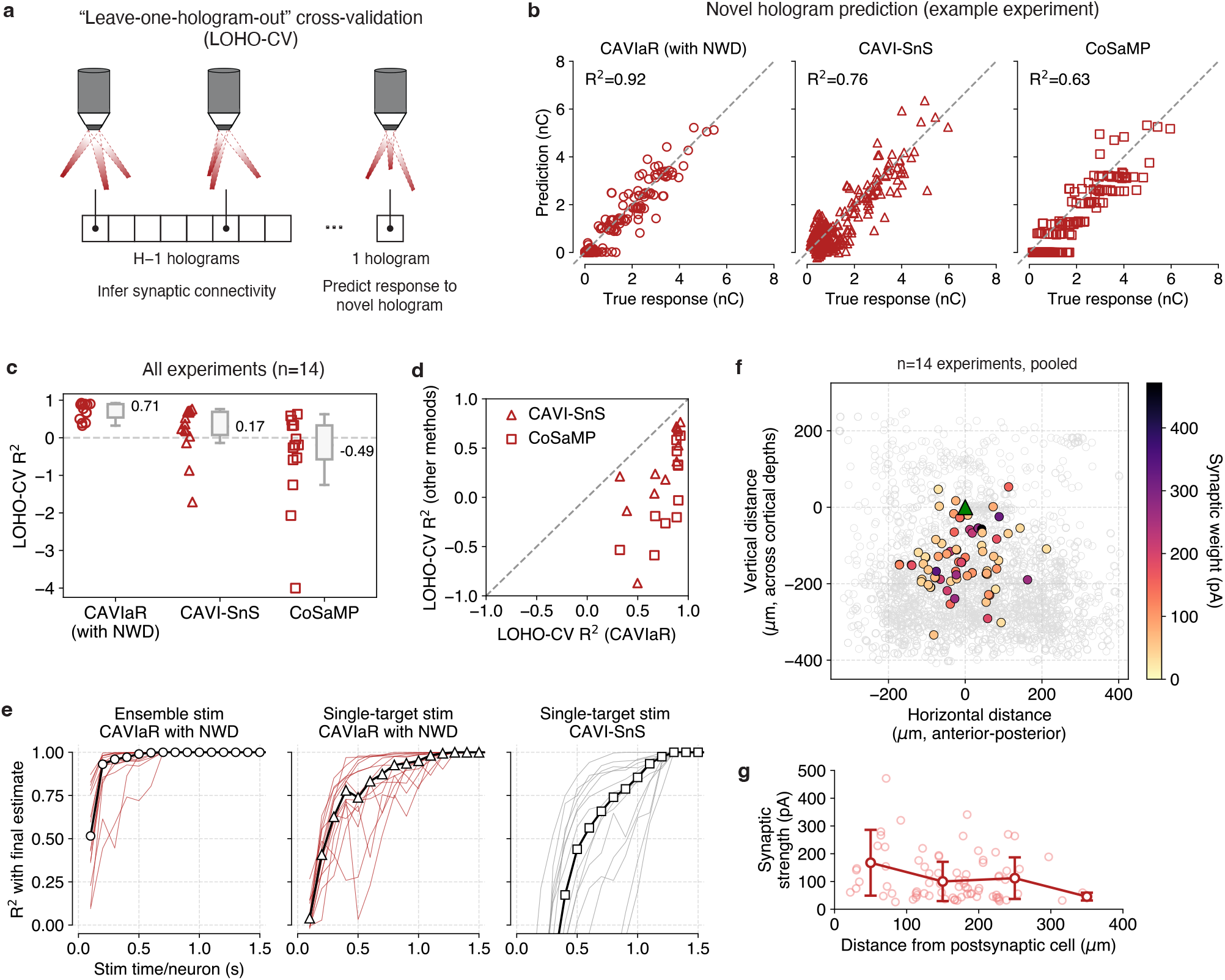
Predictive performance and convergence rates using model-based compressed sensing outperform existing methods. **a**, A “leave-one-hologram-out” cross-validation approach can be used to determine the predictive performance of connectivity mapping techniques. **b**, LOHO-CV R^2^ scores for CAVIaR, CAVI-SnS, and CoSaMP on an example mapping experiment, showing that CAVIaR obtains the highest accuracy. No method suggests recruitment of polysynaptic effects, which would manifest as systematic off-diagonal data points. For these comparisons, CoSaMP was provided with the number of connections identified by CAVIaR. **c**, Summary of LOHO-CV R^2^ scores across 14 different PV-pyramidal mapping experiments. Dashed gray line denotes an R^2^ of 0; scores below this line indicate poorer predictive performance than simply predicting the mean of the true responses without regard to which neurons were stimulated. **d**, Direct comparison of CAVIaR LOHO-CV R^2^ scores to CAVI-SnS and CoSaMP, showing that CAVIaR outperforms both methods on each dataset. **e**, Convergence time for proposed model-based compressed sensing technique (ensemble stimulation with CAVIaR and NWD) compared to conventional single-target mapping. Stimulation performed at 30 Hz. Ensemble stimulation applied to 10 targets at once. Total convergence time for each experiment calculated as the time per trial (33 ms for a 30 Hz stimulation speed) multiplied by the number of trials. For each experiment, the convergence time per neuron is obtained by dividing the total convergence time by the number of neurons in the experiment. Convergence time reported per neuron due to varying population sizes (larger populations take longer to map). Red/gray lines show convergence times for individual experiments. Each point is obtained by evaluating the R^2^ between the estimated connectivity at that time point and at the end of the experiment. Black line with markers shows median across experiments. **f**, Z-projected map of connection strength across 14 pooled experiments. Colored circles indicate position of identified presynaptic neurons, open gray circles indicate positions of all probed ROIs. Individual maps aligned to have postsynaptic neuron at location (0, 0). **g**, Decreasing synaptic strength as a function of distance from the postsynaptic neuron (14 experiments, pooled). A similar trend holds for connection probability (Figure S18). Faint circles show the strength of each connection; dark lines show mean *±* 1 standard deviation for connection strengths binned at 100 *µ*m.

LOHO-CV confirmed that CAVIaR achieved high accuracy in predicting responses to novel holograms (Figure 5b; example R^2^, 0.92; average over 14 experiments, 0.71). This indicated that CAVIaR did not miss synaptic connections that evoked large and reliable PSCs (the missed connections would appear as data points along the *y* = 0 axis in Figure 5b). Additionally, high LOHO-CV performance implied that CAVIaR did not make systematic errors arising due to polysynaptic effects, which would manifest as clear off-diagonal points in Figure 5b. We performed a similar cross-validation analysis for CAVI-SnS and CoSaMP, which achieved substantially lower performance compared to CAVIaR (Figure 5b, middle and right panels). Repeating the analysis across all 14 experiments showed that these two previously developed techniques exhibited systematic errors in their ability to predict responses to unseen holographic stimuli (Figure 5c, d), indicating that they did not learn connectivity consistent with the underlying physiology.

We next wanted to determine how efficient our protocol was for discovering synaptic connectivity as a function of continuous stimulation time (namely, without regard to experiment time devoted to pausing stimulation to measure access resistance, adjusting the seal of the patch pipette, etc.). To do so, we subsampled mapping data from the total set of available trials and, for each subset, applied NWD and CAVIaR to estimate connectivity. This allowed us to monitor the rate of convergence of CAVIaR to its final estimates in increments of tens of seconds. Note that because our principal aim was validation of the connectivity inferences themselves rather than a demonstration of speed-up specifically, each hologram was presented multiple (3) times at random throughout the experiment. This experimental design provided a means to characterize performance via LOHO-CV analysis, but could only establish a lower bound on mapping speed because simulations demonstrate that optimal speeds are obtained only when unique holograms are used on every trial (Figure S9b). Nevertheless, we found that our mapping protocol converged to an R^2^ of 0.9 with less than just 200 ms of stimulation time per neuron at 30 Hz (Figure 5e, left plot; obtained by normalizing mapping time by number of neurons in experiment). By comparison, single-target stimulation using CAVIaR and NWD reached an R^2^ of 0.9 in 800 ms per neuron (though with much higher variance; Figure 5e, middle plot), and single-target stimulation using the existing approach of CAVI-SnS required 1.1 seconds per neuron (Figure 5e, right plot), more than five times slower than model-based compressed sensing and (in accordance with our simulation and LOHO-CV results; Figure 3c, Figure 5c) with substantially lower predictive accuracy.

We also repeated the analysis by plotting the rate of convergence using single-target mapping and compressed sensing, but as a function of the number of times each neuron is stimulated and as a function of the number of stimulation trials instead of in units of time (Figure S15). This showed that single-target mapping converges in slightly fewer stimulations per neuron than compressed sensing (Figure S15a-c), presumably due to the fact that CAVIaR must solve a spike inference problem that is challenging for ensemble stimulation but straightforward for single-target stimulation. However, when plotted as a function of trials (Figure S15d-f) and as a function of trials normalized by population size (Figure S15g-i), compressed sensing converged considerably faster, as neurons are stimulated far more times over the course of an experiment using ensemble stimulation.

Aggregating the connections resulting from the use of ensemble stimulation, NWD, and CAVIaR recapitulated the characteristic [Hage et al., 2022] distance-dependent structure of PV-pyramidal connectivity (Figure 5f,g; Figure S18).

### Inference of synaptic connectivity from ensemble stimulation across multiple cortical cell types

Finally, we wanted to characterize how model-based compressed sensing performed across a variety of cortical cell-type combinations. We used an Emx-Cre line to express the more potent ChroME2s opsin in pyramidal neurons (which benefit from the greater potency of ChroME2s as the rheobase of pyramidal neurons can often be higher than interneurons, see e.g. the NeuroElectro database [Tripathy et al., 2014]) and an SST-Cre line to express ChroME2f in SST neurons. Then, in addition to our previous experiments mapping PV-pyramidal connections, we also mapped pyramidal-pyramidal (n=12 experiments), SST-pyramidal (n=8), and pyramidal-PV connections (n=5, Figure 6). In each case we used our previous experimental design, where we randomly interleaved single-target and ensemble stimulation trials, and used NWD and CAVIaR to infer connectivity independently from each trial type.

**Figure 6:**
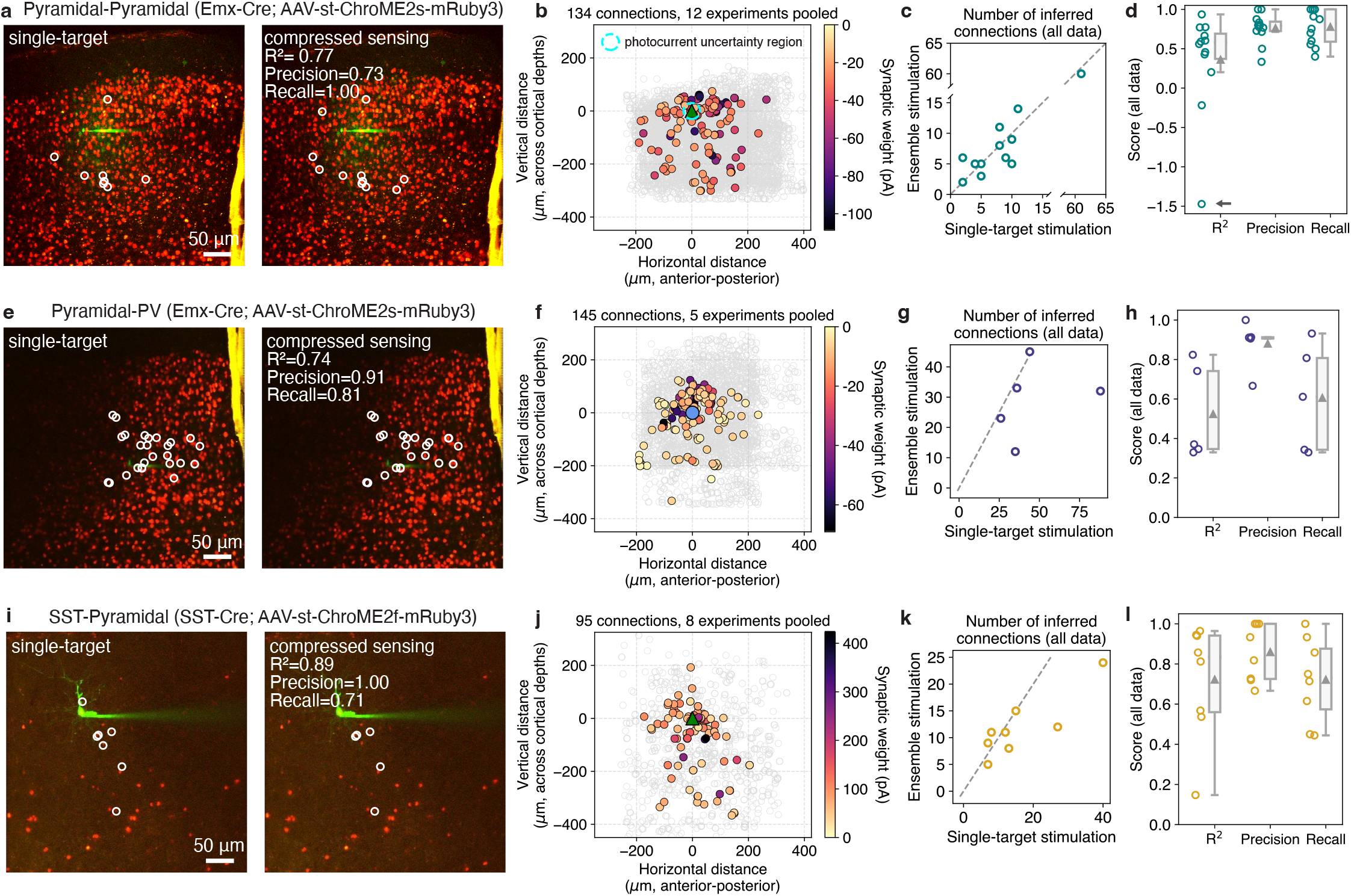
Model-based compressed sensing of synaptic connectivity applied to multiple cortical cell type pairs. **a**, Comparison of pyramidal-pyramidal connectivity maps obtained on the same population of neurons using single-target stimulation and model-based compressed sensing (10-target ensembles, 30 Hz stimulation, connections identified using CAVIaR with NWD). Experiment performed over 5 planes, image shows z-projection. Agreement between synaptic weights estimated by CAVIaR independently from single-target stimulation and ensemble stimulation trials. **b**, Map of synaptic connections identified using ensemble stimulation across 12 pooled experiments. Dashed cyan circle represents region with 30 *µ*m radius where synaptic connections are most uncertain due to potential photocurrent contamination in opsin-positive postsynaptic neurons (however see Figure S19). **c**, Number of connections identified by CAVIaR using both single-target and ensemble stimulation across 12 experiments. Dashed gray line shows identity. **d**, Agreement between connectivity maps obtained using single-target stimulation compared to model-based compressed sensing. Mean R^2^, 0.36; median R^2^, 0.57; mean precision, 0.76; mean recall, 0.77. Gray arrow indicates outlier (c.f. Figure S22). Gray triangles indicate mean values. **e–h**, same as a–d, but for mapping pyramidal-PV connections over 5 experiments. Blue circle represents postsynaptic PV neuron. Mean R^2^ in panel h, 0.52; mean precision, 0.88; mean recall, 0.6. Lower recall for pyramidal-PV connections could arise due to the faster and more irregular PSC kinetics encountered compared to PV-pyramidal connections. **i–l**, Same as a–d, but for mapping SST-pyramidal connections over 5 experiments. Mean R^2^ in panel l, 0.72; mean precision, 0.86; mean recall 0.72.

Our experiments mapping pyramidal-pyramidal connections involved probing hundreds of potential presynaptic candidates per cortical slice (Figure 6a,b; mean number of targets, 486; minimum, 214; maximum, 687). This experimental preparation posed a particular challenge because the high opsin expression density made it difficult to identify any pyramidal neurons that were entirely opsin-negative, and consequently we typically patched an opsin-positive postsynaptic pyramidal neuron (fraction of pyramidal-pyramidal mapping experiments with opsin-positive postsynaptic neuron, 10/12). When probing nearby presynaptic candidates we thus frequently encountered direct photocurrent artifacts in our whole-cell recordings that could affect our ability to discriminate synaptic connectivity (i.e. direct photocurrents could increase the false-positive rate in the immediate vicinity of the patch-electrode). However, because NWD is trained to isolate PSCs occurring within a specified window (typically 3-12 ms after stimulation), confounding photocurrents (which begin rising immediately upon stimulation) were suppressed by the demixing process (Figure S19). Following the application of CAVIaR to infer connectivity, there was (as expected) some variability between connectivity maps obtained using single-target stimulation and model-based compressed sensing, but the identity, strength, and overall number of identified connections tended to be conserved across experiments (Figure 6a, c-d; mean R^2^ between single-target and model-based compressed sensing connectivity vectors, 0.36; mean precision, 0.76; mean recall 0.77). Though the mean R^2^ was lower than in the PV-E condition, this was largely due to an outlier experiment (Figure S22), and was not representative of the distribution at large (median R^2^, 0.57).

Mapping pyramidal-PV connections involved probing similar numbers of potential presynaptic candidates as in our pyramidal-pyramidal maps due to a similarly high opsin expression density and intrinsic cell-count (Figure 6e, mean number of targets across 5 experiments, 602; minimum, 562; maximum, 661). However, because in this preparation opsin is expressed specifically in pyramidal neurons, when recording from PV neurons we faced no risk of direct photocurrent artifacts. Across 5 experiments, connectivity maps obtained using single-target stimulation and model-based compressed sensing achieved a mean R^2^ of 0.52, a mean precision of 0.88, and a mean recall of 0.6 (Figure 6f-h).

Finally, we considered whether using compressed sensing to map connections from SST neurons to pyramidal neurons could pose an additional challenge because SST-pyramidal connections are largely axo-dendritic, potentially causing significant filtering and attenuation of PSCs through the dendrites of pyramidal neurons [Cristo et al., 2004, Wang et al., 2004, Dorsett et al., 2021], though prior work has shown the feasibility of using single-target stimulation (via two-photon glutamate uncaging) to map connections of this type [Fino and Yuste, 2011]. Across 8 experiments we found that, while some variability persisted, the connectivity maps obtained using single-target stimulation and model-based compressed sensing were highly conserved, despite this more challenging experimental setting (Figure 6i-l; mean R^2^, 0.72; mean precision, 0.86; mean recall, 0.72; mean number of targets, 129; minimum 51; maximum 175).

By performing a relatively small number of experiments (5-14) for each of the mapped populations, we acquired connectivity maps that corresponded to the probing of thousands of presynaptic candidates within the local volume. These maps recapitulated the increased connection probability in the proximity of the postsynaptic neuron (≤100 *µ*m around the cell) characteristic of connectivity mapping studies (Figure 6b,f,j), which typically require pooling together many more smaller-scale experiments [Baker et al., 2016, Seeman et al., 2018, Hage et al., 2022, Campagnola et al., 2022]. While a detailed analysis of the spatial properties of individual maps of connectivity is beyond the scope of this study, our results indicate the possibility of investigating map-to-map variability (Figure 4d,e; supp figs) [Nikolenko et al., 2007]. Further, our results show the feasibility and utility of compressive connectivity mapping across a diversity of cortical cell-type combinations, despite challenges arising due to synaptic physiology or experimental preparation (i.e., direct photocurrent artifacts).

## Discussion

Uncovering how neural computations are implemented in the cortex requires knowledge of the existence and strength of the underlying synaptic connections. We therefore developed a pair of complementary computational tools (NWD and CAVIaR) that together enable high-speed acquisition of large-scale synaptic connectivity maps using two-photon holographic optogenetics. Other studies that develop compressive connectivity mapping techniques either use generic compressed sensing algorithms or have only been demonstrated in simplified simulations [Hu and Chklovskii, 2009, Shababo et al., 2013, Draelos and Pearson, 2020, Navarro and Oweiss, 2023, Chen et al., 2023]. Here, however, we show that considerable methodological extensions over these previous algorithms are required to ensure that connectivity is accurately determined in light of the substantial sources of biophysical variability that are intrinsic to real holographic stimulation experiments. We found that use of NWD and CAVIaR provided a mapping speed-up of over an order of magnitude compared to existing approaches in realistic settings, granting the speed and scalability necessary to collect large-scale maps of connectivity within individual experimental sessions.

To implement and validate these tools in real mapping experiments we used cutting-edge optogenetic techniques that offered several key advantages. Namely, since the ChroME2f opsin is potent with fast off-kinetics, we could titrate the pulse width and range of stimulation laser powers such that with high probability stimulation generated either 0 or 1 presynaptic spike(s), with minimal instances of more than one spike (Figure S1), facilitating tractable presynaptic spike inference. By comparison, potent opsins with slow off-kinetics (such as ChRmine [Marshel et al., 2019]) elicit larger and less predictable presynaptic spike counts [Sridharan et al., 2022] that could make precisely inferring synaptic weights more computationally challenging. Finally, an important assumption of CAVIaR is that, everything else held equal, the probability of eliciting a spike increases monotonically with laser power (on average). While the ChroME2 opsins used in this study exhibit such behavior, some opsins have been shown to change the probability of spiking a neuron non-monotonically under certain laser powers (described in ref. [Sridharan et al., 2022]). It is therefore critical that an appropriate opsin or range of powers is determined during a calibration session prior to mapping using CAVIaR, such that monotonicity is upheld and that no more than one spike is elicited by stimulation.

We also relied on the fact that ChroME2f expression is targeted to the soma through a fusion with the Kv2.1 tag [Mardinly et al., 2018, Sridharan et al., 2022], reducing the likelihood of triggering spikes in untargeted neurons by off-target stimulation of nearby neurites [Baker et al., 2016, Shemesh et al., 2017]. Moreover, we used two-photon holographic light sculpting to illuminate entire somas at once [Papagiakoumou et al., 2010, Hernandez et al., 2016, Pégard et al., 2017] and randomly switched the stimulation laser between three or more powers, effectively probing for presynaptic neurons at multiple excitation levels. Stimulating with low power further reduces the risk of off-target stimulation, but may fail to elicit spikes in neurons with low opsin expression or high rheobases. Notably, it cannot in general be known ahead of time whether a lack of optogenetically evoked PSCs was due to the stimulated neuron not being connected or because of a failure to elicit spikes, an ambiguity that can only be resolved through high power stimulation. On the other hand, stimulating with high power increases the risk of off-target stimulation but elicits spikes with high probability. We therefore elected to use a randomized design that probed neurons at multiple power levels, but that still minimized the risk of off-target stimulation (Figure S1).

One potential concern with our proposed connectivity mapping approach is that it may engage mechanisms for short-term plasticity (i.e., synaptic facilitation or depression) given our suggested high speed of stimulation and the fact that entire ensembles of neurons are stimulated at once. To help calibrate mapping protocols, we derived a simple combinatorial expression for the expected interstimulus interval (ISI, in seconds) for any given neuron (Methods). Assuming connectivity from *N* total neurons is mapped by stimulating a random ensemble of size *R* on each trial at a speed of *f* Hz, the mean ISI is 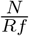. Thus, a typical experiment mapping ∼300 neurons with 10-target stimulation at 30 Hz yields an expected stimulation return time of 1 second. The risk of engaging intrinsic plasticity – resulting from repeated depolarization of presynaptic candidate neurons – can similarly be assessed with this expression. However, depolarizing ChroME2-expressing neurons at rates even up to 40 Hz has been shown to only minimally induce intrinsic plasticity effects [Sridharan et al., 2022], and direct analysis of our data did not suggest recruitment of intrinsic plasticity (Supplementary note 2).

An additional concern associated with ensemble stimulation is the recruitment of potential polysynaptic network effects arising due to recurrent connectivity in the cortex [Oldenburg et al., 2022]. However, in slice experiments we do not expect this to be a major factor influencing the quality of our inferences compared to *in vivo* conditions. Further, as CAVIaR is designed to model monosynaptic connections only, we could use LOHO-CV to assess whether polysynaptic effects were confounding our analysis. This was possible because CAVIaR would not be able to capture the variance generated by polysynaptic pathways. While LOHO-CV therefore suggested that polysynaptic effects did not compromise our compressed sensing approach (Figure 5b,c), the presence of polysynaptic effects likely depends on brain region and number of simultaneously illuminated targets.

In order to validate our compressed sensing approach, we mapped each neural population using both single-target and ensemble stimulation to determine how closely the resulting connectivity maps align. However, in practice we would not expect an experimenter to perform both single-target and ensemble stimulation within each experiment, as this would greatly inflate the required experiment time and defeat the purpose of our approach. Instead, to determine whether mapping results are accurate when only having access to ensemble stimulation, we would recommend using LOHO-CV – this enables calibration of the accuracy of the inferred connections without the need to reference single-target stimulation. If the LOHO-CV R^2^ is poor (due to e.g. deteriorating recording quality or a potential miscalibration in the optical system), the associated experiments could be excluded from further analysis.

A critical challenge in validating compressed sensing approaches using single-target stimulation is ensuring that the power delivered to target neurons is consistent between single-target stimulation and ensemble stimulation. While we have taken steps to calibrate our optical system, there can still be small yet unavoidable differences in the amount of two-photon excitation ultimately delivered to neurons (Figure S2). These subtle differences in power can sometimes evoke spikes in single-target stimulation differently to ensemble stimulation, leading to differences in which neurons are identified as being presynaptic. While our results almost always show high precision (maximum precision in this study, 0.95 for PV-pyramidal experiments; minimum, 0.76 for pyramidal-pyramidal experiments), the recall is almost always lower (maximum recall, 0.84 for PV-pyramidal; minimum, 0.6 for pyramidal-PV). This suggests that although the connections found by compressed sensing are almost always replicated by single-target connectivity mapping, disagreements between the two approaches primarily stem from putative connections being missed by compressed sensing.

What could explain this discrepancy? Our LOHO-CV analysis shows that for holograms targeting novel combinations of neurons (i.e. groups of neurons that have never been stimulated together before), we achieve excellent prediction performance (average R^2^ across 14 PV-E experiments, 0.71). If CAVIaR were simply missing connections it should have found, this would be reflected in the distribution of LOHO-CV R^2^ values, which did not appear to be the case in our data (Figure 5c). A possible explanation for the discrepancy between single-target and compressed sensing approaches under some experimental conditions is therefore that there is a small decrease in the amount of laser power ultimately delivered to some target neurons when performing ensemble stimulation compared to single-target stimulation, despite our best efforts to equalize this in calibration experiments. The extent to which such “power gaps” have been addressed during calibration should therefore be kept in mind when interpreting the precision and recall of compressed sensing methods.

We demonstrated the applicability of NWD and CAVIaR by mapping connectivity between three different cell-types (PV, SST, and pyramidal neurons), using two ChroME variants (ChroME2f and ChroME2s), and using both targeted and “blind” experimental designs. In every case, application of our tools led to a substantial improvement in signal quality and speed of connectivity mapping. Even in the case of single-target stimulation, our methods should still be of high practical value. NWD alone can ensure that single-target stimulation can be applied at much higher speeds than has previously been performed in practice. Indeed, we propose using NWD to enable 30-100 Hz stimulation rather than 10 Hz as in prior studies (e.g., refs. [Hage et al., 2022, Printz et al., 2023, Chen et al., 2023]), which immediately yields a 3-10x speedup in connectivity mapping. Finally, the CAVIaR algorithm (combined with NWD) is useful for single-target connectivity mapping due to its ability to adapt to spontaneous activity and eliminate spurious connections where PSCs happen to decrease with increasing laser power.

While in our experiments we typically mapped connectivity from several hundred neurons, this was only limited by factors outside of the influence of NWD and CAVIaR, such as the number of neurons targetable by our SLM or the rate at which phase masks could be computed. Our simulations predict that the scale and throughput of compressive connectivity mapping will only grow with the adoption of experimental steps to increase the targetable field of view (e.g., by translating the microscope or SLM stimulation field), with larger commercially available SLMs, with the use of holographic mesoscopes [Abdeladim et al., 2023], or with even more potent opsins enabling stimulation of more targets for the same laser power. Such advances would allow for precise, low-latency holographic control over larger neural ensembles or for mapping larger population sizes than those used here.

An important direction for future work is the incorporation of simultaneous calcium or voltage imaging into our connectivity mapping experiments [Baker et al., 2016]. Such an approach should, in principle, resolve exactly which presynaptic candidate neurons are activated on each trial. We expect that this would enable the mapping of larger neural populations and with greater certainty, as fewer trials are required to overcome ambiguities about which neurons elicited presynaptic spikes. However, this approach would not be without its own difficulties: our stimulation regime aims to elicit just a single spike with each pulse, resulting in low-amplitude calcium or voltage transients compared to typical imaging conditions, and synaptic failures could still occur. Additional computational methods would therefore likely be required to overcome these challenges in order to fully utilize such multimodal data.

Another direction for future research is the modelling and inference of short-term synaptic plasticity. However, this inference problem could be challenging as it is difficult to precisely resolve amplitude changes in the individual PSCs that summate to give the response to ensemble stimulation. A more effective strategy in practice would likely be to first perform compressed sensing to identify connected neurons, and then switch to single-target stimulation to probe plasticity-related variables in finer detail.

Our results indicate that NWD and CAVIaR can substantially reduce the cumulative time spent stimulating tissue to map connectivity. However, the speed of a mapping experiment can also be impacted by other factors, including the time required to identify and segment neurons, monitor the access resistance of the patch clamp, and compute hologram phase masks. Each of these factors similarly affects mapping performed using single-target stimulation and ensemble stimulation. While acquiring an image stack to identify presynaptic candidates is required for all experiments using targeted stimulation, measuring access resistance poses an additional trade-off between the speed of a mapping experiment and the desire to make regular quality checks. An ideal experimental design would therefore proceed with the longest periods of uninterrupted stimulation possible to yield the maximal benefits of NWD and CAVIaR, but with brief (∼200 ms) automatic pauses included to check access resistance. The attending experimenter could then be prompted to address inadequate access only when necessary. In the case of phase mask generation, efficient computation is critical since otherwise experiment time could be dominated by this step. We expect that high-speed phase mask estimation methods, such as those based on deep learning [Eybposh et al., 2020, Eybposh et al., 2022], could play an increasingly important role in rapid connectivity mapping.

Combining NWD and CAVIaR with two-photon holographic optogenetics should enable new experiments where, with just a few minutes of rapid ensemble stimulation, one can obtain an extensive map of connectivity. This could allow for very high throughput collection and screening of synaptic connectivity maps from brain slices, or for future *in vivo* applications [Pégard et al., 2017, Mardinly et al., 2018, Chen et al., 2023] where synaptic connectivity can be directly related to functional activity. Critically, *in vivo* experiments are impacted by higher background rates of spontaneous activity and more prevalent polysynaptic effects. While further validation must be performed to ensure accuracy *in vivo*, the impact of each of these impediments is demonstrably reduced by our computational methods (Figure S26), implying that NWD and CAVIaR may be important tools for precise and scalable mapping in this experimental regime.

## Acknowledgements

The authors thank Kenneth Kay, Darcy Peterka, Ben Shababo, and Shizhe Chen for many insightful discussions and helpful suggestions. We thank Uday Jagadisan for technical assistance related to the optical setup and Karthika Gopakumar for helping with animal husbandry and the opsin expression procedure. This work was funded by NIH awards 1RF1MH120680 and 1U19NS107613-01 to HA and LP, and K99NS135649 to MAT. MAT, BA, and LP were supported by the Gatsby Charitable Foundation and NSF NeuroNex award 1707398.

## Author contributions

MAT, MG, HA, and LP conceived of the project. MG designed and performed all experiments, with assistance from MS during the initial project phase. MAT developed the computational methods together with LP, with contributions from BA. MAT performed all simulations. MAT and MG performed all analyses. MAT and MG wrote the manuscript with input from all authors.

## Code availability

CAVIaR is implemented in Python using JAX [Bradbury et al., 2018], can be run on GPU, and is freely available at https://github.com/marcustriplett/circuitmap together with a PyTorch Lightning implementation of NWD.

## Methods

### Experimental methods

#### Animals

All experiments on animals were conducted with approval of the Animal Care and Use Committee of the University of California, Berkeley. In all experiments we attempted to use male and female mice equally. Mice used for experiments in this study were transgenic Emx-Cre, PV-Cre, or SST-Cre mice obtained by crossing the corresponding lines in-house with a wild type (CD-1 (ICR) white strain, obtained from Charles River). Mice were housed in cohorts of five or fewer in a reverse light:dark cycle of 12:12 hours, with experiments occurring during the dark phase.

#### Opsin expression method

We used a neonatal injection procedure to induce expression of ChroME2s (AAV9.CAG.DIO.ChroME-ST.P2A.H2B-mRuby6) in the visual cortex of Emx-Cre animals or ChroME2f (AAV9.CAG.DIO.Chrome2f.P2A.H2B-mRuby3.WPRE.SV43) in PV-Cre and SST-Cre animals. Both constructs expressed the mRuby3 fluorophore in opsin-positive cell nuclei. Young pups at P3 or P4 were anaesthetized by placing them on ice for approximately 3 minutes. Next, each animal was stabilized under the nanoliter-injector (WPI) and a small portion (30 nL/injection) of virus was injected directly in 3-5 places around V1 via the skin and skull and at 3-5 depths to target L2/3. After the procedure the animal was placed on a heating pad until it recovered. At the end of the procedure the injected litter was returned to their cage with their parents and housed together until reaching approximately 21 days of age.

#### 3D-SHOT holography setup

All experiments were performed using the 3D-SHOT multiphoton holography setup (see [Mardinly et al., 2018] for details). Briefly, the setup was custom built around a commercial Sutter MOM (movable object microscope) platform (Sutter Instruments) and combined a 3D photostimulation path, a fast resonant-galvo two-photon raster scanning imaging path and a wide-field one-photon epifluorescence/IR (infrared) transmitted light imaging path. The stimulation and imaging beams were merged together using a polarizing beamsplitter placed before the microscope tube lens.

A femtosecond fiber laser was used for two-photon photostimulation (Monaco 1035-80-60; 1040 nm, 1 MHz, 300 fs, Coherent). The stimulation laser was directed onto a blazed diffraction grating (600 l/mm, 1000 nm blaze, Edmund Optics 49-570) for temporal focusing. In order to be able to utilize the total available laser power (60 W laser output), the beam was enlarged by a factor of 2.5 to prevent heat damage of the grating surface. The spot on the grating was relayed onto a rotating diffuser where it formed a temporally focused spot. The rotating diffuser was used to both randomize the phase pattern imprinted on the temporally focused spot and to expand the beam in the direction orthogonal to the temporal focusing direction and fully fill the spatial light modulator (HSP 1920 1920×1152 pixels, Meadowlark Optics). The SLM plane was relayed through 4f systems to the back aperture of an Olympus 20x water immersion objective, resulting in a custom 3D distribution of temporally-focused spots at the focus of the objective. Holographic phase masks were calculated using the iterative, in-house written and GPU-optimized Gerchberg-Saxton algorithm [Gerchberg, 1972] and the intensity distribution was corrected to accommodate for diffraction efficiencies (compensating for possible attenuation effects from the SLM when targeting different regions for stimulation).

The two-photon imaging path relied on a Ti:sapphire laser, Mai Tai (Spectra Physics), with external power control via Pockels cell (Conoptics, Inc). For fast raster scanning, the system was equipped with conjugated 8 kHz resonant galvo-galvo systems. The imaging path hardware was controlled by ScanImage software and custom Matlab code was used to control the spatial light modulator for targeted photostimulation and synchronize with imaging.

Epifluorescent one-photon excitation was via a Spectra X (Lumencor) light source filtered by an appropriate excitation filter set. For slice transillumination we used a 750 nm and IR diffuser. The image was collected using an Olympus 20x magnification water-immersion objective and a CCD camera and displayed on a screen enabling targeted patch clamping.

#### Slice electrophysiology

*In vitro* slice recordings were performed on 300 *µ*m-thick coronal slices coming from 4–6-week-old animals expressing opsin in L2/3 of V1. During slicing the level of opsin expression was checked using a simple laser light to visualize the targeting of the opsin to V1 and the general brightness of the mRuby3 nuclear marker.

Whole-cell patch-clamp protocols were performed in ACSF (artificial cerebrospinal fluid) perfusion solution (in mM: NaCL 119, NaHCO3 26, Glucose 20, KCl 2.5, CaCl 2.5, MgSO4 1.3, NaH2PO4 1.3) in temperature-controlled (33^*°*^C) conditions. Patch pipettes (4-7 MΩ) were pulled from borosilicate glass filaments (Sutter Instruments) and filled with a Cesium (Cs2+)-based internal solution (in mM: 135 CeMeSO4, 3 NaCl, 10 HEPES, 0.3 EGTA, 4 Mg-ATP, 0.3 Na-GTP, 1 QX-314, 5 TEA-Cl, 295 mOsm, pH=7.45) also containing 50 *µ*M Alexa Fluor hydrazide 488 or 594 dye (ThermoFisher Scientific). For loose-patch recordings the pipettes were filled with standard ACSF. Data was recorded at 20 kHz using a 700b Multiclamp Axon Amplifier (Molecular Devices). The headstage with the electrode holder (G23 Instruments) was controlled by a motorized micromanipulator (MP285A, Sutter Instruments). All data was acquired and analyzed with custom code written in Matlab using the National Instruments Data Acquisition Toolbox.

#### Learning physiological point spread functions

An opsin-positive cell was loose-patched at various slice depths between 20-90 *µ*m and a volume of tissue surrounding that cell was then probed using a dense grid of holograms. The total size of the grid was 65×65×75 *µ*m, resulting in a 10×10×7 voxel grid. The interval between the centers of the hologram targets were 5 pixels (6.5 *µ*m) in the x/y dimensions and 12.5 *µ*m in the z dimension. The phase masks of the grid holograms were pre-computed and stored in memory before performing the measurement. Two versions of the physiological point spread function experiments were performed. One where each hologram was a single spot randomly selected from the grid (n=8 experiments), and another where each hologram contained 10 spots: 1 spot randomly taken from the grid and 9 spots (either fixed or randomly placed) outside the grid (n=7 experiments).

Evoked spiking in response to 3-5 ms laser stimulation (30 Hz) across powers between 10-60 mW/spot were collected and analysed using a custom written script in Matlab. Each trial was randomly repeated 5-7 times. Evoked spiking was calculated by simple thresholding and the mean spiking probability across each grid point and power was calculated for single and 10-spot ensemble experiments. A Gaussian fit to the per-cell normalized mean spiking probability data was applied to estimate the FWHM of the spiking physiological point spread function.

#### Whole-cell targeted mapping experimental protocol

First, a L2/3 opsin-negative cell of pyramidal shape was sealed on (below 40 *µ*m of cortical depth, opsin absence judged by fluorophore presence). The tissue was then quickly imaged with approximately 40 frames at 4-5 planes spaced by 25 *µ*m. The cell was always positioned to be in the second plane from the top within the collected stack. The position of the presynaptic candidates were automatically identified by an in-house algorithm detecting round shapes in a specified FOV due to the presence of mRuby3 fluorophore in the opsin-positive cell nuclei.

Next, we computed 20-25 different sets of holograms, where every set of holograms was designed so that every target was stimulated once. Each hologram either targeted an ensemble of presynaptic candidates, or targeted one cell at a time. Further, each set of holograms was organized into sweeps, where each sweep was composed of a baseline period, followed by a single set of holographic stimulation trials, and ended with a short baseline period. A single set of ensemble holograms was obtained by randomly partitioning all targets into 10-target ensembles (without repeating any targets). Additional sets of holograms were obtained by repeatedly performing this random partitioning.

For example, an experiment probing 100 presynaptic candidates would contain data from 45 sweeps of 100 single-target holograms across 3 powers (15 repetitions per power), and 20 sweeps representing 20 sets of 10-target holograms. Each such set of 10-target holograms would be comprised of 100 targets split into 10 different 10-target holograms.This would ultimately result in 200 unique ensemble holograms organized in 20 sweeps. Each sweep would be performed across 3 powers and repeated 3 times per power (leading to 180 ensemble sweeps). In total, this would result in 225 sweeps (45 single-target sweeps and 180 ensemble sweeps). Importantly, the order of all holograms and powers within each repetition and sweep was randomized.

Stable access to the cell was obtained during hologram computation. The cell properties (Rm and Rs) were checked and the response to one-photon photocurrent was characterized to informally determine the overall level of opsin-expression before proceeding with connectivity mapping experiments using two-photon stimulation. Following the completion of hologram computation, presynaptic candidates were mapped using randomly interleaved experimental trials containing single-target holograms or samples from the set of 10-target holograms. Each single-target hologram was repeated 7-15 times, while ensemble holograms were repeated up to 3 times. All stimulations were performed across 3-4 laser powers in both mapping regimes.

The Rm and Rs parameters were controlled and frequently logged during the mapping protocol. Every 10 trials (or 20-180 seconds, depending on the number of presynaptic targets and the power condition), a series of hyperpolarizing steps was applied and the Rm and Rs logged. During offline analysis, trials when Rs significantly changed or were above 30 MOhm were excluded from the data. Experiments where zero or one connection(s) were identified were considered as having insufficient opsin expression and excluded from analysis.

For pyramidal-pyramidal mapping, the postsynaptic neuron was targeted based on its characteristic pyramidal shape and the absence of an mRuby3 signal in the cell. In an area with high opsin expression it was difficult to identify a completely opsin-negative cell. Thus, for a large portion of putatively opsin-negative cells, stimulating near the electrode still elicited a small direct photocurrent artifact (with a typical amplitude of 50-300 pA). For mapping pyramidal-PV connections, the postsynaptic cell type was confirmed to be PV by observing a fast-spiking response type as a result of a typical current step protocol.

#### Power calibration for single and ensemble target

To ensure that the connectivity maps obtained using single-target and ensemble stimulation were comparable, we carefully characterized the power needed to evoke similar photocurrent amplitudes at each presynaptic candidate in both stimulation regimes. To do this, a set of whole-cell experiments was performed in several cell types (pyramidal, PV, and SST). In each such experiment, an opsin-positive cell was patched and a direct opsin-photocurrent was recorded while all other experimental parameters were kept the same as in the connectivity mapping experiments. We used a range of powers that were typical for our connectivity mapping protocols. The requested laser power was a multiple of the diffraction efficiency of a given hologram (either single-target or ensemble) and power per cell multiplied by a number of targets in a hologram for a given condition.

In these experiments (Figure S2), we compared the opsin-photocurrent amplitude evoked in response to stimulation of a single target (the patched opsin-positive cell) with the opsin-photocurrent amplitudes evoked by ensemble holograms containing the patched opsin-positive cell. This set of experiments provided a measurement of the difference in the power delivered to the patched-cell between these two stimulation regimes. Correcting this difference required ∼35% additional power to the ensemble holograms (Figure S2b-d). Applying the correction compensated for the necessary photocurrent and was implemented in all mapping experiments (Figure S2e-g).

#### Pair-patch experimental protocol

The pair-patch mapping protocol was similar to the single-patch protocol, but contained an additional “pre-mapping” step. After establishing stable access to the opsin-negative pyramidal cell, all presynaptic candidates were probed with 10 trials of single-power, single-target holograms. A fast online analysis based on the z-score provided the coordinates of cells that responded to this simple connection screening procedure. One such identified cell would be approached and loose-patched with a second electrode while monitoring the stability of the postsynaptic recording. The main prerequisite for deciding which cell would be loose-patched (from a handful of putatively connected cells obtained via our simple screening procedure) was the ability to identify the cell in the slice under IR illumination and its accessibility. While the loose patch was established, the surrounding area would be re-imaged to update the positions of the presynaptic candidates. Mapping would then be performed according to the above-described procedure, while recording both postsynaptic responses and presynaptic spiking.

#### Whole-cell grid mapping experimental protocol

Opsin-negative cells (or those expressing minimal amounts of opsin) were whole-cell patched at various slice depths between 30-90 *µ*m. Cells were voltage-clamped at a holding potential of -70 mV. After establishing stable access to a cell, we acquired its response to one-photon pulses of increasing duration (1, 3, and 5 ms pulses at 10 Hz). A volume of tissue surrounding that cell would then be probed using a dense grid of holograms across 3-4 powers. The total size of the grid was ∼160×160×100 *µ*m, resulting in a 25×25×4 voxel grid. The distance between the centers of the hologram targets was 5 pixels (6.5 *µ*m) in the x/y dimensions and 25 *µ*m in the z dimension. The phase masks of the grid holograms were pre-computed and stored in memory before performing the measurement. Recording parameters were frequently checked and logged by pausing the mapping protocol and probing the cell with a series of hyperpolarizing pulses.

### Computational methods

#### Neural waveform demixing network architecture and training

The NWD network is a sequential U-Net [Ronneberger et al., 2015] that forms compressed representations of 45 ms single-trial PSC traces via a “contraction path,” and generates a PSC waveform at the original trial length but with confounding synaptic currents removed via an “expansion path.” Each block in the contraction path consists of a 2x temporal decimation step, a 1d convolution, a batch-norm step, and a rectified linear activation function. Each block in the expansion path consists of a “transposed” 1d convolution (also known as a fractionally strided convolution, or as a pseudo deconvolution), a batch-norm step, a rectified linear activation function, and a 2x linear interpolation step. There are four contraction blocks and four expansion blocks, with the outputs of the contraction blocks submitted to the expansion blocks at the corresponding temporal resolution through skip connections as in the original U-Net architecture.

The NWD networks used in this study were trained using 50,000 simulated PSC traces. The simulated PSCs obeyed the following generative structure. First, a template PSC *ϱ* (·; *τ*_*r*_, *τ*_*d*_, Δ) with parameters *τ*_*r*_, *τ*_*d*_, Δ took the form

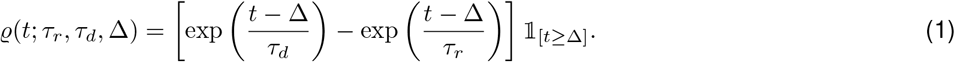

Every simulated PSC *c*(*t*) was the sum of a random number of templates, accounting for the stimulation of multiple connected presynaptic cells,

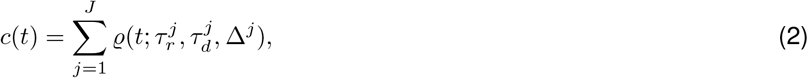

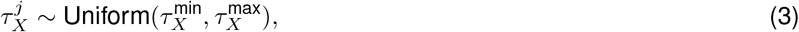

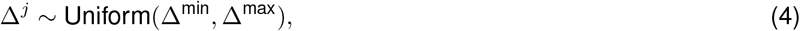

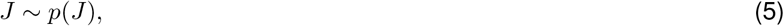

where *X* ∈ {*r*, diff} with *τ*_*d*_ = *τ*_*r*_ + *τ*_diff_ to ensure *τ*_*d*_ *> τ*_*r*_, and where *p*(*J*) determined the probability of selecting *J* ∈ N (including the possibility of *J* = 0). The PSC *c*_*k*_ for training example *k* was then given as a function of time *t* by

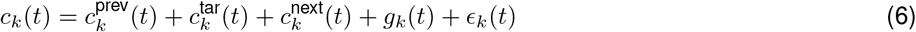

where 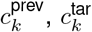 and 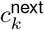 refer to PSCs from the previous trial, the target trial, and the next trial(s), which reproduce the “overlapping trials” effect when stimulating at high speeds. Note that 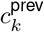 is not the same as 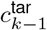 as each trial was simulated entirely independently. Here *g*_*k*_(*t*) represents temporally-correlated noise from a Gaussian process,

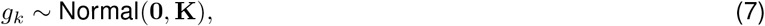

with the covariance matrix **K** defined by the radial basis function kernel with noise variance *σ*_scale_ and characteristic lengthscale 𝓁_gp_

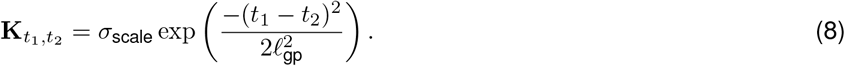

Finally, *ϵ*_*k*_(*t*) represents uncorrelated noise sampled from a univariate Gaussian,

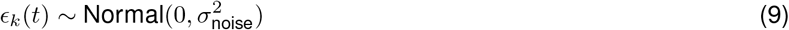

where 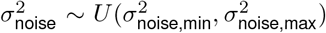. The NWD network is trained to infer 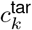 from 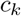 by minimizing the mean-squared error loss.

The time constants of the training data were selected to match the PSCs recorded in experiments via an interactive simulator that overlaid the template waveform on experimental traces. We found that the NWD network performed best when trained separately on training data matched to either excitatory or inhibitory synaptic currents depending on the data being processed. Indeed, because opsin is expressed only in a specific, genetically-defined cell type at once, we would not expect stimulation to recruit combinations of monosynaptic currents from multiple different cell types within the same experiment. Since the combination of presynaptic and postsynaptic cell types are known ahead of time, in practice we are free to select an NWD network trained to demix synaptic currents of a specific type. However, we note that in future work one could adapt the NWD network to account for multiple cell types simultaneously.

Further, the timescale and variance of the temporally-correlated noise *g*_*k*_ can be adjusted to match experimental conditions. For example, in the context of *in vivo* experiments it may be critical to account for e.g. neuromodulatory signals, which could be approximated by sampling from Gaussian processes with a range of timescales and amplitudes. Similarly, the uncorrelated noise *ϵ*_*k*_ can be tuned to match the electrical noise from the recording electrode.

The NWD network intrinsically performs a denoising of the evoked PSCs since each target PSC in the training data was uncorrupted by the noise terms *g*_*k*_ and *ϵ*_*k*_. We found that we could obtain further improvement in PSC denoising by randomly incorporating 45 ms snippets of pure electrical noise taken from experiments during periods without stimulation. For such “negative” templates, the network was trained to produce an output of zeros.

Finally, we performed a minor correction step to guarantee that the output of the NWD network monotonically decayed after a predefined time [Lee et al., 2020], which we found marginally improved accuracy with little computational cost. Namely, after a user-specified time *t*_monotone_, the network output *N*_*t*_ for all *t > t*_monotone_ was adjusted by recursively setting *N*_*t*_ = min{*N*_*t*_, *N*_*t*−1_}.

We implemented NWD using PyTorch Lightning. Networks were trained using stochastic gradient descent with a batch size of 64 and learning rate of 0.01 for 3000-6000 epochs, which was sufficiently long for the optimizer to converge in our training regime.

### Statistical model

CAVIaR aims to fit a statistical model to the demixed optogenetic data that relates patterns of holographic stimulation to the resulting measurements of postsynaptic current. Let **c**_*k*_ ∈ ℝ^*T*^ represent the PSC trace on trial *k*. In accordance with the trial structure used for NWD, we assume that *T* = 900 frames (or 45 ms), with photostimulation beginning at *t* = 100 (or 5 ms). To make use of compressed sensing-style techniques, we collapse the demixed, denoised PSC trace into a single number by integrating over the duration of the trial,

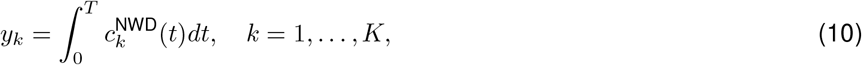

where 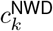 represents the demixed trace on trial *k*. The measurements *y*_*k*_ therefore represent the total synaptic charge transfer resulting from the transmission of presynaptic spikes. Assuming the postsynaptic neuron is held under voltage clamp and that space clamp imperfections do not induce substantive nonlinearities, we may model these measurements as a simple sum of optogenetically-evoked presynaptic spikes weighted by the strengths of the corresponding synapses,

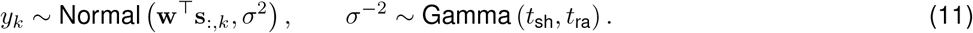

Here **w** ∈ ℝ^*N*^ is a vector of synaptic weights encoding the charge transfer resulting from a single spike for each neuron *n* = 1, … *N*. Because opsin is expressed in a specific presynaptic cell type (in this study exactly one of SST, PV, or pyramidal), there is no risk of potentially-confounding “cancellation” effects where both excitatory and inhibitory PSCs are elicited and sum to zero.

Next, **s**_:,*k*_ is the vector of presynaptic spikes on trial *k*. Note that, as described below, spike generation is highly stochastic, and hence *s*_*nk*_ is a latent variable. The synaptic weights are regularized by imposing a Gaussian prior,

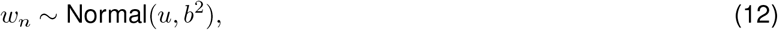

with hyperparameters *u* and *b*^2^, and the variance *σ*^2^ is intended to account for electrical noise from the recording electrode and variability in the amplitude of the evoked PSCs. Model parameters are listed in Table 1.

**Table 1:** Table of model parameters and their interpretations.

| Symbol | Interpretation | Hyperparameters | Posterior |
| --- | --- | --- | --- |
| $\mathbf{w}$ | Connection strengths | $u, b^2$ | $\boldsymbol{\mu}, \boldsymbol{\Omega}$ |
| $s_{nk}$ | Presynaptic spike | — | $\lambda_{nk}$ |
| $\phi_n^0, \phi_n^1$ | Presynaptic sigmoid coefficients | $\mathbf{v}, \mathbf{L}$ | $\boldsymbol{\nu}_n, \boldsymbol{\Sigma}_n$ |
| $\sigma^2$ | Observation noise variance | $t_{\text{sh}}, t_{\text{ra}}$ | $\theta_{\text{sh}}, \theta_{\text{ra}}$ |
| $z_k$ | Spontaneous synaptic current | | |
| $y_k$ | Observed response (integral of the PSC) | | |
| $I_{nk}$ | Laser power applied to neuron $n$ on trial $k$ | | |
| $N(n)$ | Number of neurons (neuron index) | | |
| $K(k)$ | Number of trials (trial index) | | |
| $f$ | Sigmoid function | | |

The variability of spike generation in our experimental data shows that the probability of generating a spike is not only different between cells, but also across power levels, and is therefore effectively impossible to know ahead of time. We propose a structured presynaptic model that can flexibly adjust to varying spike probabilities on a cell-by-cell basis. In particular, presynaptic spikes are assumed to be driven by holographic photostimulation according to a linear-nonlinear-Bernoulli model,

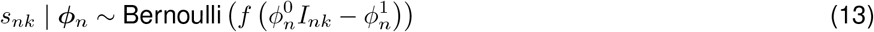

where *f* (*x*) = 1*/*(1 + exp(−*x*)) is the sigmoid nonlinearity, *I*_*nk*_ denotes the power of the stimulation laser focused on cell *n* on trial *k*, and the sigmoid coefficients follow bivariate normal distributions

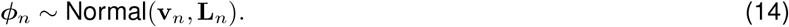

During inference we thus effectively perform a Bayesian logistic regression of each cell’s inferred presynaptic spikes on the laser targets. This presynaptic model allows spike probabilities to adapt to variation in opsin expression via the sigmoid coefficient term 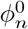, as well as to variation in rheobase via the intercept term 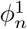.

### Posterior inference

#### Variational approximation

The goal of CAVIaR is to infer the posterior distribution over the hidden variables, including the synaptic weights and presynaptic spikes. The posterior describes the distribution of parameter values that are consistent with the observed data, given our prior assumptions. Importantly, this provides a description of uncertainty in the inferred synaptic weights. It is not computationally feasible to compute the exact posterior distribution, but standard methods have been developed to approximate the posterior [Blei et al., 2017]. Here we modify the conventional coordinate-ascent variational inference algorithm by augmenting it with several steps that account for spontaneous synaptic currents and biophysical plausibility of the inferred spikes. To this end, we note that the posterior distribution over the latent variables factorizes as

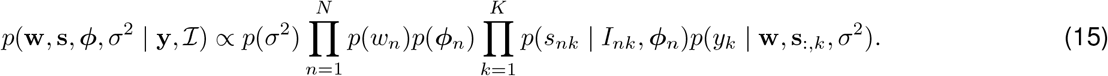

We approximate the posterior distribution *p*(Ƶ | **y**, *ℐ*) by a variational model *q*(Ƶ) (where Ƶ represents the set of latent variables) that obeys a similar factorization,

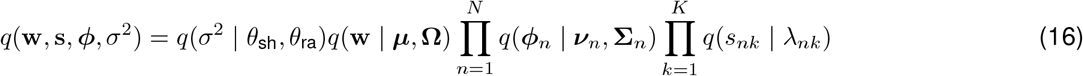

where the individual factors are *q*(**w** | ***µ*, Ω**) = Normal(**w** | ***µ*, Ω**), *q*(*s*_*nk*_ | *λ*_*nk*_) = Bernoulli(*s*_*nk*_ | *λ*_*nk*_), *q*(***ϕ***_*n*_ | ***ν***_*n*_, **Λ**_*n*_) = Normal(***ϕ***_*n*_ | ***ν***_*n*_, **Σ**_*n*_), and *q*(*σ*^−2^ | *θ*_sh_, *θ*_ra_) = Gamma(*σ*^−2^ | *θ*_sh_, *θ*_ra_).

Given the above approximation, coordinate-ascent variational inference seeks to perform an update of each factor *q*(Ƶ _*i*_) one-by-one for all Ƶ_*i*_ ∈ Ƶ, with the idea being that each update moves the approximate posterior *q*(Ƶ) closer to the true posterior *p*(Ƶ| **y**, *ℐ*) in the sense of the Kullback-Leibler divergence (KL-divergence) [Blei et al., 2017]. We take a variety of approaches to updating each factor depending on the relative tractability of the update. The complete algorithm is given in Algorithm 1.

#### Inference of synaptic weights

First, it can be shown that the optimal variational update for the synaptic strengths **w**, conditional on all other latent variables Ƶ \ **w**, obeys the equation

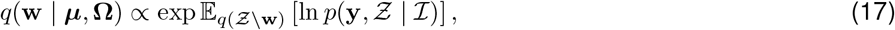

see, e.g., [Bishop, 2006]. Since both the prior on **w** and the observations are Gaussian-distributed, one can evaluate Equation 17 analytically and solve for ***µ*** and **Ω** by “completing the square”, yielding the block update

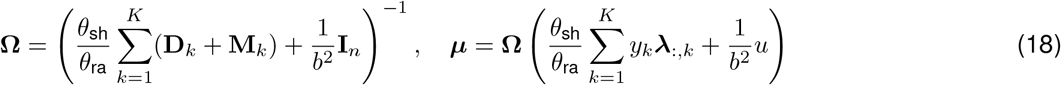

where **D**_*k*_ = diag (*λ*_1,*k*_(1 − *λ*_1,*k*_), …, *λ*_*Nk*_(1 − *λ*_*Nk*_)) ∈ R^*N×N*^ is diagonal and 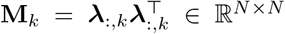. Inspecting the update for ***µ***, one sees that a neuron’s synaptic weight is only determined by the magnitude of the postsynaptic responses on trials for which that neuron actually elicited a spike. However, those spikes are not directly observed, and must themselves be inferred from the data.

#### Inference of presynaptic spikes

Given a measurement *y*_*k*_ resulting from the photostimulus *I*_:,*k*_ and our current estimates of synaptic strengths, electrical noise, and *a priori* presynaptic spike probabilities, we must decide which of the targeted cells actually elicited a spike. In early trials, this depends more on the prior (i.e., the presynaptic model) rather than on the likelihood, and this balance shifts in favor of the likelihood the more experimental data that is collected. However, unlike in Equation 17, the optimal variational update for the presynaptic spike probabilities *λ*_*nk*_ cannot be solved for analytically. Instead, one can recognize ([Bishop, 2006]) that the KL-divergence from the approximate to the true posterior, KL(*q*∥*p*), can be decomposed into the sum

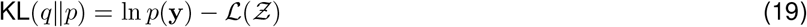

where ln *p*(**y**) = ln ∫ *p*(**y** *Ƶ*)*d* Ƶ is the logarithmic model evidence and where

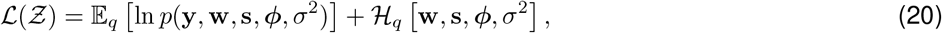

with ℋ _*q*_[*x*] the Shannon entropy of variable *x* under density *q*. Since the KL-divergence is always non-negative, the inequality ℒ (Ƶ) ≤ ln *p*(**y**) always holds, and therefore ℒ (Ƶ) is a lower bound on the logarithmic model evidence (referred to as the “evidence lower bound”). By maximizing the evidence lower bound as a function of the variational model parameters, ℒ (Ƶ) approaches ln *p*(**y**), and hence drives KL(*q*∥*p*) towards zero. If the bound is tight, i.e., if ℒ (Ƶ) = ln *p*(**y**), the KL-divergence is zero and the variational approximation to the posterior becomes exact. To this end, differentiating ℒ and solving for *λ*_*nk*_ yields the update

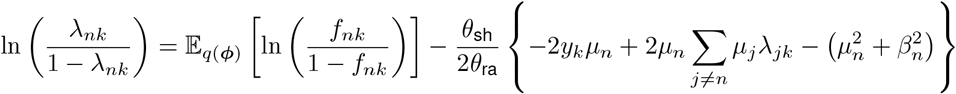

where we have defined 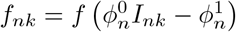. The expectation in the above term is analytically intractable due to multiple nonlinearities. We thus make a Monte Carlo approximation to the expectation

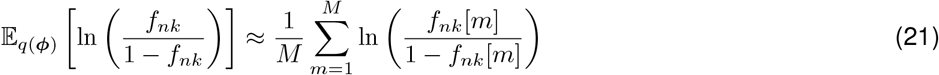

With 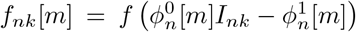, and for each pair 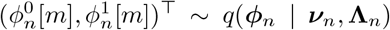. As discussed below, the sigmoid coefficients are sampled from a *truncated* multivariate normal distribution. However, sampling from truncated multivariate normal distributions is computationally demanding and remains an open area of research [Pakman and Paninski, 2014]. Instead, the terms 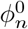 and 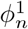 are sampled independently from their truncated marginal distributions.

Inspecting the updates for *λ*_*nk*_, one notices that each update depends on all other neurons in the population, but is independent of all other trials. Thus, we update the spike probabilities for all trials simultaneously, but perform this one neuron at a time (where the order of the neuron updates is randomized every iteration). Immediately following the inference of a neuron’s spikes, we evaluate their biophysical plausibility as a function of laser power. Patch-clamp studies with ChroME2 opsins show that the probability of eliciting an action potential increase monotonically with laser power [Sridharan et al., 2022]. We therefore perform an isotonic regression through the inferred spike rate using the pool adjacent violators algorithm (PAVA) [Ayer et al., 1955].

Concretely, let 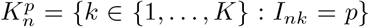 be the set of all trials on which neuron *n* was stimulated with power *p*, and let

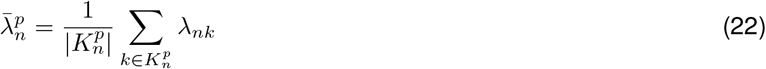

denote the average inferred spike probability for neuron *n* at power *p*. PAVA computes an isotonic regression function 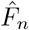 by solving the optimization problem

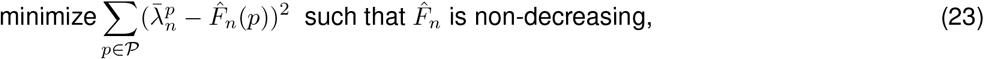

where 𝒫 is the set of non-zero powers used in the experiment. This provides us with an isotonically-constrained optogenetic “power curve” 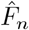 that we use to judge whether or not the putative synapse connecting neuron *n* to the patched neuron is sufficiently plausible. In particular, we require that at the maximal power used in the experiment the inferred probability of successfully initiating and transmitting a spike is greater than some threshold *θ*_PAVA_ ∈ [0, 1], which we refer to as the “minimum spike rate at maximum power” (MSRMP). However, some fraction of the inferred spikes may result from spontaneous synaptic currents, which can lead to false positives if left unaccounted for. We therefore infer the rate of spontaneous currents (*λ*_spont_, described below) and add this to the user-defined threshold *θ*_PAVA_ to adaptively adjust the plausibility criterion. Neurons with 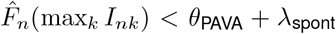 (i.e., whose adjusted maximal spike rate is below *θ*_PAVA_) are considered unconnected and have their synaptic weights and inferred spike probabilities set to zero. We used *θ*_PAVA_ = 0.4 for mapping inhibitory presynaptic neurons, and *θ*_PAVA_ = 0.3 for mapping excitatory presynaptic neurons.

Note that the likelihood of spurious connections arising from spontaneous PSCs only at the highest laser power is vanishingly small (Supplementary note 1).

We also use a “masking” procedure to prevent small, noise-driven postsynaptic measurements from causing spurious spike inferences. To do this, we evaluate a test statistic *τ* on each trace *c*_*k*_. If *τ* (*c*_*k*_) *< τ*_min_ we force *λ*_*nk*_ = 0 for all *n* = 1, …, *N*. The statistic *τ* is typically chosen to be the sample autocorrelation.

#### Inference of presynaptic sigmoid coefficients

The presynaptic sigmoid coefficients ***ϕ*** (governing the presynaptic models) are updated using Laplace’s method. Namely, rather than solving for the optimal variational approximation or optimizing the evidence lower bound, we learn a posterior distribution over each ***ϕ***_*n*_ by making a second-order Taylor approximation about the posterior mode, with the posterior covariance matrix given by the inverse of the Hessian matrix appearing in the Taylor series. In some cases we encountered pathological spike inference when the sigmoid coefficients ***ϕ***_*n*_ became negative, and hence we enforce a positivity constraint in the mode-finding algorithm described below. Moreover, the Monte Carlo approximation described in Equation 21 depends closely on the non-negativity of the sampled variates. Thus we truncate the Laplace approximation to the non-negative real line,

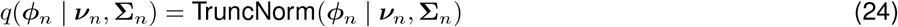

with 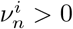 for *i* ∈ {0, 1} and supp 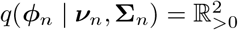.

Note that, while PAVA is a non-parametric estimator of the spike rate, its use differs from the presynaptic spike model which takes a parametric form. The presynaptic model is used to evaluate the prior probability of eliciting a spike, whereas PAVA is used to evaluate the plausibility of the inferred spikes.

We use Newton’s method with a log barrier and backtracking line search to identify the posterior mode ***ν***_*n*_. To this end, define the objective function with barrier penalty *t* as

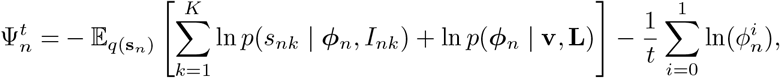

where we average over the spike uncertainty. Note that the presynaptic models depend only on the spikes **s**, and are independent conditional on **s**, allowing us to compute Laplace approximations in parallel across cells.

The mode-finding algorithm starts with an initial *t*. We then solve for the mode by iterating the Newton steps

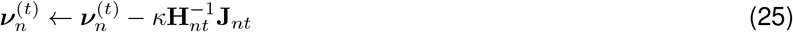

until convergence, where **J** and **H** are respectively the Jacobian and Hessian of the presynaptic model for neuron *n* with barrier sharpness *t*

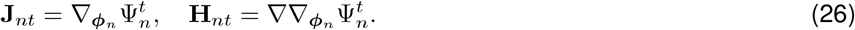

The stepsize *κ* is adaptively selected using a standard backtracking line search rule. We then increase *t* to sharpen the log barrier and repeat as required. Typically we only sharpen *t* two or three times as this proved sufficient for our data.

#### Inference of observation noise variance

To infer the observation noise variance, we use the same approach as with the update rule for the synaptic weights (Equation 17). In particular, noting that

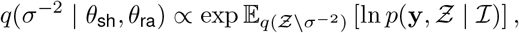

and recognizing that the gamma prior is conjugate to the Gaussian likelihood, one obtains the variational update

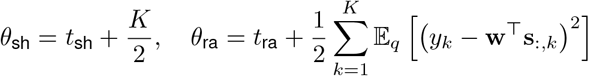

where the expectation above can be evaluated analytically, though we leave it in the above form for legibility.

#### Inference of spontaneous synaptic currents

Finally, we estimate spontaneous synaptic currents *z*_*k*_ using soft-thresholding. The idea is that, given a numerical tolerance *ϵ* (where typically *ϵ* = 0.05) and positively rectified residuals

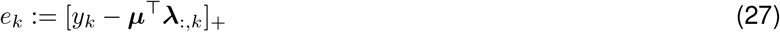

we apply a soft-thresholding function *S* with penalty *γ* defined by

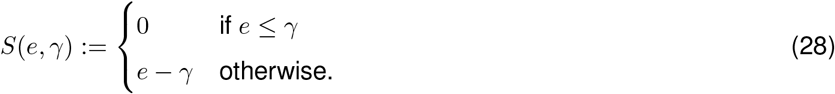

We then iteratively shrink the penalty *γ* until the norm of the residual data comprises no more than *ϵ* of the norm of the observed data; i.e., until

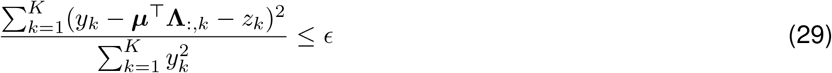

where *z*_*k*_ is obtained via the soft-thresholded residuals. However, we also require that (*z*_1_, …, *z*_*K*_) be approximately orthogonal to ***λ***_*n*_ for all *n*, and apply the masking procedure as noted in the spike inference step above, leading to the spontaneous synaptic current estimator

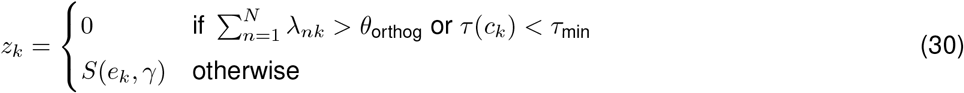

where *θ*_orthog_ ≈ 0, but is not exactly zero to allow for numerical imprecision. The spontaneous events are constrained to be approximately orthogonal to the inferred spikes to prevent variability in PSC amplitude causing false positives.

### Post-hoc scan for false negatives

Occasionally the CAVIaR algorithm will erroneously declare a neuron to be disconnected, either due to a failure to meet the PAVA-based plausibility criterion in the earliest iterations of the algorithm, or due to the biological data violating the assumptions of the underlying statistical model. To correct for this, following inference we perform a post-hoc scan for potential false negatives. The idea is to reconnect neurons that CAVIaR originally declared to be disconnected by checking whether spontaneous PSCs coinciding with the stimulation of a selected neuron could constitute valid postsynaptic responses when the rigidity of the statistical model is relaxed. Since the overwhelming bulk of connections are already identified by the first CAVIaR pass, missed connections are comparatively rare and thus we found a simple greedy algorithm to be effective.

The false-negative scanning algorithm is given in Algorithm 3. Briefly, the algorithm begins by collecting all neurons *S*_disc_ declared to be disconnected by the first CAVIaR pass. It then selects the neuron *n*^*^ ∈ *S*_disc_ with the greatest number of coincidental spontaneous PSCs, and checks if assigning these PSCs to neuron *n*^*^ would satisfy the PAVA criterion. If so, neuron *n*^*^ is declared connected, with the posterior distribution of the reconnected neuron’s parameters determined by sample statistics of the corresponding spontaneous PSCs. Neuron *n*^*^ is then removed from *S*_disc_ (whether reconnected or not), and the algorithm repeats until *S*_disc_ = ∅.

### Inference of canonical postsynaptic current waveforms

Once we have inferred the presynaptic spike matrix **Λ**, we can obtain accurate PSC waveforms **r**_*n*_ ∈ ℝ^*T*^ using ridge regression. Collecting the waveforms in the rows of a matrix **R**, they can be obtained simultaneously by solving the non-negative *L*_2_ problem

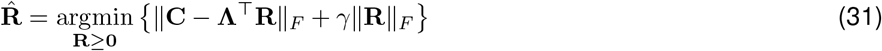

where **C** ∈ ℝ^*K×T*^ is the matrix of PSC traces (**c**_1_, …, **c**_*K*_), *γ >* 0 is the ridge penalty, and ∥ ·∥_*F*_ is the Frobenius norm. Note that by using the spike matrix **Λ** instead of the optogenetic “design matrix” (where each element of the matrix determines which neurons were merely stimulated as in [Hu and Chklovskii, 2009], rather than which spiked as a result of photostimulation), the estimated waveforms are much less biased by trials in which neurons were not photoactivated. We used this ridge regression approach to determine synaptic weights from PSCs throughout the paper.

#### Algorithm 1: Coordinate-ascent variational inference and isotonic regularization (CAVIaR)

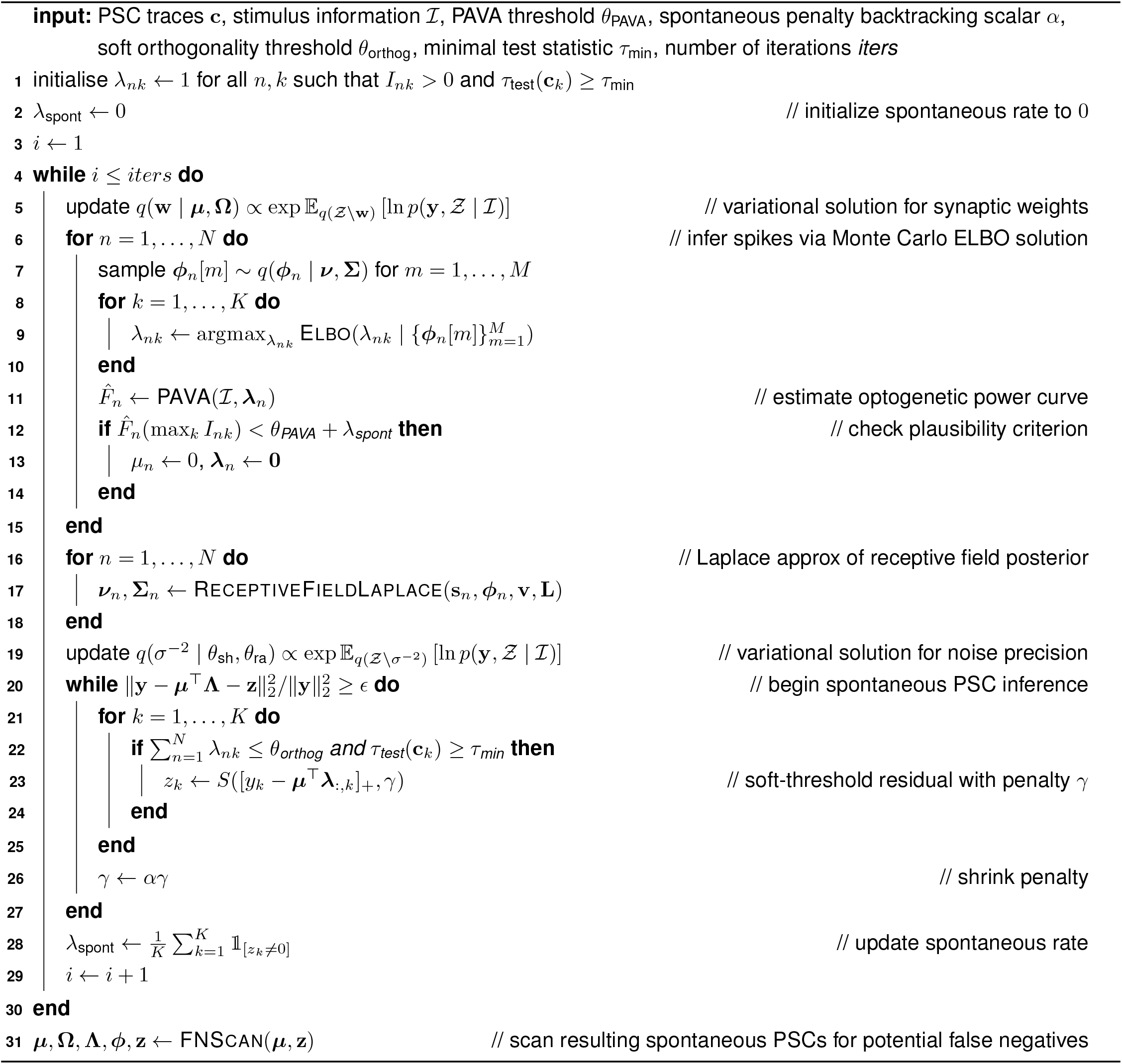

#### Algorithm 2: RECEPTIVEFIELDLAPLACE

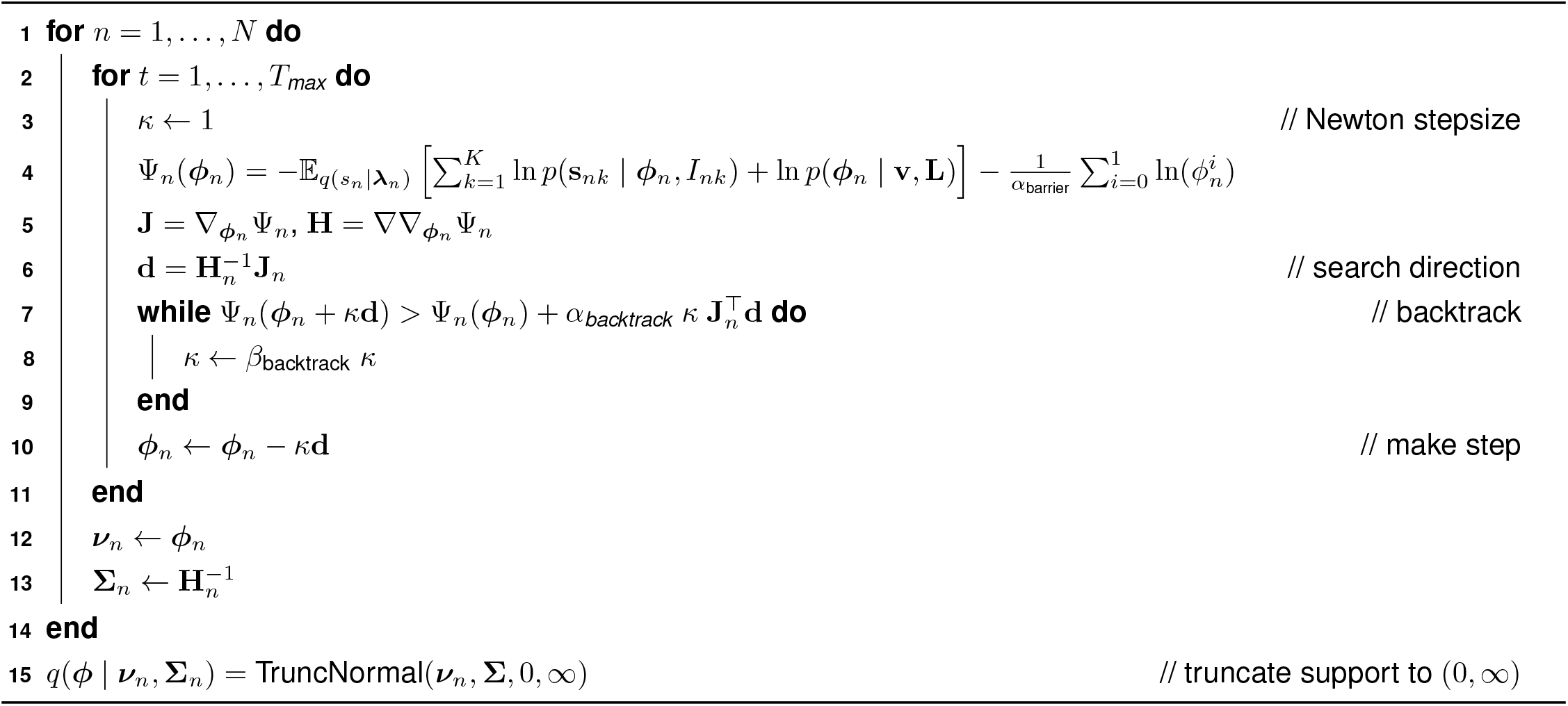

#### Algorithm 3: FNSCAN

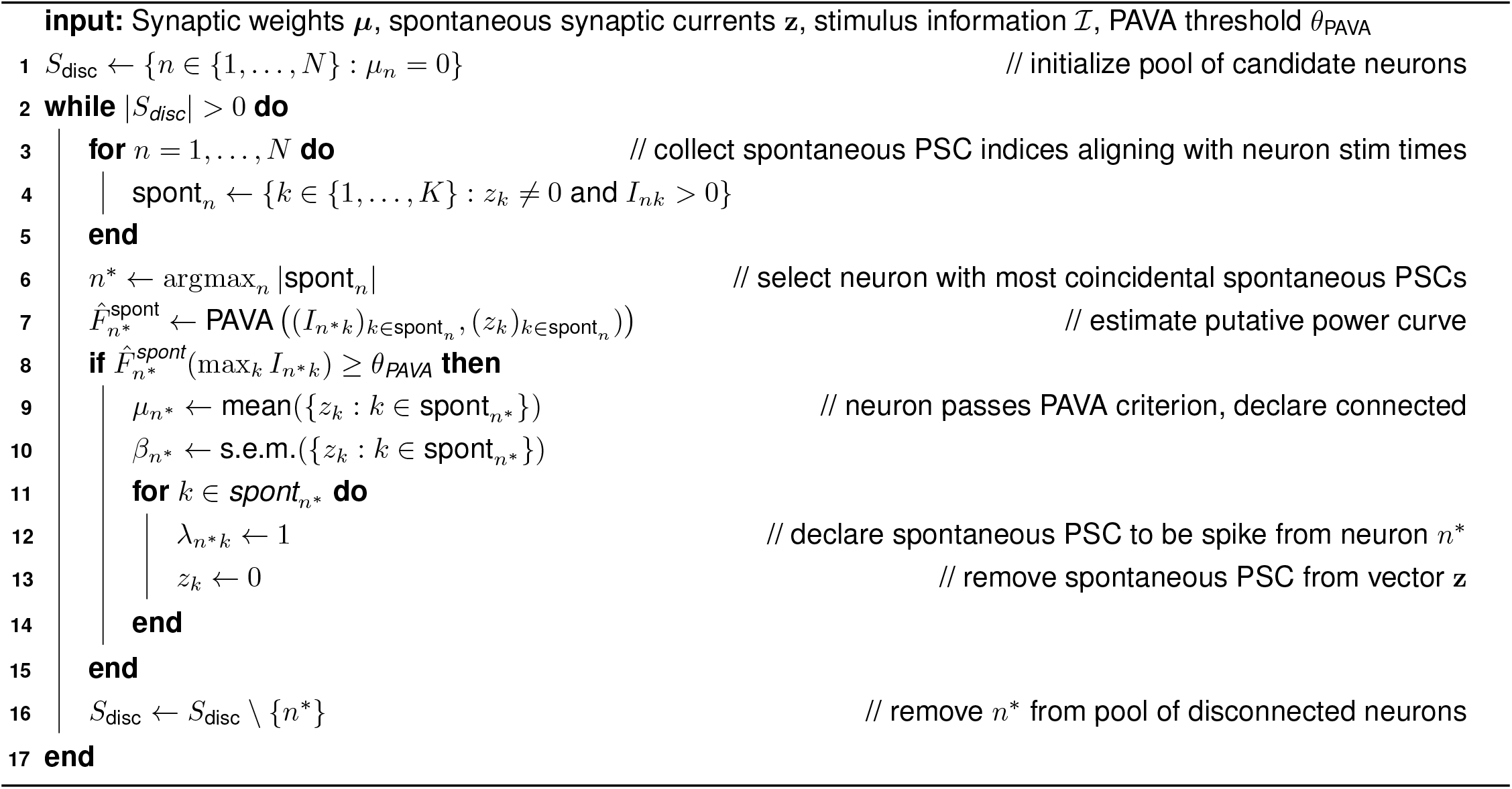

### Comparing estimated connectivity vectors

To quantify the similarity between connectivity vectors obtained using single-target stimulation and ensemble stimulation, we used the coefficient of determination (R^2^, computed using the scikit-learn Python package) as well as the precision and recall, defined as

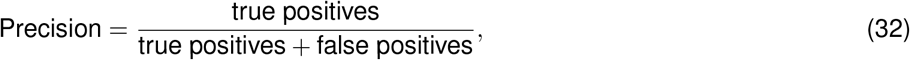

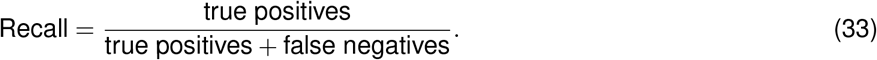

Intuitively, the precision represents the fraction of connections found by compressed sensing (CAVIaR) that are “true”, and the recall represents the fraction of “true” connections that are correctly found by compressed sensing. Since we do not have access to the ground truth connectivity in real experiments, we treat the connections identified using single-target stimulation as the target for compressed sensing.

### Leave-one-hologram-out cross-validation

We assess the accuracy of the CAVIaR inferences using leave-one-hologram-out cross-validation (LOHO-CV, Algorithm 4). Let H represent the complete set of hologram designs; i.e., *h* ∈ H determines which neurons will be targeted for stimulation, regardless of laser power. LOHO-CV proceeds by selecting *h* ∈ H, fitting CAVIaR to the PSC traces and stimuli corresponding to all other holograms H\{*h*}, and then averaging samples from the posterior predictive distribution over responses to hologram *h* to obtain an estimate of the postsynaptic response (at each power level).

#### Algorithm 4: LOHO-CV

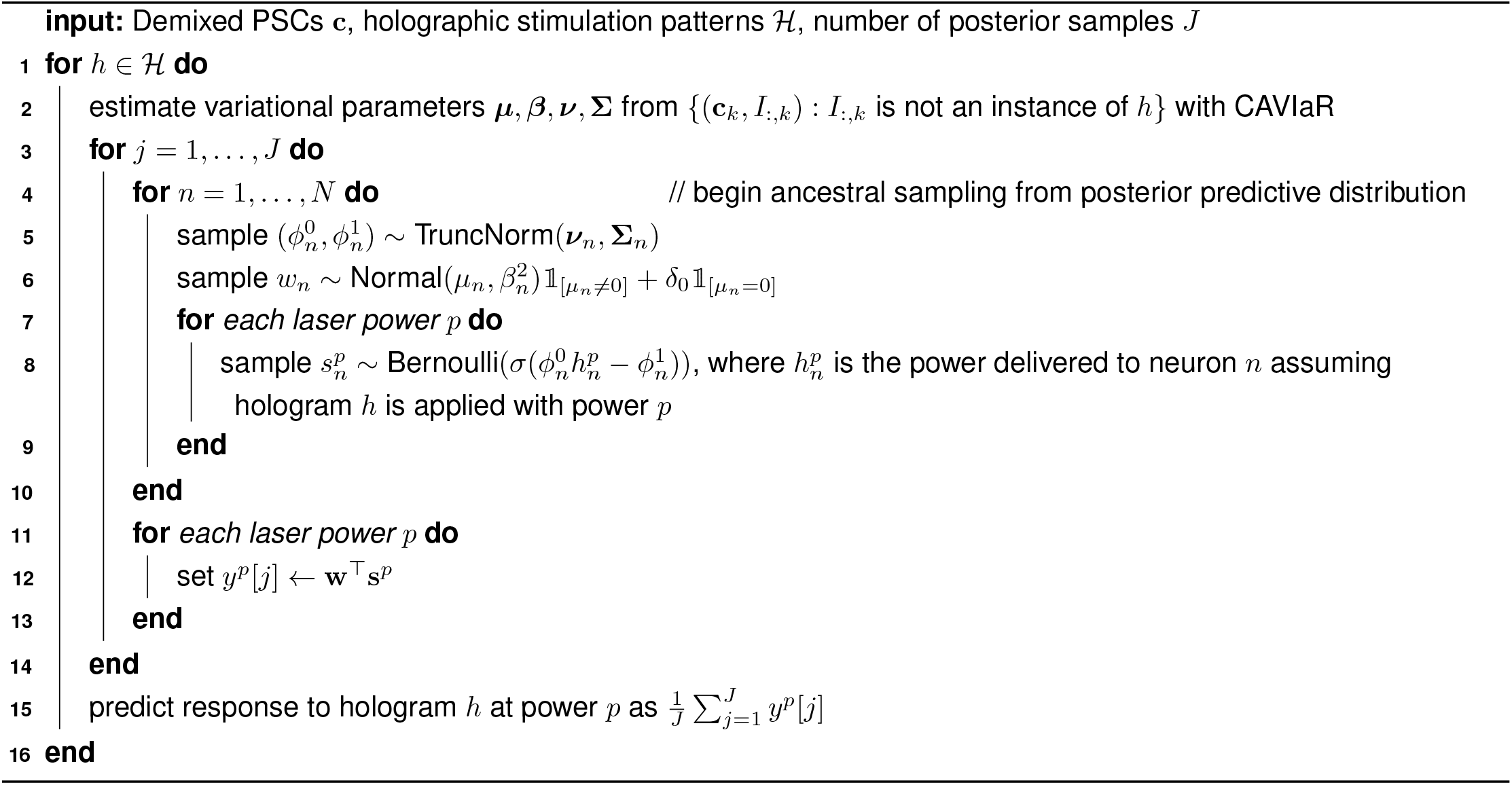

### Simulated circuit mapping experiments

We used simulated data to characterize the performance of the techniques we tested. To ensure the accuracy of this characterization, rather than sampling directly from the generative model that we propose, we added several layers of biophysical realism. In particular, we sampled noisy synaptic currents that were first demixed using NWD before being supplied as input to the connectivity inference algorithms. This way we could test for the combined accuracy of NWD and connectivity inference.

The simulated data was generated as follows. First, sigmoid parameters 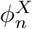 were sampled uniformly from 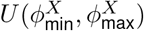 for *X* ∈ {0, 1}. Then, presynaptic spikes *s*_*nk*_ were sampled from a linear-nonlinear-Bernoulli model

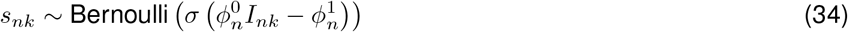

where the laser power on each trial was randomly selected from a discrete set matched to the experimental data. Each neuron had a canonical PSC transient 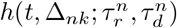 that took a form similar to that used for training the NWD networks. The unnormalized transient 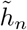 was defined by

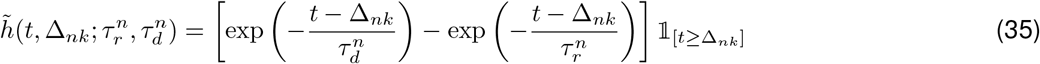

Where 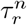 and 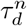 are rise and decay time constants and Δ_*nk*_ represents the spike delay (i.e., the combined spike activation and transmission latencies) for neuron *n* on trial *k*. The transient was then normalized to take integral 1,

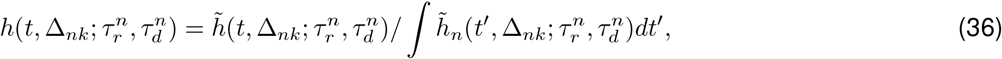

such that the synaptic weight for that neuron (*w*_*n*_, defined as the synaptic charge transfer below) was preserved when multiplying the weight by the PSC.

The spike delays were laser power-dependent so that, in accordance with experimental data for the opsins used in this study [Sridharan et al., 2022], neurons initiated and propagated spikes faster if stimulated at higher powers. Concretely, for laser power *I*_*nk*_ targeted at neuron *n* on trial *k*, the spike time was sampled from a right-shifted gamma distribution,

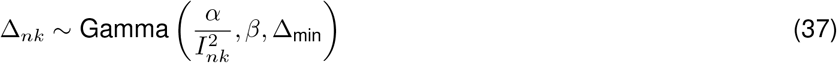

Where

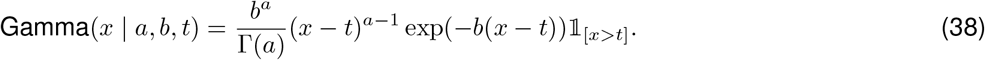

Note that since the mean of the shifted gamma distribution is *t* + *a/b*, the optogenetically evoked spikes followed – in expectation – an inverse-square dependence of time on power, consistent with the physics of two-photon absorption [Papagiakoumou et al., 2020]:

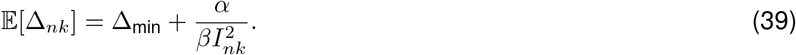

The synaptic weights *w*_*n*_ are intended to model the total synaptic charge transfer resulting from the transmission of a presynaptic spike to the postsynaptic neuron. We sampled the weights in a way that reflects the typical observations from our experiments. Namely, of the neurons that were chosen to be synaptically connected, a small number of them were strongly connected (in our simulations this was 20% of the connected population, though the precise value did not notably impact our results) and a large number of them were more weakly connected (80%). Specifically, for a given connectivity rate *α* ∈ (0, 1), a subset of ⌈*αN* ⌉ neurons were randomly selected as being connected to the postsynaptic neuron. Then, if a neuron *n* was chosen to be strongly connected, its weights were sampled as

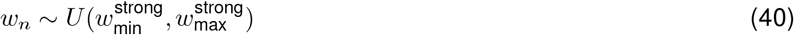

where 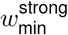 and 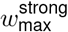 respectively represent the lower and upper bounds of the uniform distribution. If *n* was weakly connected then its weights were sampled as

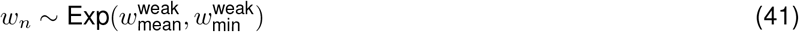

where the exponential distribution above is in its two-parameter, right-shifted form

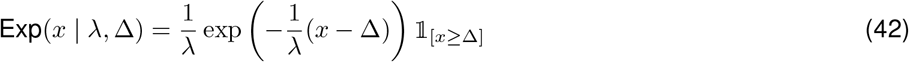

with Δ ≥ 0.

We used two approaches to generating PSC traces. Either we directly simulated individual 45 ms trials for each *k* = 1, …, *K*, or we simulated continuous circuit mapping experiments at 20 kHz sampling resolution (matched to the experimental data) that lasted for tens of minutes. Using the latter approach we could very closely mimic the exact contribution of confounding synaptic currents arising from stimulation at very high frequencies. Simulations performed in this continuous manner were subsequently restructured into the usual 45 ms snippets of activity.

In the first “trial-wise” approach to simulating data, the presynaptic spikes, synaptic weights, and PSC kernels were used to generate the postsynaptic responses *y*_*k*_ as

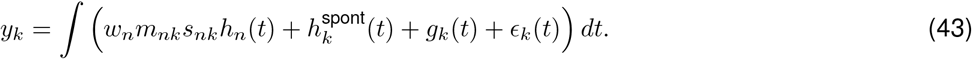

Here *m*_*nk*_ is a multiplicative noise term, 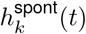 is a spontaneous PSC term, *g*_*k*_(*t*) is a temporally correlated noise term, and *ϵ*_*k*_(*t*) is an additive noise term.

The multiplicative noise term *m*_*nk*_ accounted for the fact that, in our experimental data, the precise amplitudes of PSC transients were observed to vary from trial to trial. We sampled *m*_*nk*_ from a log-normal distribution,

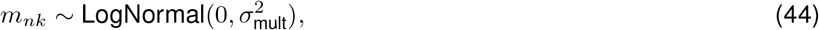

such that the median postsynaptic charge transfer following a presynaptic spike still took the value *w*_*n*_. The spontaneous term 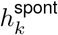 was either a PSC as in Equation 36 with random time constants, amplitudes, and spike times, or the zero vector, depending on the probability of spontaneous events. Temporally correlated noise, **g**_*k*_ ∈ ℝ^*T*^, was sampled from a Gaussian process,

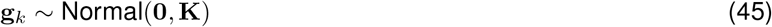

where **K** was defined by the radial basis function kernel,

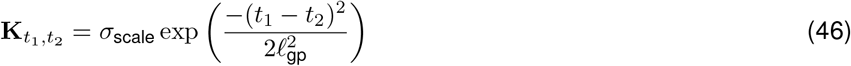

for *t*_1_, *t*_2_ = 1, …, *T*. Finally, the additive noise *ϵ*_*k*_(*t*) was sampled independently and identically from a zero-mean Gaussian, *ϵ*_*k*_(*t*) ∼ Normal(0,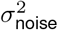). Due to high computational tractability this trial-wise approach was used to generate the heatmaps in Figure 3.

The second “continuous experiment” approach to simulation was much more computationally demanding. Simulating a 30 minute experiment at 20 KHz requires 36,000,000 time points, making, for example, the use of Gaussian processes prohibitive. Our approach was to generate vectors encoding spike times for each neuron and convolve this with the corresponding neuron’s PSC kernel while batching over blocks of time. To this end, an experiment of length *T* was evenly subdivided into trials according to the stimulation frequency *f*. For each such trial, we sampled spikes, spike latencies, and multiplicative noise for each neuron as described above. Then, using this collection of variables, we defined spike vectors ***ζ***_*n*_ ∈ ℝ^*T*^ for *n* = 1, …, *N* by setting

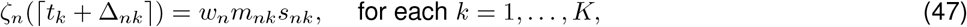

where *t*_*k*_ is the timebin at which the *k*th trial begins, and *ζ*_*n*_(*t*) = 0 at all other timebins. Similarly, we generated spon-aneous spike vectors 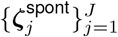 at a specified rate *λ*_spont_ (in Hz), where each 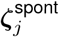 encodes the activation of a single spontaneous PSC with waveforms 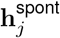 defined by randomly sampled time constants, amplitudes, and onset times. The full-length postsynaptic measurement vector was then obtained by convolving the spike vectors with the PSC waveforms and summing across neurons and spontaneous inputs,

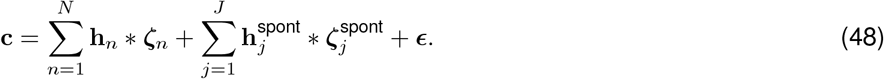

Here the noise follows a first-order autoregressive process

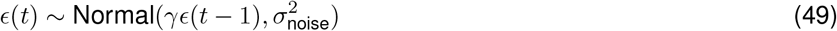

with autoregressive coefficient *γ* ∈ (0, 1), which was suitably scalable for continuous simulated experiments.

The postsynaptic responses *y*_*k*_ used by the connectivity inference algorithms were extracted from **c** by setting

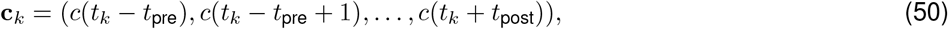

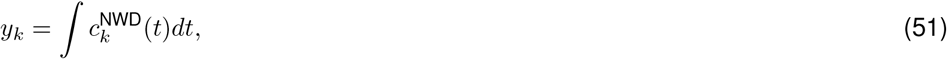

where *t*_*k*_ represents the beginning of the *k*-th trial, *t*_pre_ = 100 and *t*_post_ = 800, and 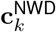 represents the trace **c**_*k*_ after demixing. Note that as the rate of stimulation increases, the interstimulus-interval decreases, and the trial windows increasingly overlap in time. This leads to a confounding of the observed PSCs as in Figure 2, which the NWD network must suppress.

### Combinatorics of expected interstimulus intervals

Let *N* represent the total number of potential presynaptic neurons, *R* the size of the ensemble, and *f* the rate of stimulation (in Hz). Assuming neurons are chosen uniformly at random, the probability of selecting neuron *n* ∈ {1, …, *N*} is

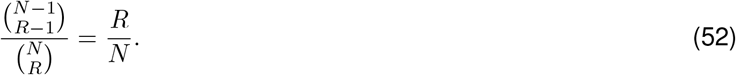

Hence, one must stimulate *N/R* times on average to return to the same neuron. If stimulation occurs at *f* Hz, this takes *N/Rf* seconds.

## Supplementary material

**Table S1:**
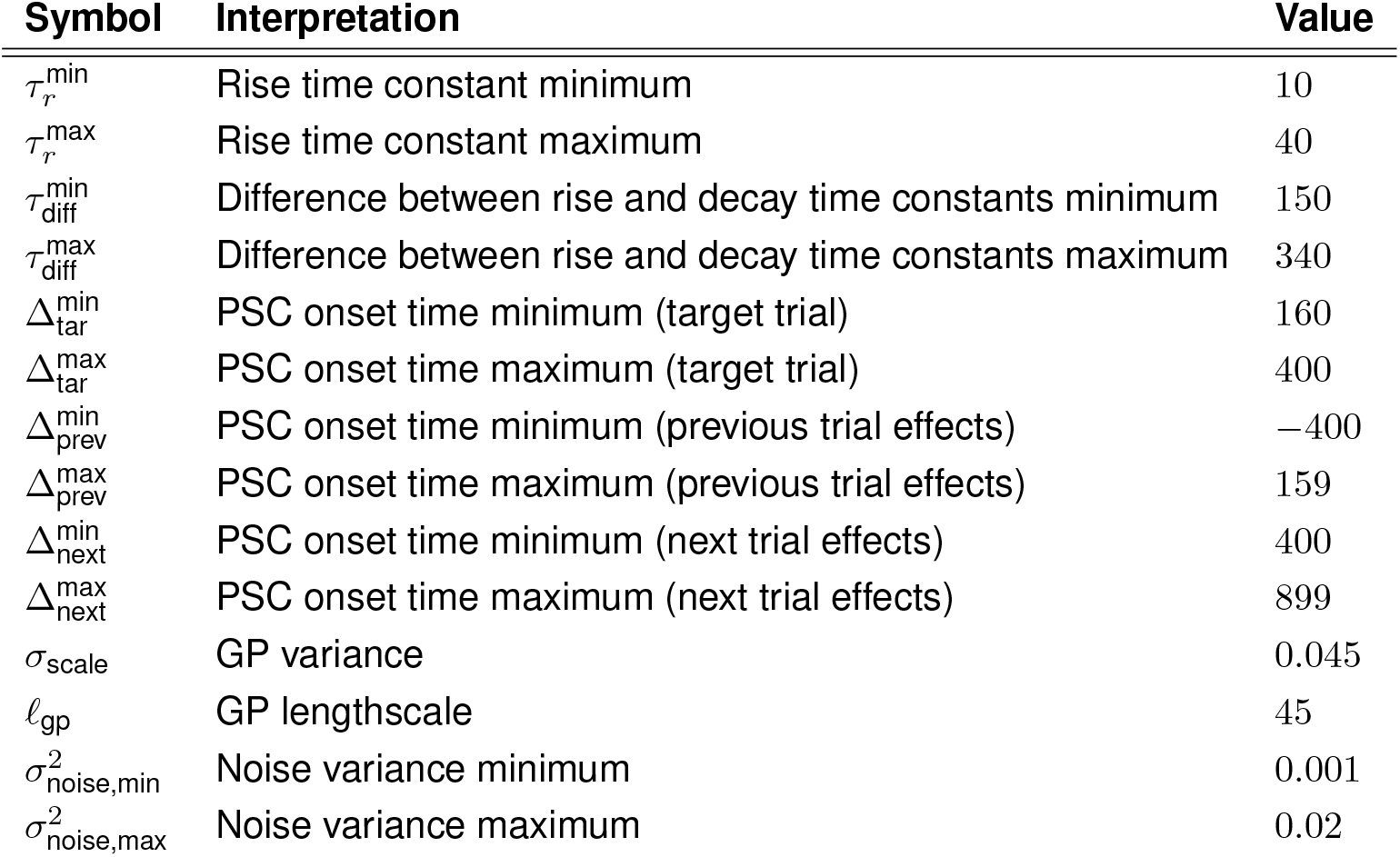
Parameters used for training an NWD network for inhibitory-to-excitatory mapping experiments. Assumes timesteps measured at 20 kHz.

| Symbol | Interpretation | Value |
| --- | --- | --- |
| $\tau_r^{\min}$ | Rise time constant minimum | 10 |
| $\tau_r^{\max}$ | Rise time constant maximum | 40 |
| $\tau_{\text{diff}}^{\min}$ | Difference between rise and decay time constants minimum | 150 |
| $\tau_{\text{diff}}^{\max}$ | Difference between rise and decay time constants maximum | 340 |
| $\Delta_{\text{tar}}^{\min}$ | PSC onset time minimum (target trial) | 160 |
| $\Delta_{\text{tar}}^{\max}$ | PSC onset time maximum (target trial) | 400 |
| $\Delta_{\text{prev}}^{\min}$ | PSC onset time minimum (previous trial effects) | -400 |
| $\Delta_{\text{prev}}^{\max}$ | PSC onset time maximum (previous trial effects) | 159 |
| $\Delta_{\text{next}}^{\min}$ | PSC onset time minimum (next trial effects) | 400 |
| $\Delta_{\text{next}}^{\max}$ | PSC onset time maximum (next trial effects) | 899 |
| $\sigma_{\text{scale}}$ | GP variance | 0.045 |
| $\ell_{\text{gp}}$ | GP lengthscale | 45 |
| $\sigma_{\text{noise,min}}^2$ | Noise variance minimum | 0.001 |
| $\sigma_{\text{noise,max}}^2$ | Noise variance maximum | 0.02 |

**Table S2:** Parameters used for training an NWD network for excitatory-to-excitatory mapping experiments. Assumes timesteps measured at 20 kHz.

| Symbol | Interpretation | Value |
| --- | --- | --- |
| $\tau_r^{\min}$ | Rise time constant minimum | 10 |
| $\tau_r^{\max}$ | Rise time constant maximum | 40 |
| $\tau_{\text{diff}}^{\min}$ | Difference between rise and decay time constants minimum | 60 |
| $\tau_{\text{diff}}^{\max}$ | Difference between rise and decay time constants maximum | 120 |
| $\Delta_{\text{tar}}^{\min}$ | PSC onset time minimum (target trial) | 160 |
| $\Delta_{\text{tar}}^{\max}$ | PSC onset time maximum (target trial) | 400 |
| $\Delta_{\text{prev}}^{\min}$ | PSC onset time minimum (previous trial effects) | -400 |
| $\Delta_{\text{prev}}^{\max}$ | PSC onset time maximum (previous trial effects) | 159 |
| $\Delta_{\text{next}}^{\min}$ | PSC onset time minimum (next trial effects) | 400 |
| $\Delta_{\text{next}}^{\max}$ | PSC onset time maximum (next trial effects) | 899 |
| $\sigma_{\text{scale}}$ | GP variance | 0.045 |
| $\ell_{\text{gp}}$ | GP lengthscale | 45 |
| $\sigma_{\text{noise,min}}^2$ | Noise variance minimum | 0.001 |
| $\sigma_{\text{noise,max}}^2$ | Noise variance maximum | 0.02 |

**Table S3:** Table of default parameters for simulation studies.

| Symbol | Interpretation | Default value |
| --- | --- | --- |
| $\phi_{\min}^0$ | Presynaptic sigmoid coefficient (0) minimum | 0.2 |
| $\phi_{\max}^0$ | Presynaptic sigmoid coefficient (0) maximum | 0.25 |
| $\phi_{\min}^1$ | Presynaptic sigmoid coefficient (1) minimum | 10 |
| $\phi_{\max}^1$ | Presynaptic sigmoid coefficient (1) maximum | 15 |
| $\tau_{r,\min}$ | Rise time constant minimum | 10 |
| $\tau_{r,\max}$ | Rise time constant maximum | 40 |
| $\tau_{\Delta,\min}$ | Difference between rise and decay time constants minimum | 250 |
| $\tau_{\Delta,\max}$ | Difference between rise and decay time constants maximum | 300 |
| $\alpha$ | Rate parameter for gamma-distributed spike times | $10^4$ |
| $\beta$ | Shape parameter for gamma-distributed spike times | 15 |
| $\Delta_{\min}$ | Minimum spike time | 60 |
| $w_{\min}^{\text{strong}}$ | Strong synaptic weight minimum | 20 |
| $w_{\max}^{\text{strong}}$ | Strong synaptic weight maximum | 40 |
| $w_{\text{mean}}^{\text{weak}}$ | Weak synaptic weight mean (of the unshifted distribution) | 4 |
| $w_{\min}^{\text{weak}}$ | Weak synaptic weight minimum | 5 |
| $\ell_{\text{gp}}$ | GP lengthscale | 50 |
| $\sigma_{\text{scale}}$ | GP noise variance | $4^{-3}$ |

**Figure S1:**
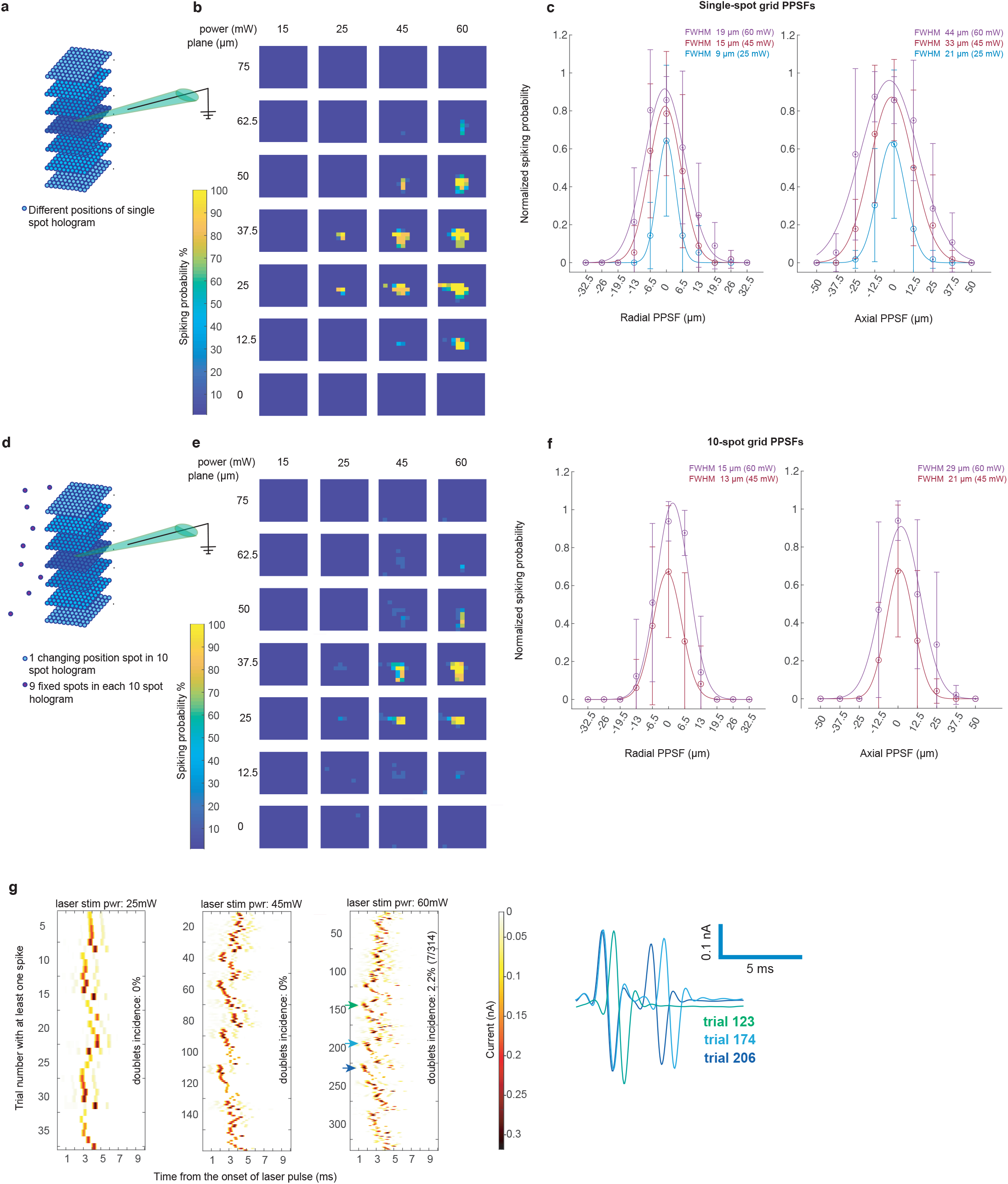
Physiological point spread functions (PPSFs) for holographic single-target and ensemble stimulation. **a**, Illustration of multi-plane grid of stimulation sites used to obtain single-target PPSFs. A loose-patched, opsin-positive cell was positioned to be in the center plane of the grid (37.5 *µ*m). Planes separated by 12.5 *µ*m. **b**, Single-cell example of mean spike probability when stimulating individual points on grid (7 repetitions per power). **c**, Mean radial (left panel) and axial (right panel) PPSF for a population of opsin-expressing PV cells. Error bars indicate standard deviation over n=8 cells. **d**, Illustration of multi-plane grid of stimulation sites used to obtain 10-target ensemble PPSFs. In these experiments, one point on the grid is targeted by the hologram while the remaining 9 “control” targets are outside of the grid area and kept fixed (i.e., do not change from stimulus to stimulus). **e**, Single-cell example of mean spike probability when stimulating using a 10-target hologram (with one target being the opsin-positive cell; 7 repetitions per power). Panels (a) and (c) display results from the same cell. **f**, Mean radial (left panel) and axial (right panel) PPSF for a population of opsin-positive PV cells when stimulating 10 targets at once. Error bars indicate standard deviation over n=7 cells. **g**, Incidence of multiple action potentials (i.e., doublets) in response to a single laser pulse. Trials shown are those where the patched cell spiked at least once. Postsynaptic neuron same as in panel (b). 10 ms time windows from the start of a 3 ms laser pulse are displayed. Exemplary traces displayed on the right are marked by arrows.

**Figure S2:**
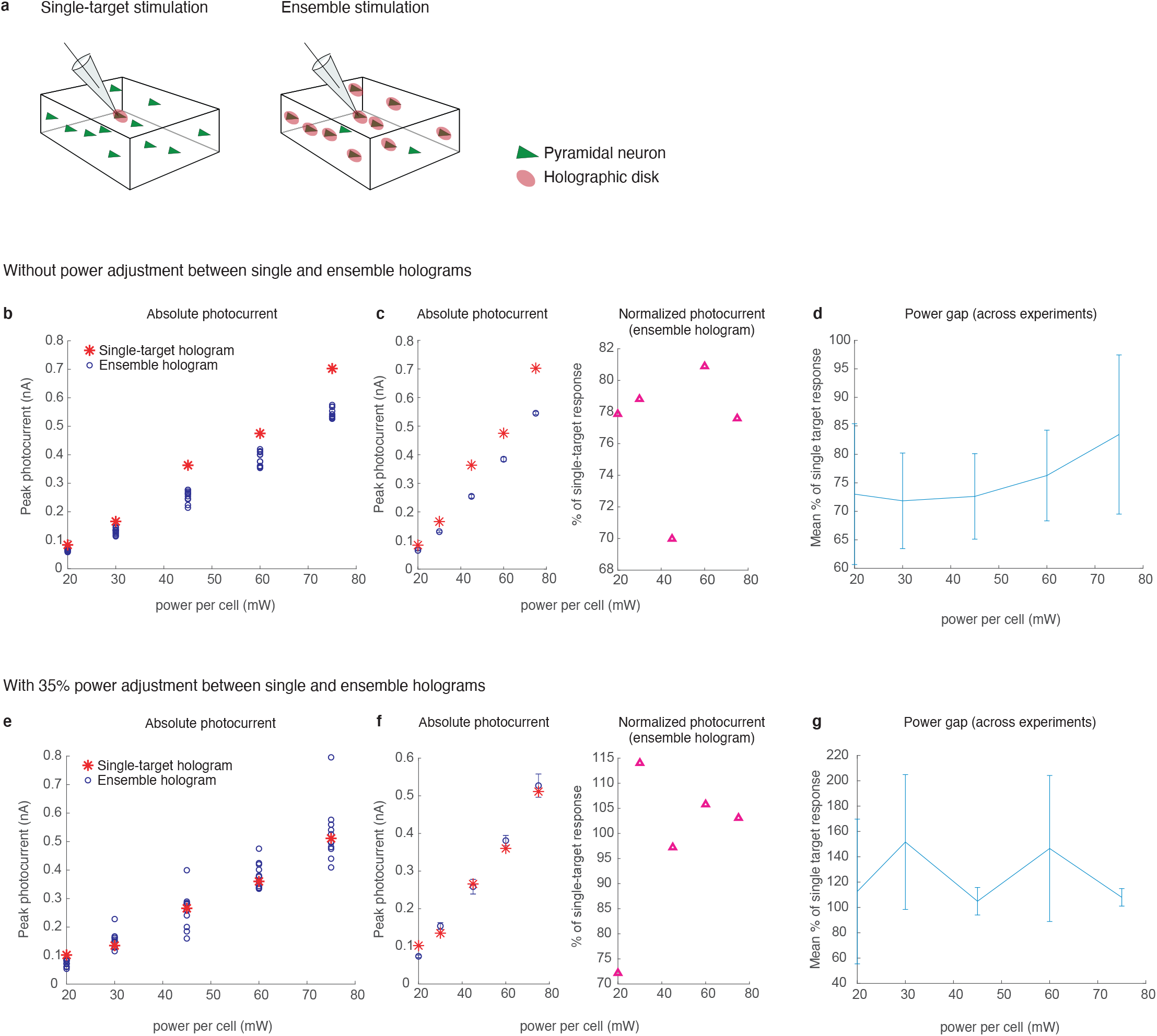
Resolving the “power gap” between single-target and ensemble holograms. **a**, Illustration of the two compared stimulation conditions. An opsin-positive cell was patched and photocurrents were recorded in whole-cell configuration while the regular connectivity mapping protocol was used. Trials when single-target holograms were targeting the patched cell (left) were compared with trials when the patched cell was a part of a 10-target ensemble hologram (right). **b**, Photocurrents from an example experiment. Red stars indicate the photocurrent amplitude across increasing stimulation power for a single-target hologram applied to the patched cell, blue circles represent the response amplitude across power for different ensemble holograms containing the patched cell. **c**, Same data as in b, but showing the mean *±* s.e.m. for the different ensemble hologram sets (left), and normalized to the corresponding single-target photocurrents (right, magenta triangles). **d**, Mean photocurrents evoked by ensemble holograms, normalized to photocurrents evoked by single-target holograms at the corresponding laser powers. Error bars show one standard deviation. Data collected across a population of cells; n=6 experiments performed on 1 pyramidal cell (Emx-Cre), 1 PV cell (PV-Cre), 4 SST cells (SST-Cre). **e-g**, same as b-d but with laser power for ensemble holograms increased by 35%. For g, data collected across n=2 experiments (1 pyramidal cell, 1 SST cell).

**Figure S3:**
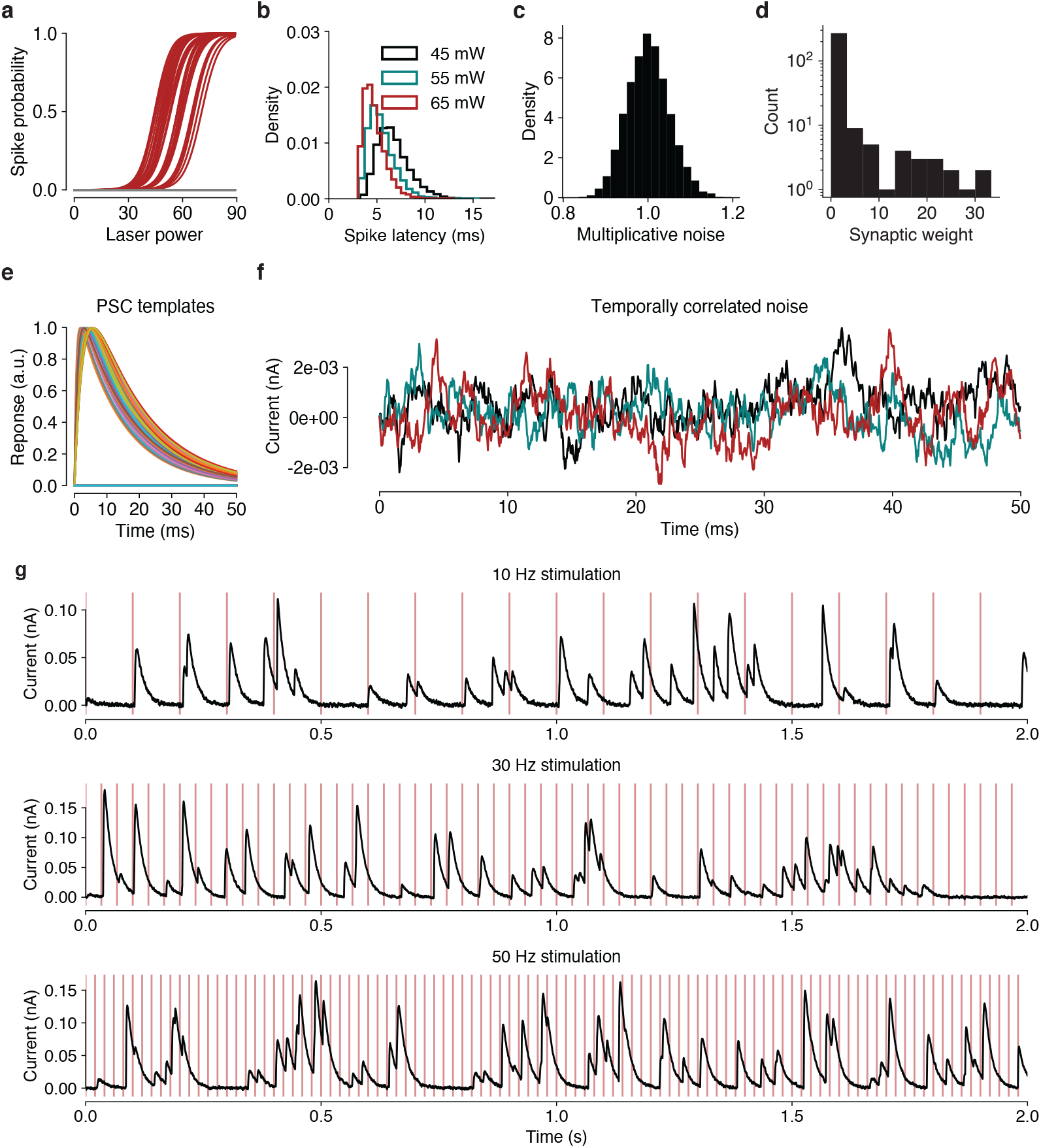
Components of an example simulated connectivity mapping experiment. **a**, Presynaptic spike probabilities as a function of laser power. Connected neurons shown in red, disconnected neurons shown in gray. **b**, Example histograms (shown as normalized densities) of presynaptic spike latencies for three typical laser powers. Average spike and transmission times decrease quadratically with increasing power. **c**, Example log-normal distribution of multiplicative noise terms that induce trial-to-trial variability in PSC amplitude. **d**, Distribution of synaptic weights, shown on log scale. **e**, Example PSC templates. Disconnected neuron templates represented by the zero vector. **f**, Three example noise processes sampled from a first-order autoregressive process. **g**, Example connectivity mapping experiment under increasing rates of stimulation. Simulation has 300 neurons with 10% connection probability and spontaneous synaptic currents occuring at 5 Hz.

**Figure S4:**
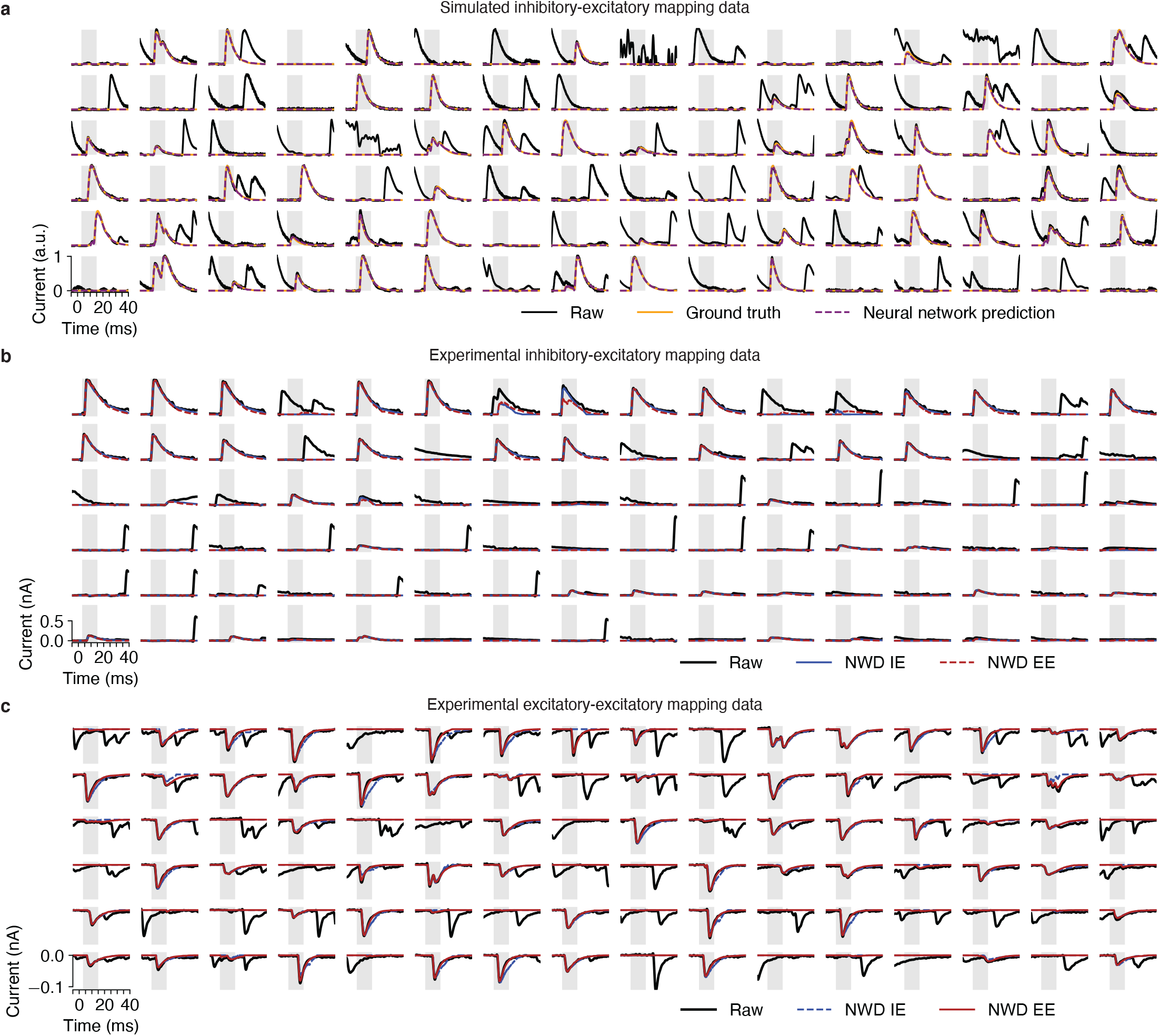
**a**, NWD network performance on simulated test data. Solid black line shows raw data given as input to the neural network. Orange line shows target trace. Dashed purple line shows prediction from the NWD network. **b**, Performance of NWD trained on simulated data matched to IE mapping experiments (solid, blue). For comparison, NWD trained on simulated data matched to EE mapping experiments shown as dashed red line. **c**, Performance of NWD trained on simulated data matched to EE mapping experiments (solid, red). For comparison, NWD trained on simulated data matched to IE mapping experiments shown as dashed blue line.

**Figure S5:**
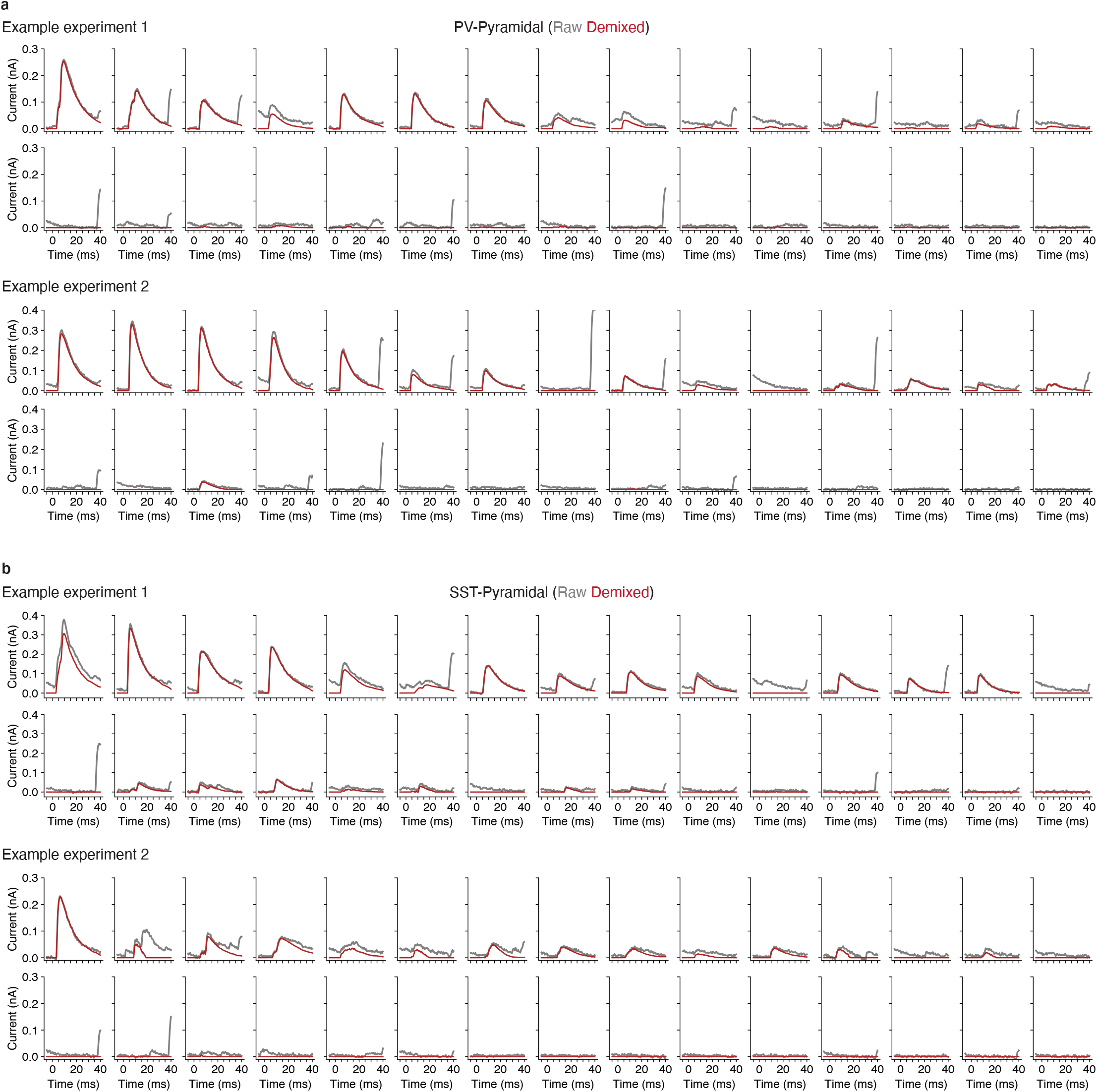
Example application of NWD to individual PSCs evoked by holographic ensemble stimulation in PV (a) and SST (b) to pyramidal mapping experiments. PSCs selected uniformly at random and sorted by magnitude of postsynaptic response.

**Figure S6:**
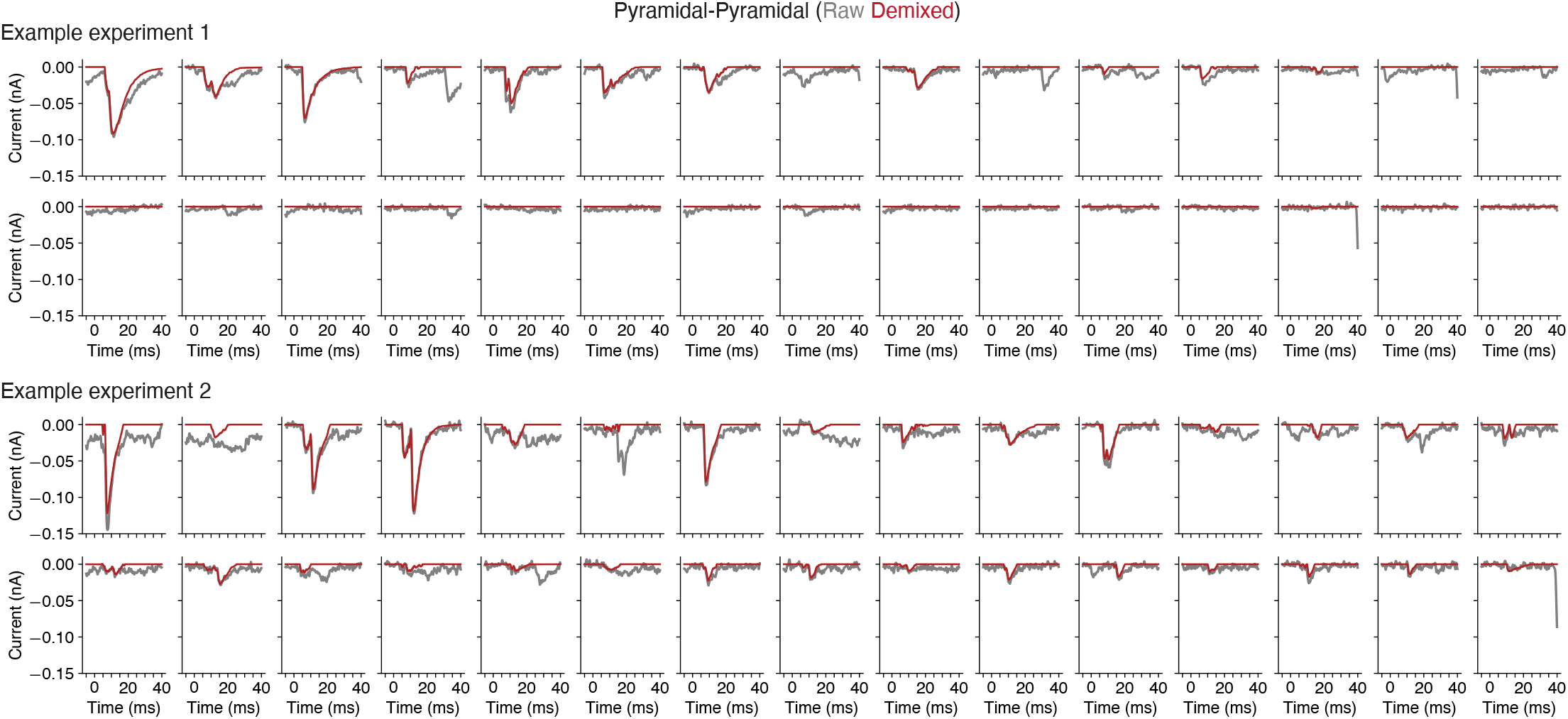
Example application of NWD to individual PSCs evoked by holographic ensemble stimulation in a pyramidal-pyramidal mapping experiment. PSCs selected uniformly at random and sorted by magnitude of postsynaptic response.

**Figure S7:**
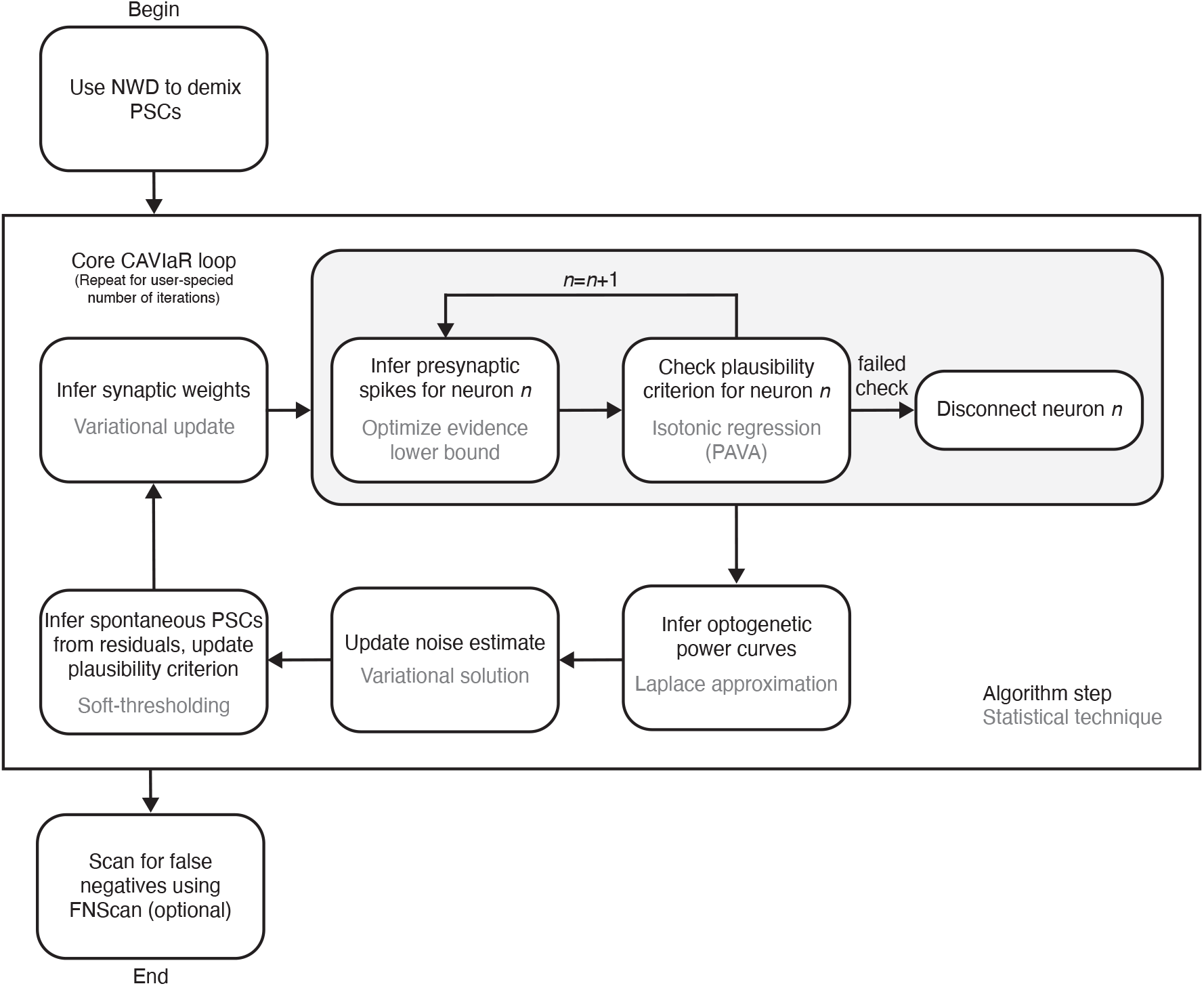
Flowchart detailing how CAVIaR operates (c.f. Algorithm 1).

**Figure S8:**
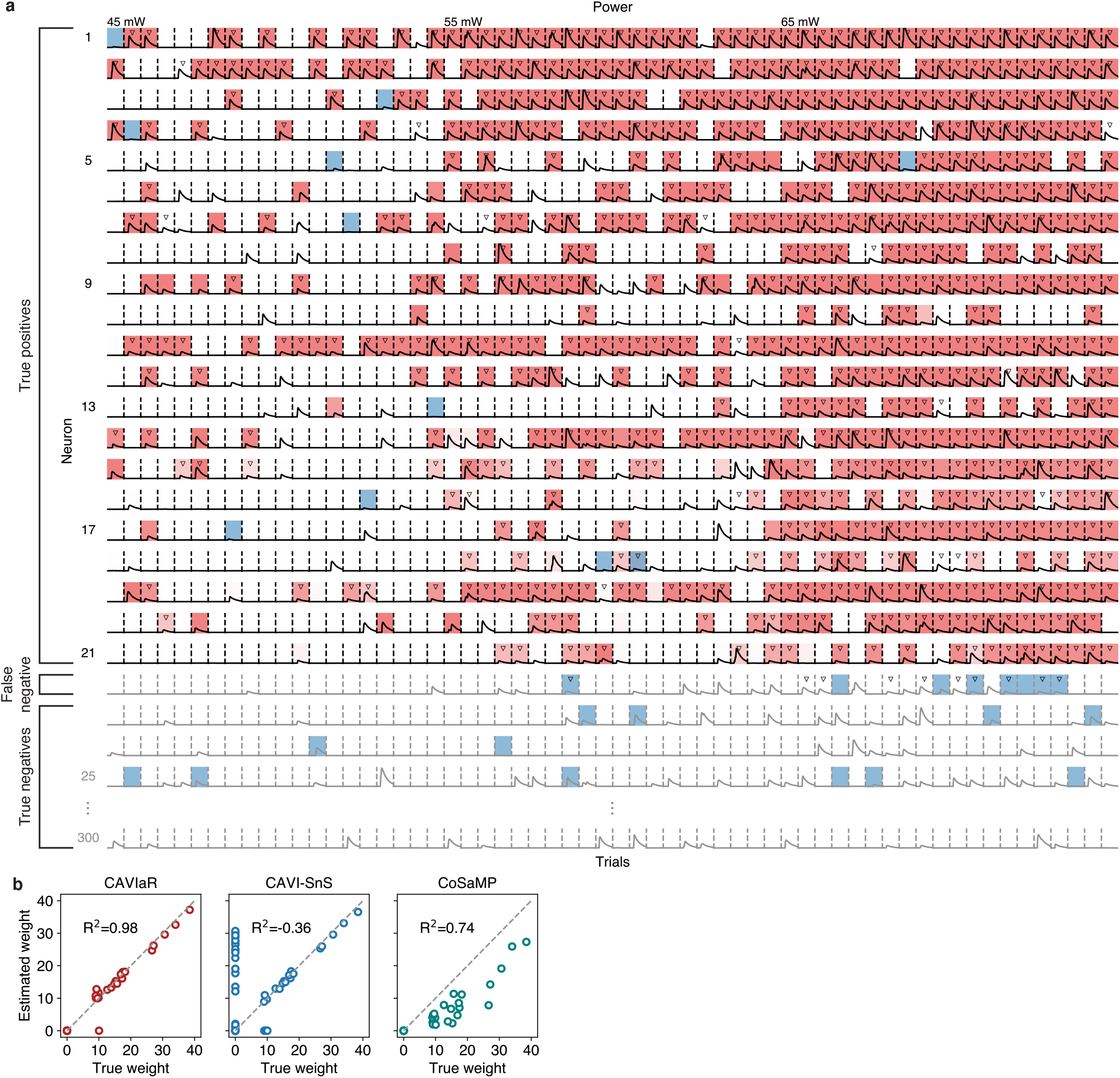
Large-scale classification of presynaptic spikes, spontaneous synaptic currents, and synaptic connectivity in a 3 minute simulated connectivity mapping experiment. *N* =300 neurons mapped using 10-target ensemble stimulation at 10 Hz. Spontaneous PSCs occur at a rate of 5 Hz. **a**, “Checkerboard” visualization of CAVIaR model inference. Each row shows sample of PSCs evoked by stimulating the listed neuron across multiple powers. In this example, 9 neurons in addition to the listed neuron are stimulated on each trial. Shaded red cells indicate detected presynaptic spike for listed neuron. Shaded blue cells indicate detected spontaneous PSC. Triangles indicate ground-truth presynaptic spikes. Accuracy of inferred presynaptic spikes for this simulation, 94.3%. Traces shown in gray indicate neurons that the model declared disconnected. Out of 300 neurons mapped, one putative connection was a false negative (neuron 22) due to CAVIaR incorrectly assigning its PSCs to other stimulated neurons or to spontaneous activity. **b**, Comparison of connectivity inference accuracy between CAVIaR, CAVI-SnS, and CoSaMP.

**Figure S9:**
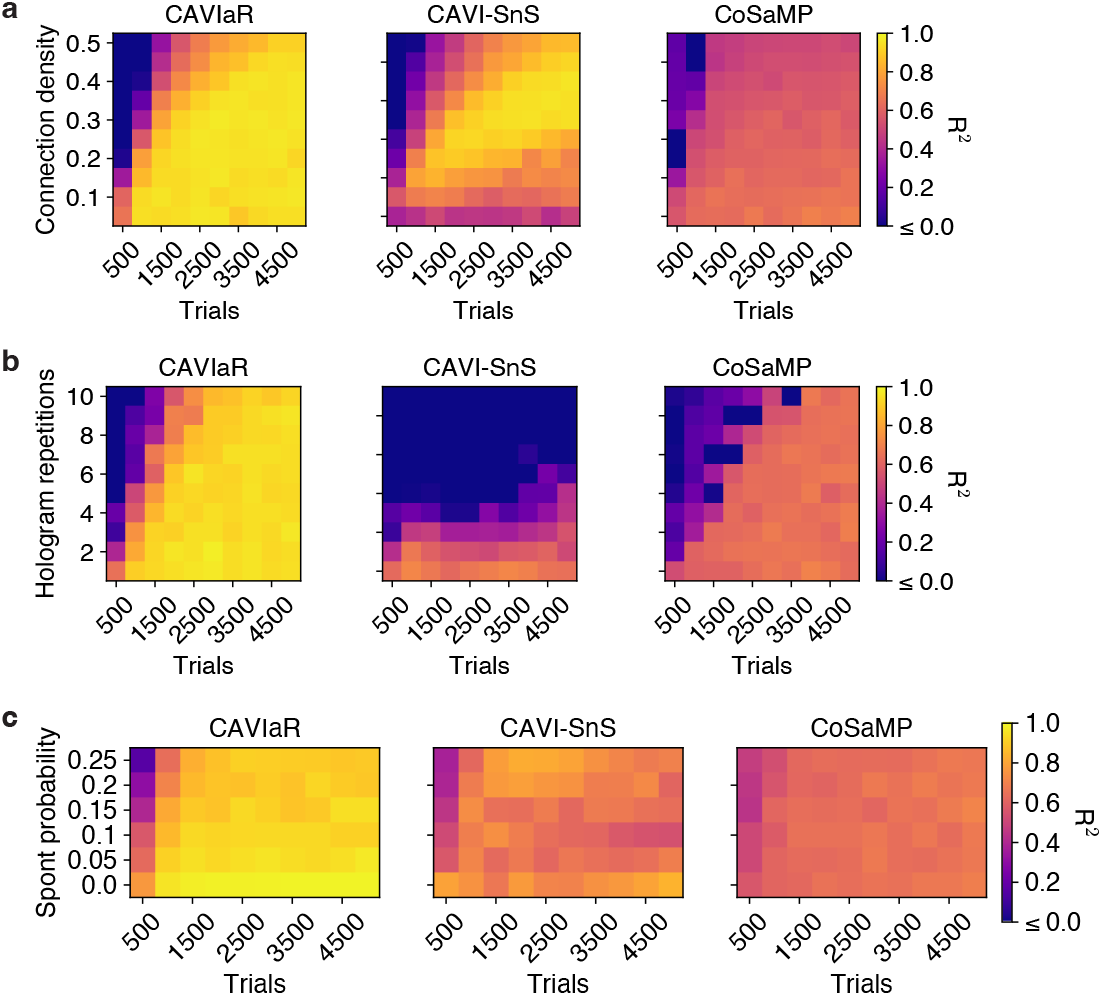
Performance and phase transitions for three connectivity inference techniques for a population of 300 candidate presynaptic neurons. **a**, Performance as a function of underlying density of synaptic connectivity (connection probability). Same as Figure 3b. **b**, Performance as a function of stimulus diversity (hologram repetitions). **c**, Performance as a function of the number of simultaneously targeted neurons. **d**, Performance as a function of probability of spontaneous PSC lying within admissible PSC initiation window. Fraction of connected neurons in b-d, 0.1. Note that in order to illustrate the behaviour of the three algorithms, these simulations involve varying only one parameter at a time while the others are kept fixed. However, the precise shape of these heatmaps could change if parameters are varied jointly. Default parameters: connection density, 0.1; hologram repetitions, 1; spontaneous PSC probability, 0.05.

**Figure S10:**
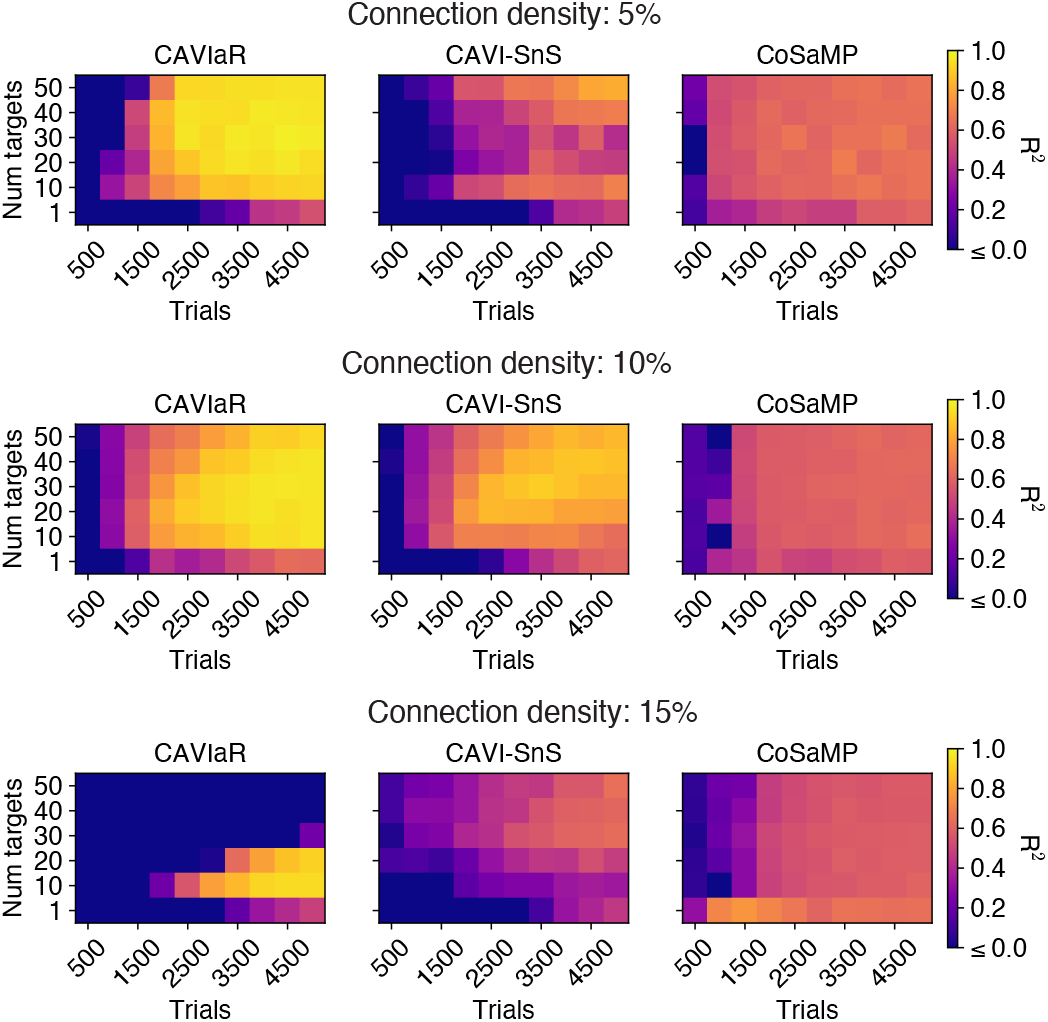
The optimal number of targets for CAVIaR, CAVI-SnS, and CoSaMP depends on the density of synaptic connectivity. Number of presynaptic candidate neurons, 1000; spontaneous PSC probability, 0.05; number of hologram repetitions, 1. When mapping populations with a 10% connection density, 20-target stimulation is optimal. However, at 5% connection density this increases to 30-40 targets, and at 15% connection density this reduces to 10 targets.

**Figure S11:**
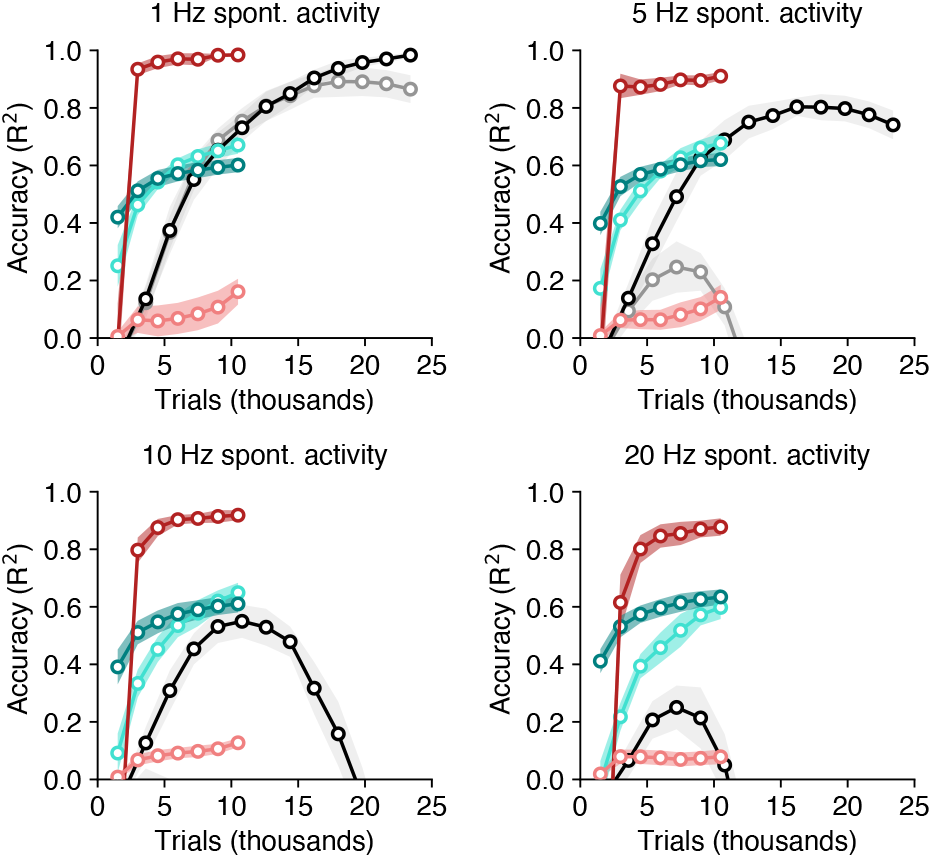
Convergence rates of each method as function of stimulation trials. C.f. Figure 3c. While mapping as a function of stimulation trials neglects the critical speedup enabled by NWD, CAVIaR nevertheless shows a substantial improvement in mapping efficiency.

**Figure S12:**
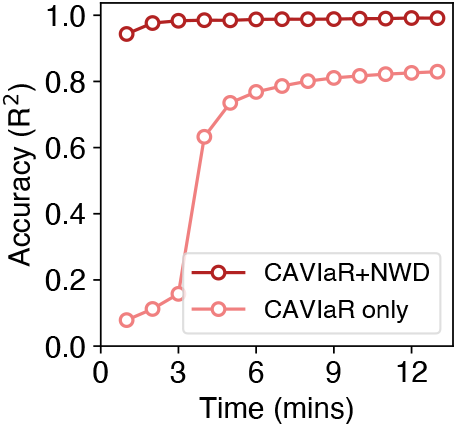
Performance of CAVIaR without NWD for longer experiment time shows eventual performance improvement, but does not match performance with NWD. Each data-point is an average of 10 simulations.

**Figure S13:**
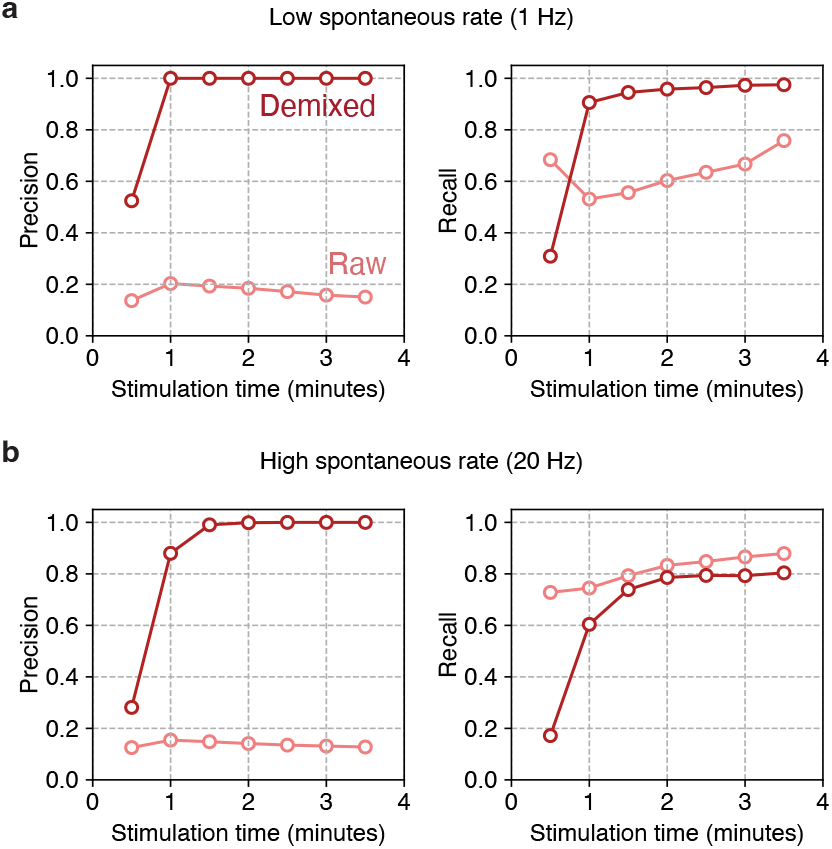
Convergence of precision and recall using CAVIaR in simulations. **a**, Performance of CAVIaR with low rates of spontaneous PSCs (1 Hz). **b**, Same as a, but with a 20 Hz rate of spontaneous PSCs. Population size, 1000; connection density, 10%; number of simultaneously stimulated targets, 20; stimulation frequency, 50 Hz. Dark line represents performance of CAVIaR with NWD; light line, without NWD. Note that recall is higher in b without NWD due to an excessive number of neurons being declared connected (c.f. precision *<* 0.2 for CAVIaR without NWD). Each data point is an average over 10 simulations.

**Figure S14:**
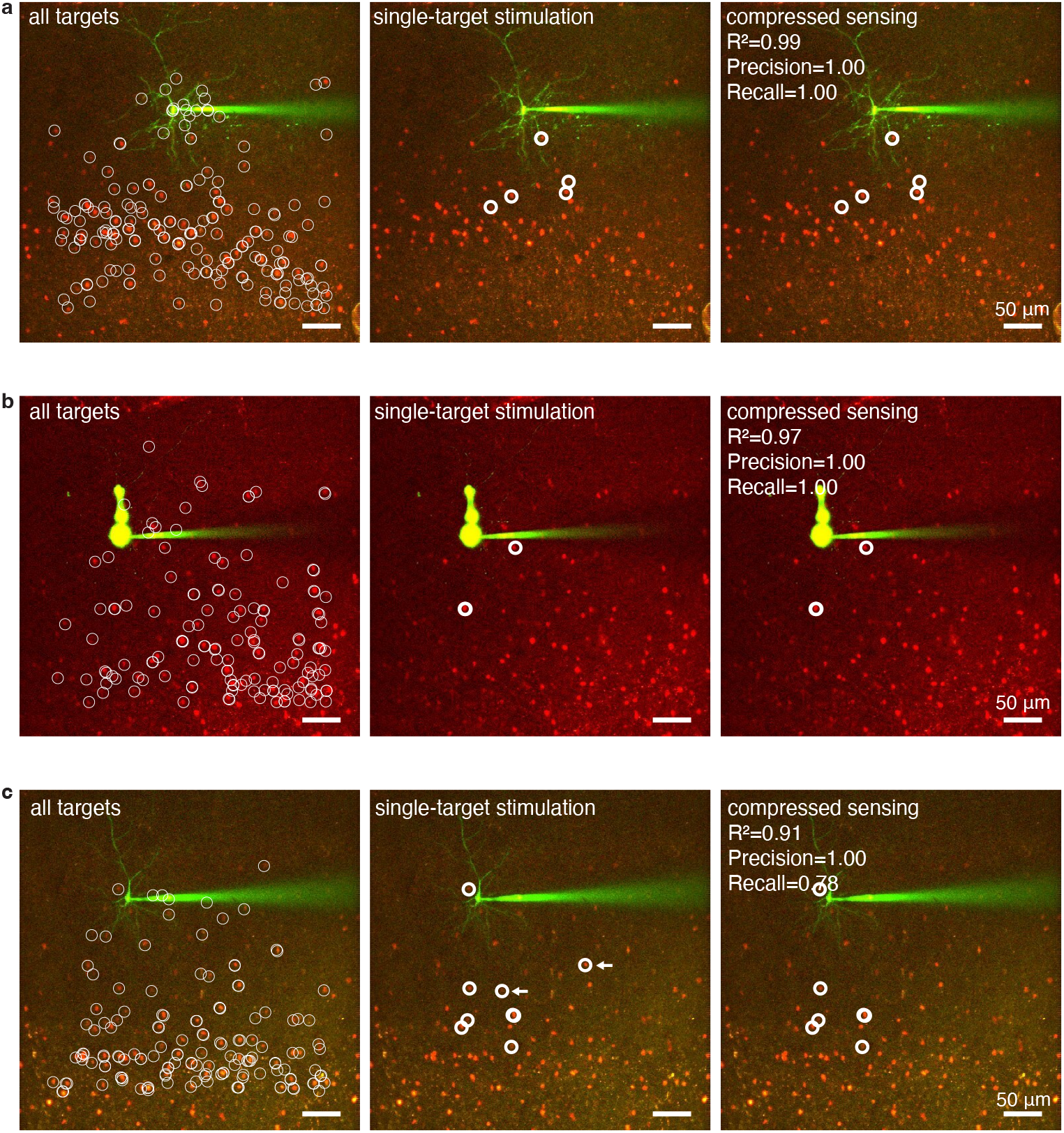
Additional comparisons between connectivity maps obtained using single-target and ensemble stimulation. Arrows in panel c show connections found using single-target stimulation that were not identified using ensemble stimulation.

**Figure S15:**
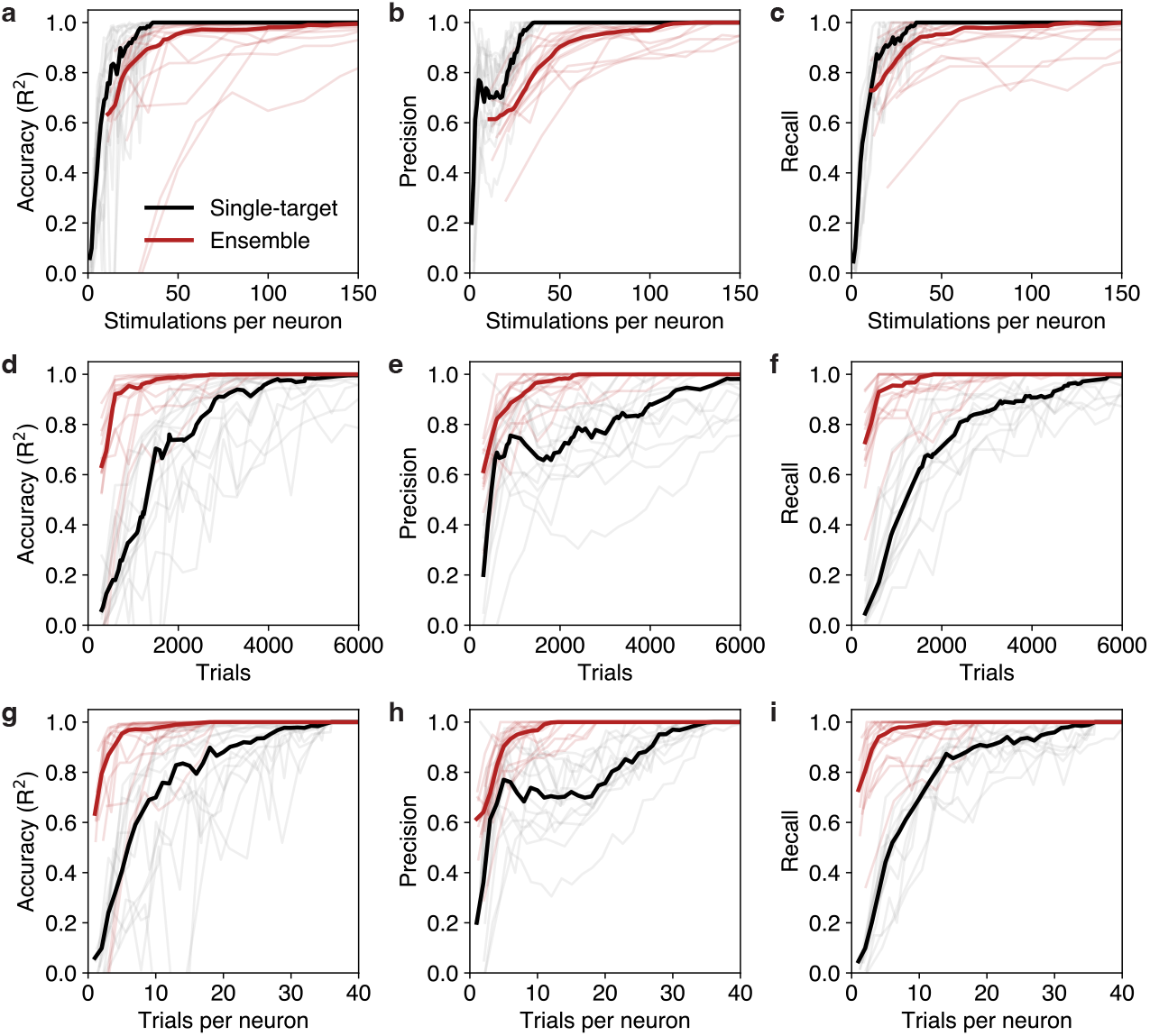
Analysis of convergence speed using single-target stimulation (black lines) and compressed sensing (red lines). **a-c**, Convergence of accuracy metrics (R^2^, precision, recall) as a function of number of times any given neuron is stimulated. **d-f**, Same as a-c, but for number of stimulation trials. **g-i**, same as d-f, but where trial counts have been normalized by population size (due to each experiment mapping different numbers of neurons). Dark lines show medians over 14 PV-pyramidal mapping experiments, faint lines show individual experiments. Note that performance metrics are defined with respect to the final estimates of each method, and therefore necessarily converge to 1. Hence “convergence” indicates “convergence to each method’s final estimate of connectivity”.

**Figure S16:**
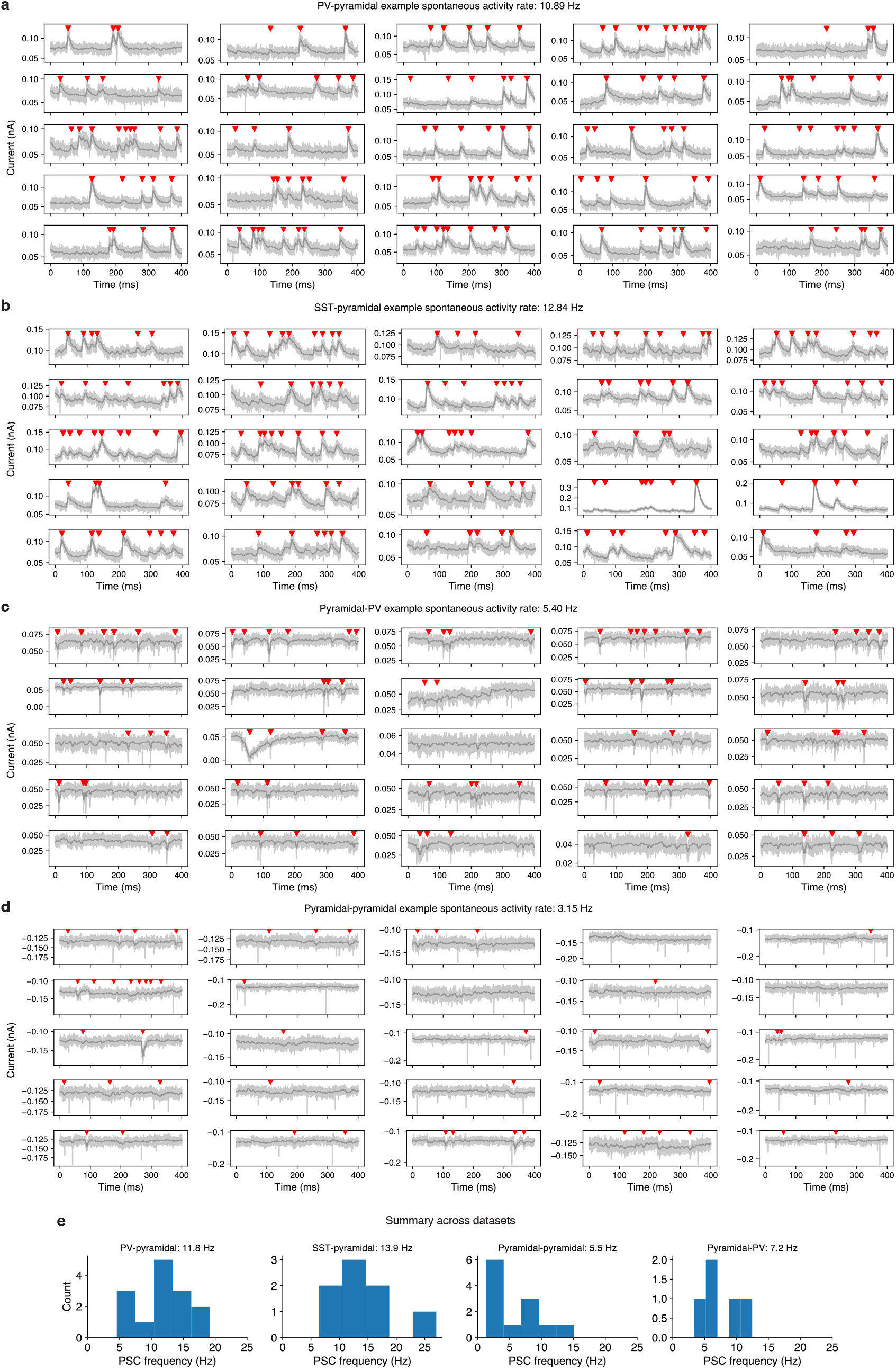
Determination of spontaneous PSC rates across all cell-type combinations used in this study. **a-d**, Example detected spontaneous PSCs (red triangles) from 400 ms windows of intracellular recordings without stimulation. Panels show only a subset of trials without stimulation to facilitate visualization. **e**, Distribution of spontaneous PSC frequencies across all datasets used in this study. Panel headings show corresponding averages. Note that different spontaneous PSC rates can arise due to the different holding potentials used for the patch-clamped neuron.

**Figure S17:**
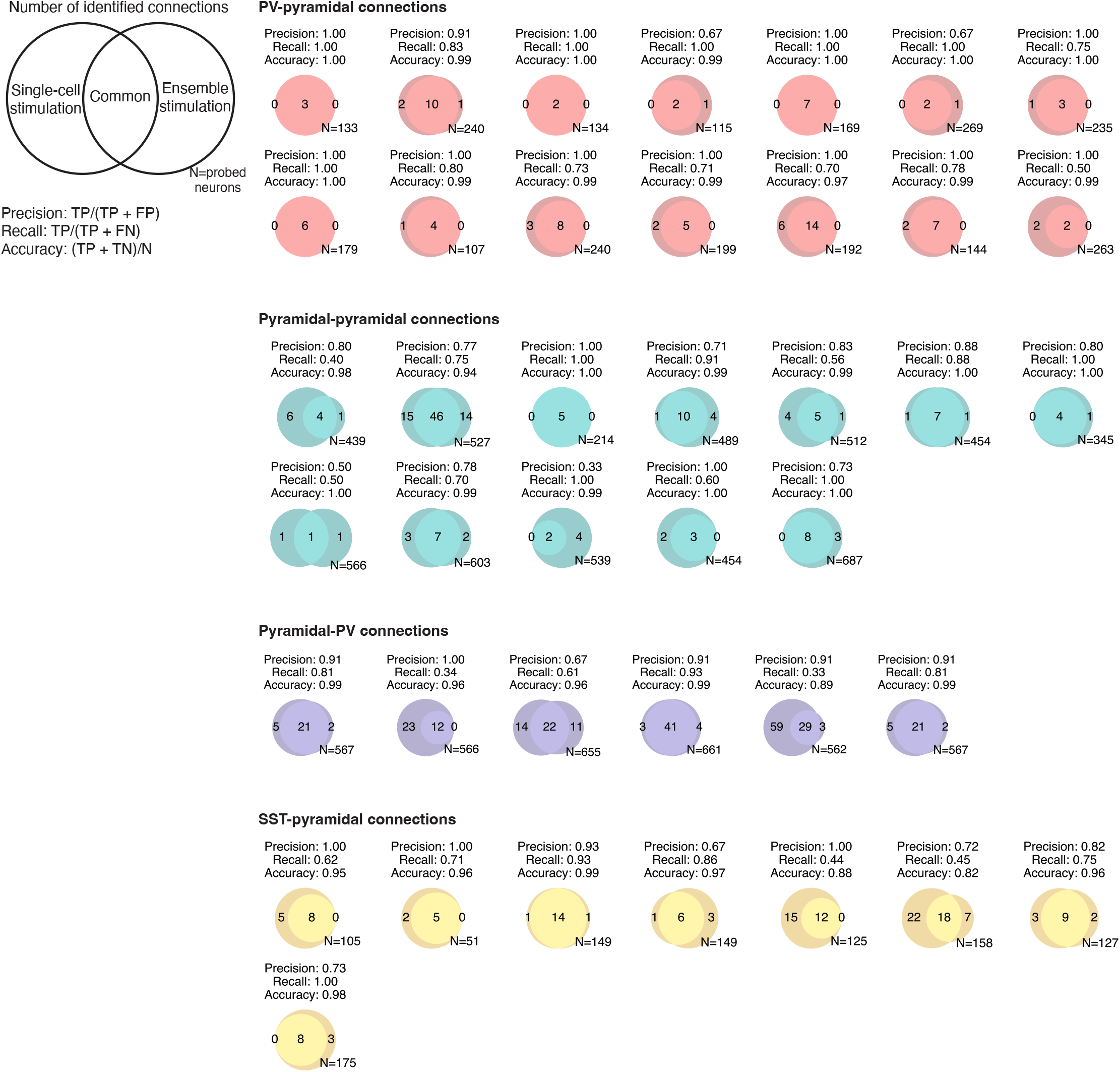
Comprehensive overview of CAVIaR performance for all mapping experiments used in this study. Top-left: The performance of CAVIaR is measured using the precision (fraction of connections identified using ensemble stimulation that are “true” connections), recall (fraction of “true” connections that are correctly identified using ensemble stimulation), and accuracy (total fraction of correctly classified connections). Here “true” connections are those identified using single-target stimulation, though note that single-target stimulation can also yield false-positives and false-negatives. TP: true positive, TN: true negative, FP: false positive, FN: false negative, N: total number of probed targets.

**Figure S18:**
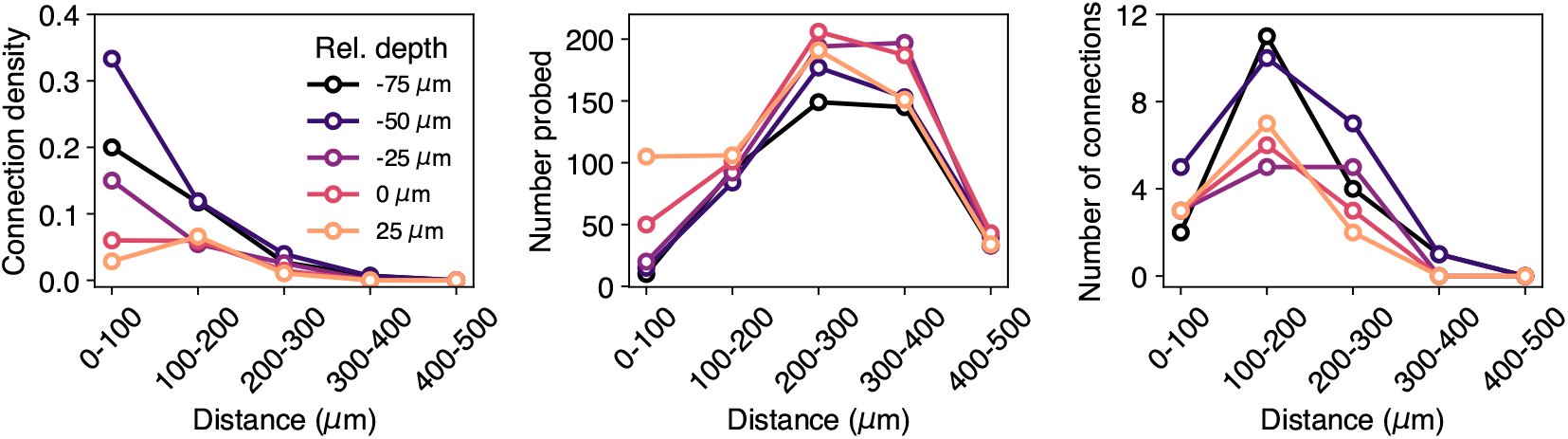
Connection density, number of probed targets, and number of identified connections across 14 PV-pyramidal mapping experiments. Each line represents the result of probing for connectivity among PV neurons at a different depth relative to the postsynaptic pyramidal neuron. Distance from the postsynaptic neuron is evaluated in two dimensions (c.f. Figure 5f). Depth is with respect to the mediolateral dimension. Pyramidal neurons were patched in L2/3, but could be at different depths within L2/3 themselves.

**Figure S19:**
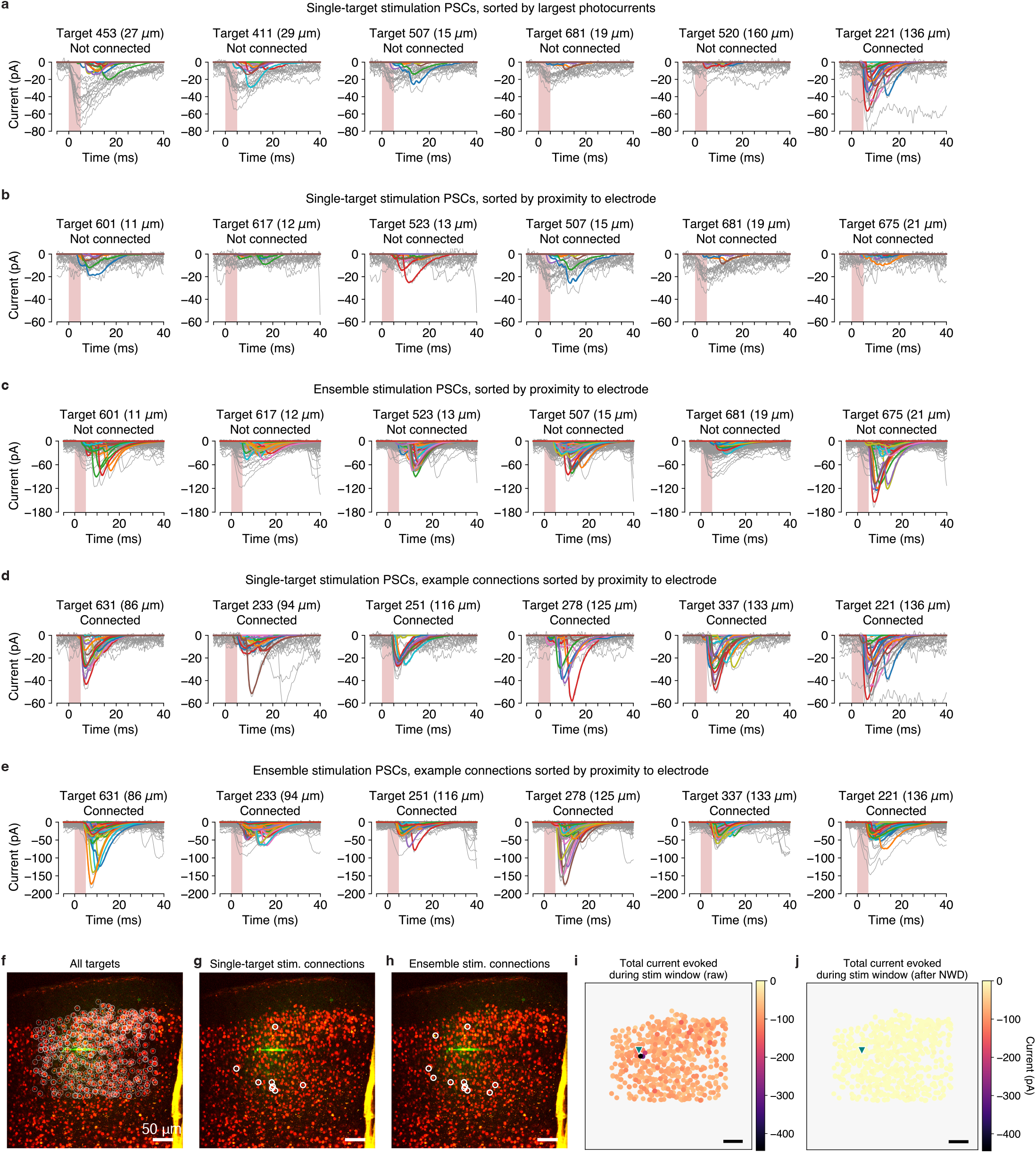
Analysis of the effect of NWD on direct photocurrent artifacts in pyramidal-pyramidal (Emx-Cre; AAV-st-ChroME2s-mRuby3) mapping experiments (corresponds to Figure 6a). **a-e**, Photocurrents and postsynaptic responses for multiple conditions (single-target stimulation, ensemble stimulation), connection statuses (connected, not connected), and orderings (sorted by largest photocurrents, proximity to electrode). Raw traces shown in gray, demixed traces shown in color. Numbers inside parentheses show distance from electrode tip. See Figure S20 for single-trial comparison between raw and demixed current traces. **f-h**, Stimulation FOVs showing all targets (f), connections inferred using single-target stimulation (g), and connections inferred using ensemble stimulation (h). **i, j**, Spatial representation of the magnitude of evoked currents during stimulation period before (i) and after (j) applying NWD. Magnitude given as the total summed current (in picoamperes) during the stimulation window (shaded red regions in a-e) across all stimulation trials and powers. Heightened currents near the electrode tip (base of green triangle) correspond to direct photocurrents on the patched cell.

**Figure S20:**
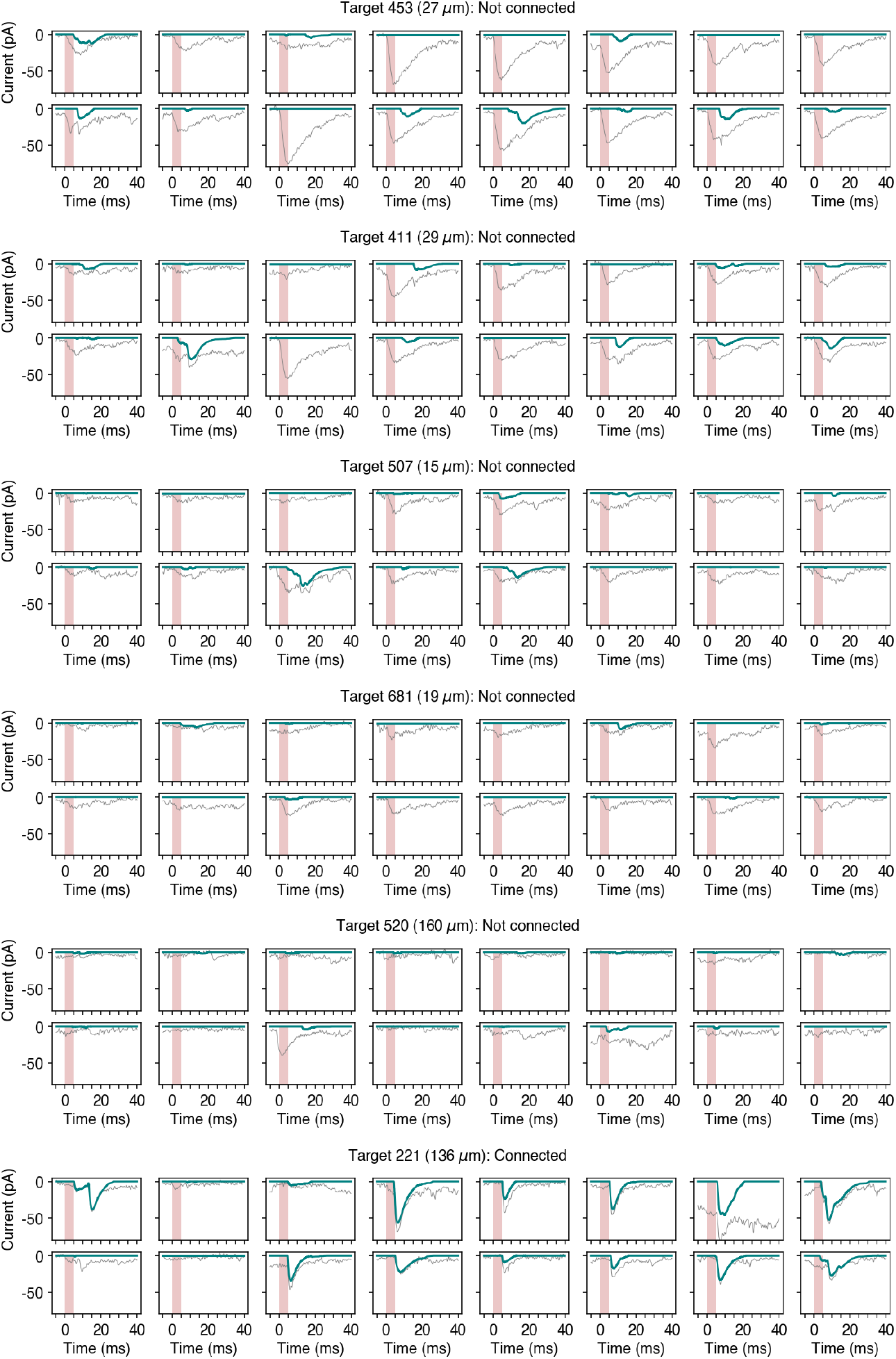
Comparison between raw (gray) and demixed (teal) current traces in the presence of photocurrent artifacts. Traces correspond to all shown targets in Figure S19a (i.e. PSCs evoked by single-target stimulation, ordered by largest photocurrents).

**Figure S21:**
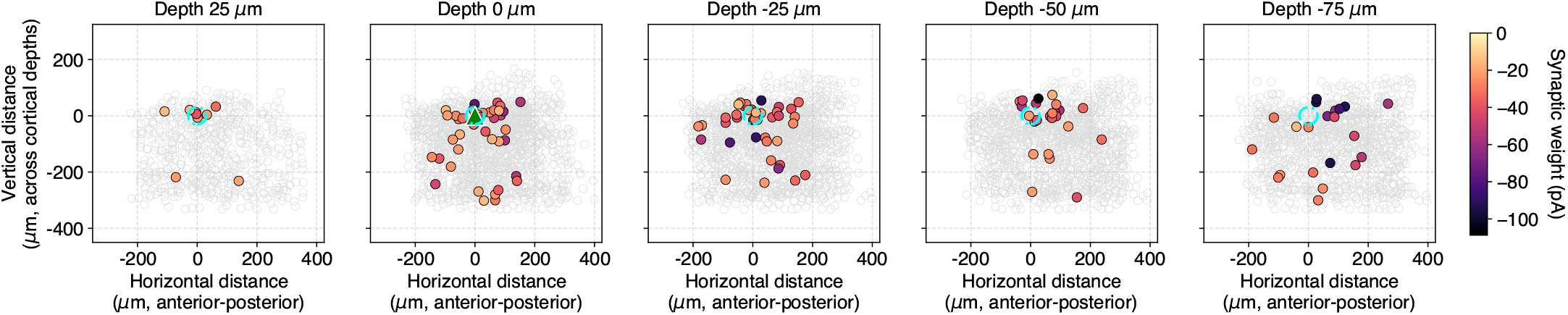
Pyramidal-pyramidal connections across 12 experiments, split by plane. Cyan circle denotes region with 30 *µ*m radius where photocurrents are most likely.

**Figure S22:**
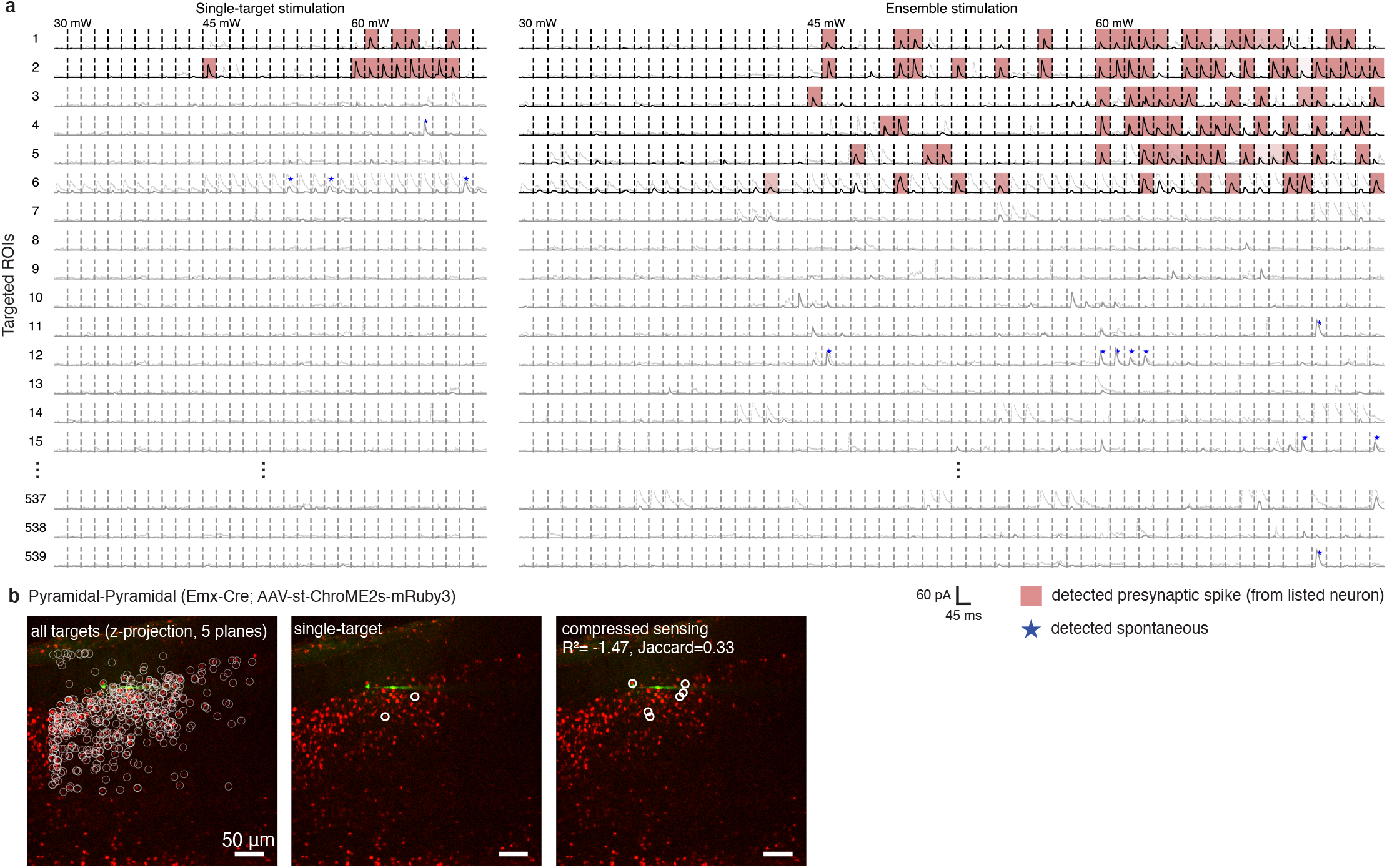
Outlier experiment corresponding to marked datapoint in Figure 6d. **a**, Checkerboard visualization of synaptic currents and photocurrents evoked by single-target (left) and holographic ensemble (right) stimulation. Solid gray traces correspond to demixed PSCs from ROIs considered unconnected; dashed faint gray traces are the same but without demixing. Note the presence of photocurrents in row 6 largely suppressed by NWD. Note that the sign of the excitatory currents are reversed for ease of visualization. **b**, Z-projected stimulation FOV (left), and comparison between connections identified using single-target stimulation (middle) and holographic ensemble stimulation (right). Two factors could contribute to this outlier experiment. The first is photocurrent contamination: in some cases NWD does not perfectly subtract photocurrents arising from direct stimulation in pyramidal-pyramidal mapping experiments. Thus, residual photocurrents could lead to false positive connections (in this case when mapping using ensemble stimulation specifically). The second is potential differences in the amount of laser power ultimately delivered to neurons during stimulation. Although we have taken considerable steps during calibration to ensure that the amount of power delivered per target is the same whether performing single-target or ensemble stimulation (Figure S2), we optimized this on average across experiments. Therefore, subtle differences in power delivery between single-target and ensemble stimulation can still arise in individual experiments, potentially leading to the observed discrepancy in this experiment even if CAVIaR makes the correct inference.

**Figure S23:**
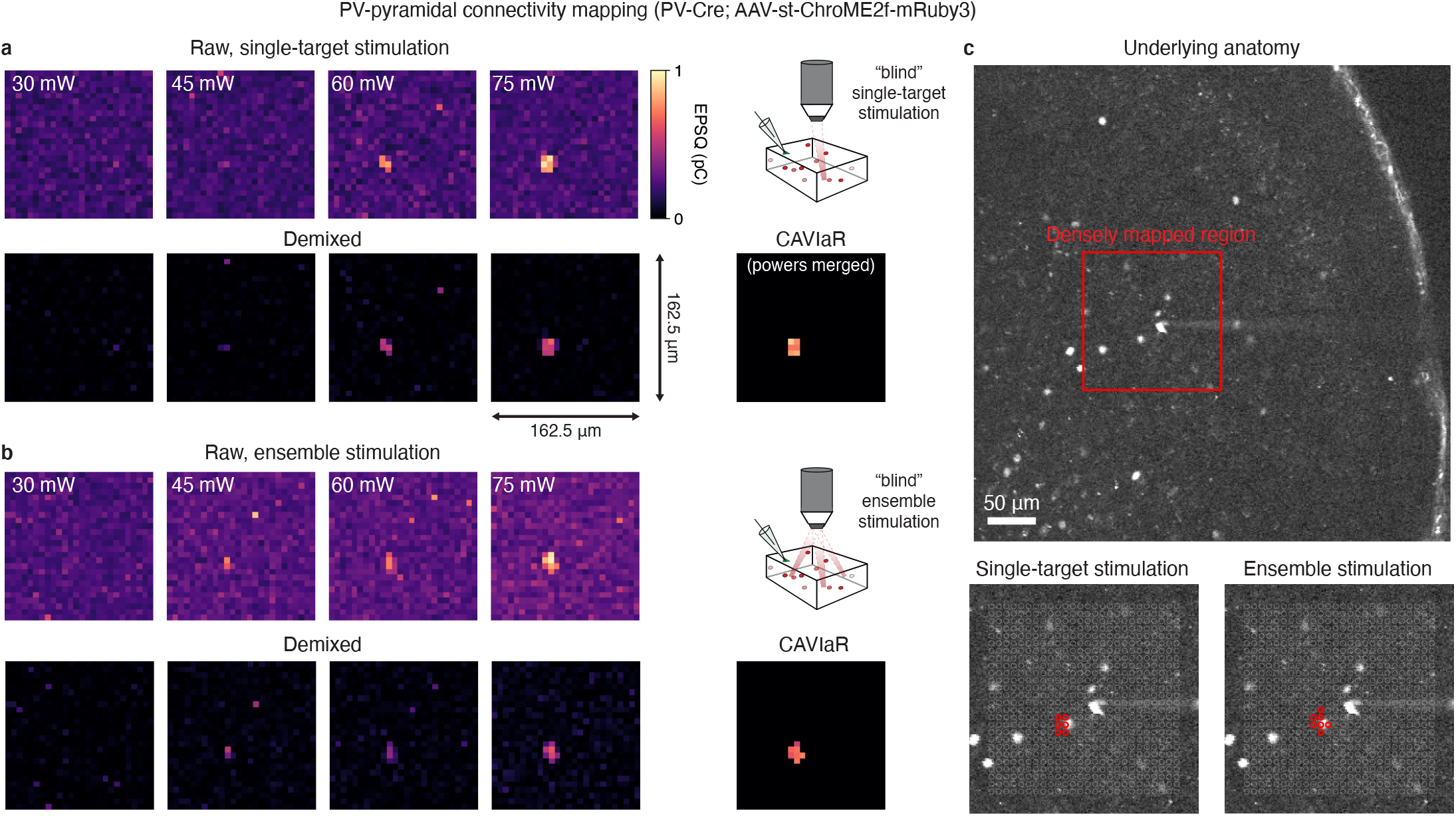
Demixing and connectivity inference for “blind” grid mapping experiments. **a**, Grid mapping of PV-pyramidal connectivity using single-target stimulation over four powers (30-75 mW). One out of five planes are shown as an example. Note that maps of synaptic connectivity obtained by CAVIaR “merge” the multi-power demixed maps into a single map. **b**, Grid mapping of same PV-pyramidal experiment using holographic ensemble stimulation (on five planes over 25 to -75 *µ*m, separated by 25 *µ*m each; only plane 0 *µ*m shown for comparison) over four powers. Agreement between the two maps validates the use of CAVIaR in this regime. Each pixel in the raw maps shown in (a) and (b) is obtained by averaging across all PSCs evoked by stimulation of an ensemble containing that pixel. **c**, Overlay of connected pixels (red circles) on underlying anatomy (obtained by imaging expression of mRuby) confirms that grid mapping using both single-target and ensemble stimulation correctly identifies an opsin-expressing neuron. Also note that this example experiment includes laser powers up to 75 mW (exceeding the range used to characterize PPSFs in Figure S1), which should be considered when interpreting the sizes of the connected regions.

**Figure S24:**
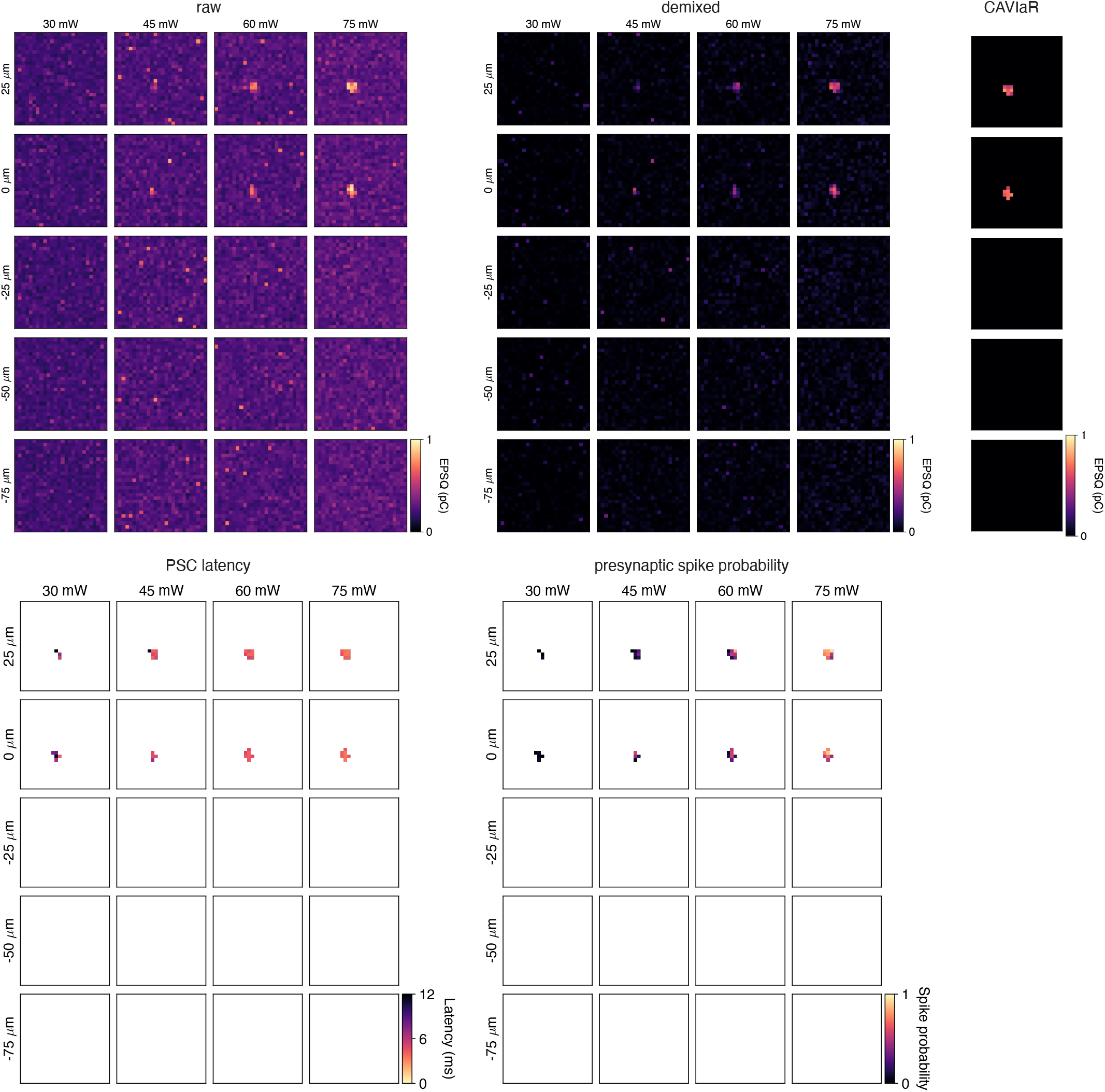
Demixing and denoising of optogenetic “blind” grid mapping data using NWD and CAVIaR. Example shows PV-pyramidal connectivity mapping (PV-Cre ChroME2f). Postsynaptic cell is on plane 0 *µ*m.

**Figure S25:**
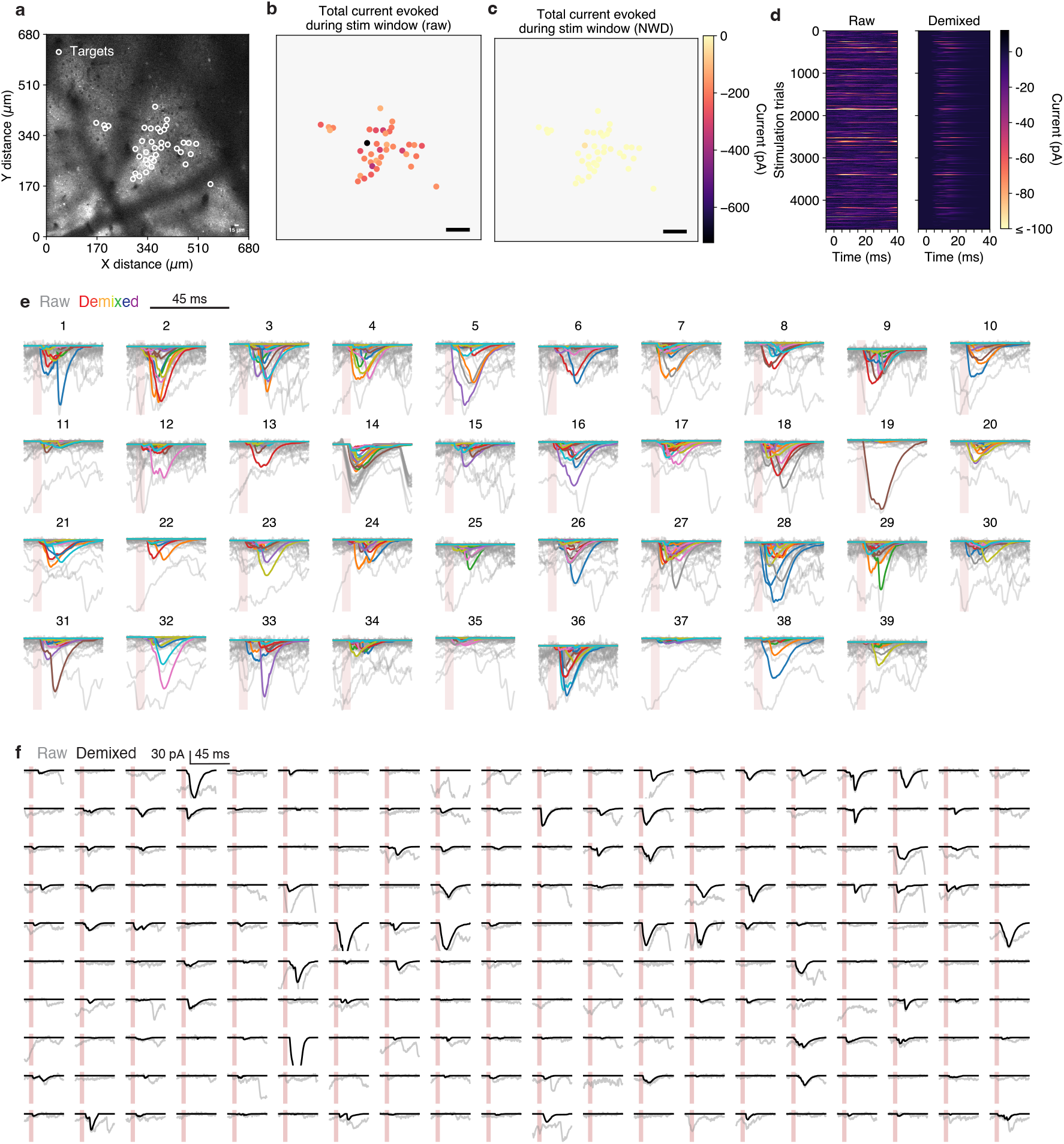
Application of NWD to an *in vivo* connectivity mapping experiment. **a**, Field of view for an experiment mapping pyramidal-to-SST connections. Pyramidal neurons and some SST neurons (including the patched postsynaptic neuron) express a GCaMP8m-ChroME2s fusion. White circles represent stimulation targets. **b**, Map of the total current evoked when stimulating at each location during the 5 ms stimulation window. Total current defined as the cumulative sum of the current measurement over time. **c**, Same as b, but after applying NWD to suppress photocurrents. **d**, Matrix visualization of the postsynaptic current evoked by stimulation before (left) and after (right) demixing. **e**, Comparison of raw (gray traces) and demixed (colored traces) synaptic currents evoked by stimulation. Each cluster of traces corresponds to the stimulation of a different target (indicated by number above traces). Traces normalized between 0 and 1 for each target. C.f. raw vs demixed EPSCs in Figure S19 for an analogous *in vitro* experiment. **f**, Comparison of raw (gray) and demixed (black) traces for individual trials. Shaded red bars correspond to 5 ms stimulation periods. NWD successfully isolates plausible EPSCs under *in vivo* conditions.

**Figure S26:**
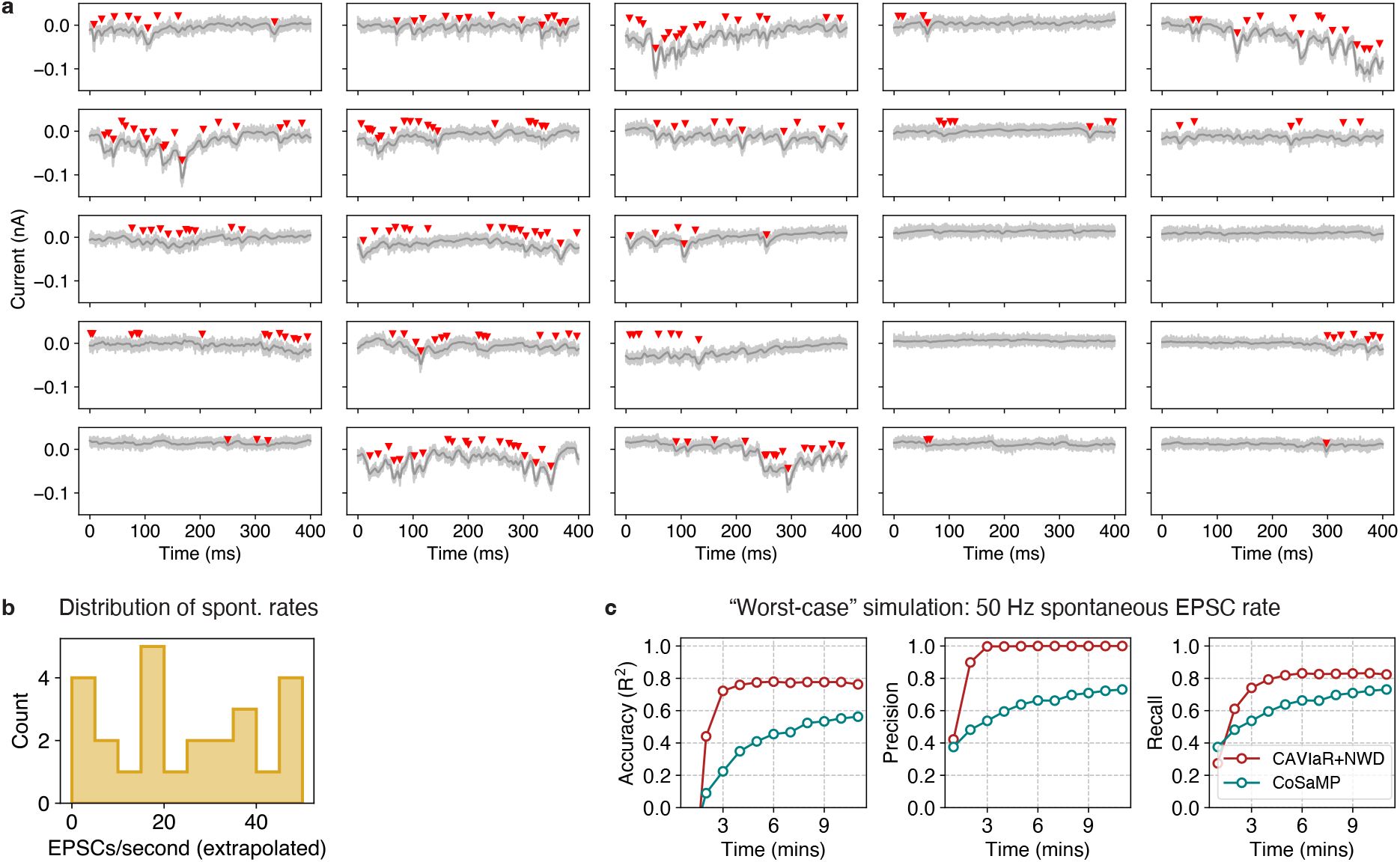
Analysis of CAVIaR performance under *in vivo*-like conditions. **a**, Detection of spontaneous EPSCs from an *in vivo* experiment during 25 periods without stimulation. Light gray traces show raw data, dark gray traces are low-pass filtered. Red triangles indicate putative EPSC peaks. **b**, Distribution of EPSC rates across 25 periods of *in vivo* spontaneous activity. Mean, 23.6 Hz. Max, 50 Hz. EPSC rates extrapolated from 400 ms windows. **c**, Performance of CAVIaR (with NWD) and CoSaMP with a 50 Hz rate of spontaneous PSCs, intended to reflect *in vivo* noise conditions. N=1000 neurons, 20-target stimulation, 50 Hz stimulation speed, 10% connection density.

### Supplementary note 1

In this note, we consider the probability of encountering a spurious connection where spontaneous PSCs arrive only when stimulating an unconnected neuron at the maximum laser power, but not at any of the lower powers. In particular, we show that the probability of a spurious connection only appearing at the highest laser power is vanishingly low.

Assume we stimulate a putative neuron 10 times per power at three different powers, and that the probability of a spontaneous PSC arriving within the stimulus response window is *p*. The probability that at least 3/10 trials show a spontaneous PSC is given by summing the binomial probability

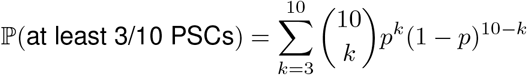

Thus, the probability of at least 3/10 spontaneous PSCs at the highest power and none at the lower two powers is

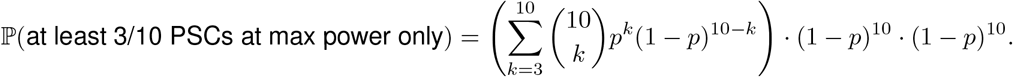

Plotting the above expression as a function of spontaneous PSC probability *p* yields Figure S27.

**Figure S27:**
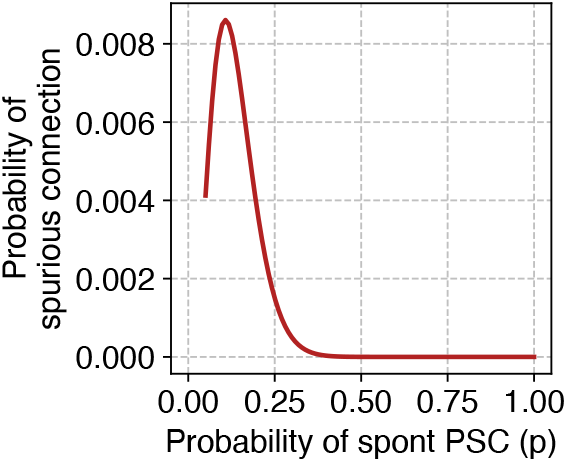
Probability of a spurious connection arising due to spontaneous PSCs at maximum power only.

From Figure S27 one can see that as the rate of spontaneous activity increases, it becomes increasingly unlikely for spontaneous PSCs to arrive only at the maximum power. The maximum value of the probability curve is just 0.86% (attained when the probability of a spontaneous PSC coinciding with a stimulation trial is 0.1). Hence this scenario is so rare as to not be worth consideration in practice.

### Supplementary note 2

In this note, we consider whether intrinsic plasticity could have occurred as a result of repeated ensemble stimulation. Note that, as we argue in the discussion, the typical rate of stimulation for any given neuron is low – less than 3 Hz in our experiments, which is too slow to induce intrinsic plasticity for neurons expressing the ChroME2 opsins [Sridharan et al., 2022]. Nevertheless, we sought to confirm that no such intrinsic plasticity effects were present in our data.

To precisely determine variables related to intrinsic plasticity (e.g. the spiking threshold and the current-to-spike relationship) in our experiments, one would need to know exactly when neurons are spiking in response to ensemble stimulation. Technically, this would require challenging experiments based on voltage imaging, which are beyond the scope of our study. Fortunately, CAVIaR performs inference of when neurons spike in response to stimulation, thereby providing a useful approximation.

Since we cannot directly access the input currents and action potentials for the neurons we map, we have approximated the spiking threshold and the current-to-spike relationship in the following way. First, the spike threshold was approximated as the minimal laser power required to spike a neuron with at least 0.25 probability. Second, the current-to-spike relationship was approximated by the “power-to-spike relationship” (i.e., spike probability as a function of laser power). In both cases, the spikes were inferred using CAVIaR.

**Figure S28:**
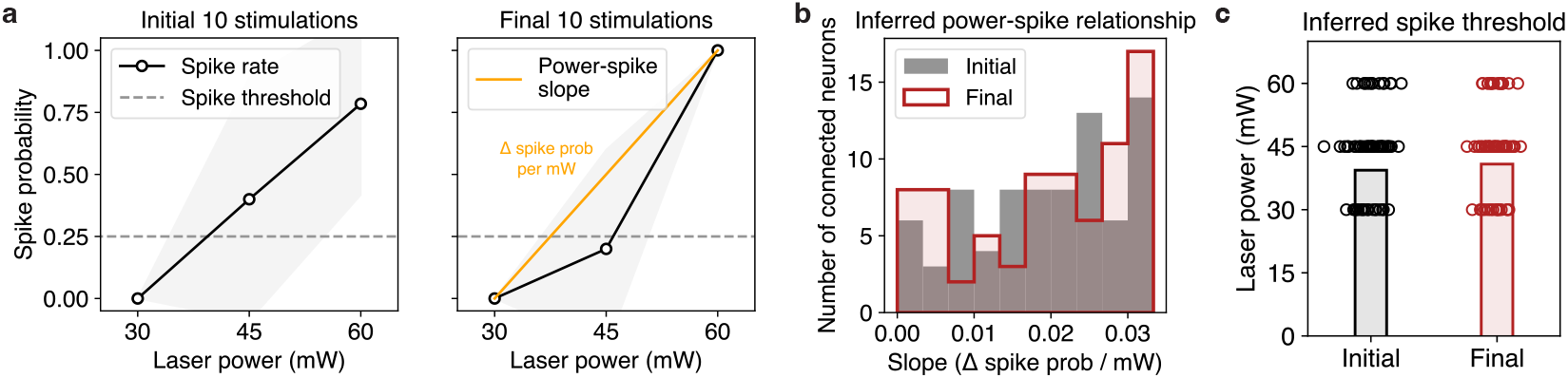
Analysis of potential changes in spike threshold or power-dependence as a result of repeated ensemble stimulation. **a**, Probability of spiking as a function of laser power using the initial 10 spikes (left) vs the final 10 spikes (right). Shaded error bars represent one standard deviation. Dashed horizontal line represents a spike probability of 0.25. The least power required to cross a spike probability of 0.25 is considered the “spike threshold” laser power. Orange line represents the “power-spike” slope; i.e. the change in spike probability per mW laser power. The power-spike slope is intended to approximate the slope of the current-to-spike relationship. **b**, No statistically significant difference between spike threshold before (initial) vs after (final) repeated ensemble stimulation (p=0.16, Wilcoxon signed-rank test). Shaded bars show means across all neurons and all experiments. **c**, No statistically significant difference between power-spike slope before vs after ensemble stimulation (p=0.57, Wilcoxon signed-rank test). All spikes are inferred from PSCs using CAVIaR. Analysis performed on all 78 detected presynaptic neurons across all PV-pyramidal experiments.

Next, to determine whether such variables changed with repeated stimulation, we estimated the power-spike relationship using both the initial 10 ensemble stimulation trials and the final 10 ensemble stimulation trials (Figure S28a) for every neuron across every experiment. This showed firstly that spike threshold did not significantly change over the course of the experiments (p=0.16, Wilcoxon signed-rank test; Figure S28b), and secondly that the slope of the power-spike relationship (i.e. the change in spike probability per mW laser power) also did not significantly change (p=0.57, Wilcoxon signed-rank test; Figure S28c). However, we acknowledge that these metrics rely on CAVIaR’s inferences (which are not necessarily perfect), and that with perfect knowledge of how each neuron spiked these results could change. Nevertheless, we believe these results represent the best we could achieve without performing a fundamentally new study based on voltage imaging.

We also verified that the input resistance of the postsynaptic neuron at the beginning and end of the experiment were not significantly different (mean input resistance at beginning of experiments, 105 MΩ; mean input resistance at end of experiments 109 MΩ; p-value of difference=0.85, independent t-test).

## Notes

https://github.com/marcustriplett/circuitmap

### Competing Interest Statement

The authors have declared no competing interest.

### Summary of Updates

Revised manuscript to include improved reporting metrics, further analysis of experimental data, additional simulations.

